# Activity dependent translation in astrocytes dynamically alters the proteome of the perisynaptic astrocyte process

**DOI:** 10.1101/2020.04.08.033027

**Authors:** D. Sapkota, M.S.J. Kater, K. Sakers, K.R. Nygaard, Y. Liu, A.M. Lake, R. Khazanchi, R.R. Khankan, A.B. Smit, S.E. Maloney, M.H.G. Verheijen, Y. Zhang, J.D. Dougherty

## Abstract

Gene expression requires two steps – transcription and translation – which can be regulated independently to allow nuanced, localized, and rapid responses to cellular stimuli. Neurons are known to respond transcriptionally and translationally to bursts of brain activity, and a transcriptional response to this activation has also been recently characterized in astrocytes. However, the extent to which astrocytes respond translationally is unknown. We tested the hypothesis that astrocytes also have a programmed translational response by characterizing the change in transcript ribosome occupancy in astrocytes using Translating Ribosome Affinity Purification(TRAP) subsequent to a robust induction of neuronal activity *in vivo* via acute seizure. We identified a change in transcripts on astrocyte ribosomes, highlighted by a rapid decrease in transcripts coding for ribosomal and mitochondrial components, and a rapid increase in transcripts related to cytoskeletal dynamics, motor activity, ion transport, and cell communication. This indicates a set of dynamic responses, some of which might be secondary to activation of Receptor Tyrosine Kinase(TRK) signaling. Using acute slices, we quantified the extent to which individual cues and sequela of neuronal activity can activate translation acutely in astrocytes. We identified both BDNF and ion concentration changes as contributors to translation induction, with potassium using both action-potential sensitive and insensitive components. We showed this translational response requires the presence of neurons, indicating the response is non-cell autonomous. We also show that this induction of new translation extends into peripheral astrocyte processes (PAPs). Accordingly, proteomics following fear conditioning in mice, showed that new translation influences peri-synaptic astrocyte protein composition *in vivo* under physiological conditions. Regulation of translation in astrocytes by neuronal activity suggests an additional mechanism by which astrocytes may dynamically modulate nervous system functioning.

**Main Points:** Astrocytes have a programmed, transcript-specific translational response to neuronal activity.

Both BDNF and K+, cues of neuronal activity, trigger this response.

This response requires the presence of neurons.

This response alters the astrocytic protein composition at the synapse.

## Introduction

Activity-dependent transcription of genes and translation of mRNA into proteins are both essential for long-term alterations of synaptic strength and efficacy, as well as for the consolidation of memory. This essentiality has been extensively shown with classical studies using pharmacological inhibitors in cultures, acute slices and behaving animals (Agranoff et al., 1966; Flexner et al., 1963; Kang and Schuman, 1996). At the transcriptional level, physiological activity, particularly calcium influx, is known to activate transcription factors, such as cyclic AMP response element binding protein (CREB), resulting in a cascade of gene expression changes in neurons and glia (Hardingham et al., 2001; Pardo et al., 2017; Yap and Greenberg, 2018). In neurons, this involves the initial induction of immediate early genes, such as *Fos* and *Jun*, transcription factors which support sustained alterations in synapses by increasing the production of a variety of transcripts encoding synaptic proteins.

Traditionally, when profiling mixed cultures or brain following stimulation, most of the transcriptional response to synaptic activity has been presumed to be derived from neurons, as this response can certainly be detected in purified populations of neurons. However, it is well known that other cell types—astrocytes, in particular—also respond to neuronal activity. Transcriptionally, neuronal maturation directly affects expression of the astrocyte glutamate transporter, GLT-1 (Perego et al., 2000; Swanson et al., 1997), and other genes (Hasel et al., 2017). Physiologically, astrocytes in the barrel cortex respond with an increase in cytosolic calcium after stimulation of whiskers (Wang et al., 2006). Furthermore, astrocytes increase *Fos* expression in a region-dependent manner (Chai et al., 2017). The RNAseq experiment performed by Chen et al.’s also detected robust changes in transcripts normally associated with glia (Chen et al., 2017). There is also a clear induction of astrocyte gene expression when stimulated in co-culture with neurons (Goudriaan et al., 2014; Hasel et al., 2017), and single-cell experiments in the visual cortex demonstrate astrocytes responding transcriptionally to visual stimulation as robustly as many classes of neurons (Hrvatin et al., 2018).

In addition, while a transcriptional response to activity in mixed cultures and brain has been widely studied, we (Dalal et al., 2017) and others (Chen et al., 2017; Cho et al., 2015) have recently shown that a distinct translational response is also occurring. By sequencing mRNA specifically bound to ribosomes, we assessed the change in occupancy of mRNAs by ribosomes in response to sustained depolarization (Dalal et al., 2017). While increased transcription of a gene often drove a corresponding increase in translation, at least 40% of the variance in translational response was independent of transcript levels. Likewise, Cho et al. applied ribosome footprinting to the hippocampi of animals exposed to a fear conditioning paradigm and quantified a substantial translational response (Cho et al., 2015). However, both of these studies were conducted on mixed populations of cell types, making the relative contributions of individual cell types difficult to assess. To this end, Chen et al. used Translating Ribosome Affinity Purification (TRAP) (Heiman et al., 2008; Sanz et al., 2009) to specifically enrich for ribosomes from hippocampal pyramidal neurons after long term potentiation, and sequenced the bound mRNA (TRAPseq). The authors identified a time-dependent enrichment of a variety of transcripts on neuronal ribosomes, much of which was undetectable in parallel RNAseq experiments from the same slices (Chen et al., 2017). This suggests that astrocytes might also be undergoing dynamic translational responses that are not apparent in prior transcriptional studies.

However, it has not been thoroughly assessed whether or not astrocytes have such a programmed *translational* response to cues of neuronal activity. A rapid translational response takes advantage of mRNA already present in the cell to produce new proteins without the time- and energy-consuming process of transcription. Furthermore, as dozens of ribosomes can bind a single transcript, regulation of translation can rapidly amplify protein production. Finally, regulation of translation can also allow for localized production of protein in specific subcellular compartments, such as near synapses, for which local translation has been observed both in the neuronal elements (Cajigas et al., 2012; Hafner et al., 2019; Ouwenga et al., 2017, 2018; Steward et al., 2015) and in peripheral processes of astroglia, including the perisynaptic astrocyte process (Boulay et al., 2017; Pilaz et al., 2016; Sakers et al., 2017).

We have previously shown that contextual fear-memory learning is accompanied by dynamic changes in both the neuronal as well as the astrocyte components of the hippocampal tripartite synapse (Rao-Ruiz et al., 2015). Astrocyte proteomic changes included downregulation of neurotransmitter transporters, such as SLC1A2/GLT-1, SLC1A3/GLAST and SLC6A11/GAT3. To what extend local translation is implicated in activity-induced changes in PAP protein levels remains to be determined. In this work, we set out to assess whether astrocytes also respond translationally to cues of neuronal activity *in vitro* and *in vivo*. We first use the TRAP method to test the hypothesis that a stimulus known to robustly activate neurons *in vivo*—a pentylenetetrazol (PTZ) induced seizure—will alter translation in astrocytes. We found that ribosomal occupancy of hundreds of transcripts is altered within minutes of seizure induction, whereas the vast majority of the corresponding genes had not yet changed transcription. To better understand this result, we turned to imaging-based methods in acute slices to quantify the alteration of translation in astrocytes in response to pharmacological manipulations that mimic aspects of neuronal activity. When performing these pharmacological studies in primary immunopanned astrocytes we show the bulk of these responses is lost in the absence of neurons. Finally, a proteomic analysis of synaptic fractions in contextual fear conditioned mice was performed to determine the *in vivo* effect of blocked translation on the peri-synaptic astrocyte proteome. We found that many of the proteomic changes induced by fear conditioning were blocked when translation was inhibited.

## Materials and Methods

### Mouse lines

All procedures were performed in accordance with the guidelines of Washington University’s Institutional Animal Care and Use Committee. Mice were maintained in standard housing conditions with food and water provided ad libitum and crossed at each generation to wildtype C57BL/6J mice from Jackson labs. The TRAP line was B6.FVB-Tg(Aldh1L1-EGFP/Rpl10a)^JD130Htz^, hereafter Astrocyte-TRAP (Doyle et al., 2008). Fear conditioning studies were conducted on wildtype C57BL/6J mice.

### PTZ treatment and TRAP

Three-week-old Astrocyte-TRAP mice of both sexes were intraperitoneally injected with PTZ at 60 mg/kg (Sigma P6500, 5 mg/ml stock solution in normal saline). Mice showing persistent convulsions at 8-10 minutes were subjected to rapid decapitation using decapicones (Braintree Scientific) for harvesting the brains. Control mice were injected with 110 μl normal saline (Sal) and decapitated similarly after 10 minutes. The harvested brains were snap-frozen in liquid nitrogen and stored at -80°C for TRAP. Six mice (three per treatment) were used.

TRAP was performed as described (Heiman et al., 2008) with a few modifications. Briefly, the brains were homogenized in ice in a buffer (20 mM HEPES pH 7.4, 150 mM KCl, 5 mM MgCl2, 0.5 mM dithiothreitol, 100 μg/ml cycloheximide (CHX), protease inhibitors, and RNase inhibitors). The lysates were cleared by centrifuging at 2000 xg for 10 min at 4°C and treated with DHPC (to 30 mM, Avanti) and NP-40 (to 1%, Ipgal-ca630, Sigma) for 5 min in ice. Lysates were further cleared by centrifuging at 20,000 xg for 15 min at 4°C. A 1/10^th^ volume of the cleared lysate was saved as the input control and used to generate RNAseq samples, and the rest was mixed with protein L-coated magnetic beads (Invitrogen), previously conjugated with a mix of two monoclonal anti-GFP antibodies (Doyle et al., 2008), and incubated with rotation for 4 h at 4°C. Beads were washed 5 times with a high-salt buffer (20 mM HEPES pH7.4, 350 mM KCl, 5 mM MgCl2, 1% NP-40, 0.5 mM dithiothreitol, and 100 μg/ml CHX) and finally resuspended in 200 ul normal-salt buffer (150 mM KCl, otherwise as above).

RNA was extracted from the input and TRAP samples using Trizol LS (Life Technologies) and a purification kit (Zymo Research, R1014) then quality-tested using RNA Pico Chips and BioAnalyzer 2100 (Agilent Technologies). All RIN values were > 8.

### RNAseq and TRAPseq

Libraries were prepared from 1 ug RNA using a library prep kit (NEB #7770), rRNA depletion kit (NEB #E6310), and Ampure XP beads (Beckman #A63881), and following manufacturer’s instructions. Prior to sequencing, quality was tested using a High Sensitivity D1000 ScreenTape (Agilent Technologies). Each library was sequenced as 2×150 bp fragments to a depth of about 30 million reads using Illumina HiSeq3000 machines.

### RNAseq and TRAPseq analysis

Sequencing results were quality-tested using FastQC (version 0.11.7). Illumina sequencing adaptors were removed using Trimmomatic (version 0.38) (Bolger et al., 2014), and reads aligning to the mouse rRNA were removed using bowtie2 (version 2.3.5) (Dobin et al., 2013). Surviving reads were then aligned, using STAR (version 2.7.0d) (Langmead and Salzberg, 2012), to the mouse transcriptome (Ensembl Release 97). The number of reads mapped to each feature were counted using htseq-count (version 0.9.1). All data are available on GEO: GSE147830.

Differential expression analysis was done using edgeR (version 3.24.3) (Robinson et al., 2010). Only genes with > 5 CPM in at least 3 out of 12 samples were retained for further analysis (11,593 genes). A negative binomial generalized log-linear model (GLM) was fit to the counts for each gene. Then likelihood ratio tests (LRT) were conducted for comparing PTZ samples with Sal samples. The comparisons were listed as follow:

1. Ribosome Occupancy Changes = TRAPseq^PTZ^ - TRAPseq^Sal^
2. Transcriptional Changes = RNAseq^PTZ^ - RNAseq^Sal^
3. TRAP enrichment after PTZ = TRAPseq^PTZ^ - RNAseq^PTZ^
4. TRAP enrichment after Saline = TRAPseq^Sal^ - RNAseq^Sal^
5. TRAP enrichment, combined = (TRAPseq^PTZ^ + TRAPseq^Sal^) / 2 – (RNAseq^PTZ^ + RNAseq^Sal^) / 2

TRAP-enriched transcripts (Supplemental Table 1) were defined as the union of all transcripts significantly enriched by TRAP (FDR < 0.1) in the last three comparisons (#3-5). Transcriptionally altered transcripts (Supplemental Table 2), were defined by having FDR < 0.1 in comparison #2. Translationally altered astrocyte transcripts (Supplemental Table 3) were defined as the intersect of Supplemental Table 1 with the significant (FDR < 0.1) transcripts from comparison #1, but removing those that were altered transcriptionally (i.e., also present in comparison #2). Results of all comparisons are included in Supplemental Table 5. Code for these analyses are available on Bitbucket: https://bitbucket.org/jdlabteam/ptz_sal/src/master/RNASeq_analysis/.

### qPCR

PTZ and saline treatment, brain harvest, and TRAP were performed as described above. Only mice with a 10-minute seizure response were included. cDNA synthesis, reaction mixture setup, and qPCR were performed as per manufacturer’s instructions (QuantaBio #101414, Thermo # A25743, ABI QuantStudio 6Flex). Beta actin was used a loading control. The primers used were as follows:

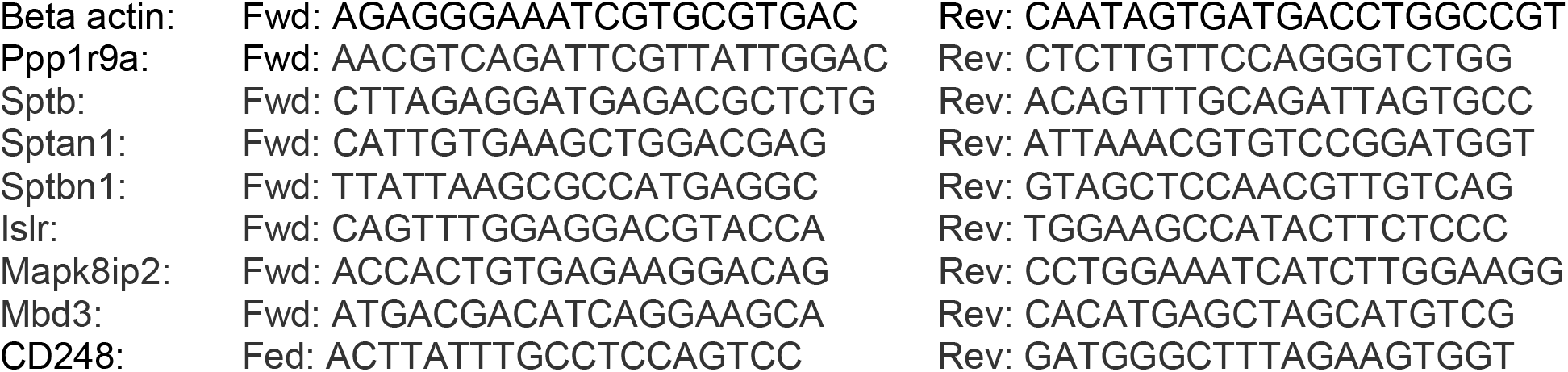

Fold changes in response to PTZ were calculated as 2^-DeltaDeltaCt^. A mixed linear model (DeltaCt ∼ treatment + 1|subject) was constructed, and the lme4 package was used in R to test DeltaCt as a function of PTZ treatment. P-value was calculated using a likelihood ratio test of the full mixed model with treatment against a model lacking treatment. Three animals per treatment were used.

### Gene Ontologies and pathway analysis of translationally regulated genes

The BiNGO (Maere et al., 2005) tool within Cytoscape was used to define and plot enrichment of Gene Ontology categories within differentially expressed genes. The *Mus musculus* GOSlim_Generic annotation was used, inputting either the list of 102 PTZ upregulated or 315 PTZ downregulated (Supplemental Table 3) astrocyte transcripts, and displayed categories reaching 0.05 significance by hypergeometric test after Benjamini-Hochberg correction for multiple testing. This analysis was conducted once in an exploratory setting, using the whole genome as a reference, and a second time using a more conservative reference of the 3,410 astrocyte-enriched transcripts (Supplemental Table 1), as highlighted in results. Full GO results are included in Supplemental Data 5. The same gene lists were examined using the COMPBIO literature mining tool (https://gtac-compbio.wustl.edu/), with default parameters.

### Comparison to curated gene lists

TRAP enriched transcripts were compared to prior markers of astrocytes as defined by a pSI < 0.01 in a prior TRAP experiment (Dougherty et al., 2012), a pSI < 0.01 of the immunopanned astrocyte expression (Zhang et al., 2014), analyzed as described (Ouwenga et al., 2017), and transcripts with a log2FC of >1, compared to nuclear transcriptomes of both neurons and oligodendrocytes (Reddy et al., 2017). TRAPseq upregulated and downregulated transcripts were further compared to the 102 genes identified by the TADA algorithm (Satterstrom et al., 2018) as causing ASD, and those genes curated as syndromic by the SFARI Gene database (Abrahams et al., 2013), accessed 8/9/19. Finally they were also compared to locally translated transcripts in neurons (Ouwenga et al., 2017) or astrocytes (Sakers et al., 2017). All comparisons were conducted using a Fisher Exact Test, with the maximum number of genes set to the number measurably expressed in the current experiment (11,593).

### Imaging based analysis of translation in astrocytes *in vivo*

We sparsely labeled astrocytes, as previously described (Sakers et al., 2017). Briefly, 2 μl of AAV9-CBA-IRES-GFP virus (concentration: 10^12^ vector genome (vg) / mL, obtained from the Hope Center Viral Vectors Core at Washington University) was injected at postnatal day 1-2 (P1-P2) pups bilaterally in the cortex, 1.5 mm lateral from the midline in two regions: 1 mm caudal to bregma and 2 mm rostral to lambda, using a 33g needle (Hamilton #7803-05) with a 50 μl Hamilton syringe (#7655-01).

At P21, acute cortical slices (300 µm) were prepared in artificial cerebrospinal fluid (aCSF, in mM: 125 NaCl, 25 glucose, 25 NaHCO_3_, 2.5 KCl, 1.25 NaH_2_PO_4_, equilibrated with 95% oxygen-5% CO_2_ plus 0.5 CaCl_2_, 3 MgCl_2_; 320 mOsmol) using a vibratome. 5 mM KCl, and 50ng/mL recombinant human/murine/rat BDNF (PeproTech, #450-02) were added to slices with 3μM puromycin (Tocris #40-895-0) in aCSF and allowed to incubate for 10 minutes at 37°C. In the anisomycin condition, 1 mM anisomycin (Sigma #A9789) in aCSF was added to slices 30 minutes before puromycin at 37°C. Slices were fixed with 4% paraformaldehyde in PBS for 30 minutes, followed by 30 minutes in 30% sucrose and freezing in OCT (Sakura #4583) for cryosectioning.

We cryosectioned 40 μm sections into PBS and incubated with Chicken Anti-GFP (Aves, 1:1000), and Mouse Anti-Puromycin(PMY) (Kerafast, 1:1000) at room temperature, followed by detection with appropriate Alexa conjugated secondary antibodies (Invitrogen), and counterstained nuclei with DAPI. We performed confocal microscopy on an AxioImager Z2 (Zeiss). Representative images represent 2-3 animals per condition with total cell number indicated in the figure legends.

Fluorescence intensity of PMY and GFP area were quantified using custom macros in ImageJ (NIH) and subsequently analyzed in R 3.4.1 software. Data were analyzed by ANOVA using the car package in R and post-hoc pairwise t-tests with Bonferroni correction for multiple comparisons.

### Imaging based analysis of translation in astrocytes *in vitro*

Postnatal rat astrocytes were purified according to previously published immunopanning protocols (Foo et al., 2011; Li et al., 2019; Zhang et al., 2016). First, we coated separate immunopanning dishes (150 mm) with antibody against CD45 (BD 550539), Itgb5 (eBioscience 14-0497-82), and a hybridoma supernatant against the O4 antigen (Zhang et al., 2016). Next, cerebral cortices were dissected from P3 rat pups and digested the tissue into a single cell suspension using papain. We incubated dissociated cells sequentially on the CD45 and O4-coated panning dishes to remove microglia/macrophages and oligodendrocyte precursor cells. Astrocytes were isolated by incubating the single cell suspension on the Itgb5-coated panning dish. After washing away non-adherent cells, we lifted astrocytes bound to the Itgb5-coated dish using trypsin and plated them on poly-D-lysine coated plastic coverslips in a serum free medium containing Dulbecco’s modified Eagle’s medium (DMEM) (LifeTechnologies 11960069), Neurobasal medium (LifeTechnologies 21103049), sodium pyruvate (LifeTechnologies 11360070), SATO (Foo et al., 2011), glutamine (LifeTechnologies 25030081), N-acetyl cysteine (Sigma A8199) and HBEGF (Sigma E4643) on 24-well culture plates. We replaced half of the medium with fresh medium every 2-3 days.

AAV9-GFAP-LCK-CFP-MYC virus (concentration: 10^11^ vg / mL) was added to each well of rat astrocytes after 2 days *in vitro*. Medium was changed 24 hours after infection and analyzed cells 5 days after infection to obtain ∼30% sparse labeling prior to puromycylation. Astrocytes were treated with drugs in the same manner as acute slices with the following additions: KCl was tested at both 5 mM and 50 mM, tetrodotoxin (TTX) was tested at 1 µM (Abcam #120055) with and without KCl, and untreated wells were used as control. Duration of treatment was identical to acute slices methods, followed by two washes with fresh media and immediate 4% paraformaldehyde fixation.

Cells were permeabilized with 0.2% Triton X100 and blocked with 10% donkey serum concurrently. Then we incubated coverslips with blocking solution, rabbit anti-GFAP (Dako, 1:1500), and mouse anti-PMY (Kerafast, 1:1000) overnight at 4°C, followed by detection with appropriate Alexa conjugated secondary antibodies (Invitrogen, 1:1000), and counterstained nuclei with DAPI. We acquired 5-10 images per well on a Zeiss Imager M2 microscope with an AxioCam MRm camera. Three independent experiments were conducted. Fluorescence intensity of PMY signal was quantified by drawing regions of interest (ROI) around individual cells using ImageJ (NIH) and analyzed using R as above.

### Fear Conditioning

Two cohorts of mice were tested in a fear conditioning paradigm described previously (Maloney et al 2019 GBB, Nygaard et al 2021 GBB). All animals were habituated to handling for multiple days prior to the start of testing. During testing, all males were run first, followed by females. All animals received subcutaneous injections of either 50 mg/kg cycloheximide solution or saline solution 30 minutes prior to the first day of conditioned fear testing. The first cohort of mice (n = 16 males, 16 females) underwent all three days of conditioned fear testing as previously described to assess the effect of cycloheximide on fear recall. Testing occurred on three consecutive days, during which fear response was measured by quantifying percent of time freezing using FreezeFrame (Actimetrics) software. On the first day, animals were placed in the testing chamber, scented with mint extract, for five minutes. Freezing behavior was assessed during a two minute baseline period and followed by three minutes of training during which a 2 sec 1.0 mA foot shock was paired at the end of a 20 sec 80 dB tone to condition a fear response to the tone and context. The second day tested contextual fear recall for eight minutes by placing the mouse in the same chamber but without the tone or shock. On the third day, the animal was placed in a new chamber for unique context. During the first two minutes, baseline freezing to the new context was quantified, followed by eight minutes of the 80 dB tone to assess cued fear recall. One week after fear conditioning, animals were tested in a shock sensitivity protocol to assess reactivity to shock. Data were analyzed using SPSS (v27) and outliers or animals with unsuccessful injections were removed.

### Synaptosome isolation

A second cohort of male C57BL6/jJ males were treated with cycloheximide (n=9) or saline (n=10), then underwent only the first day of fear conditioning, as described above. Following shock they were sacrificed via cervical dislocation within four to six hours; the brain was removed, split in two, and the dorsal hippocampi were dissected, frozen on dry ice, then stored at -80°C until use. Synaptosomes were isolated on a discontinuous sucrose gradient as described previously (Gonzalez-Lozano et al., 2020). In brief, dorsal hippocampus was homogenized in homogenisation buffer (0.32M sucrose, 5mM HEPES, in PBS. pH 7.4 with protease inhibitor cocktail (Roche)) using a dounce homogenizer for 12 strokes at 900 rpm. After centrifugation at 1000g for 10 min, the supernatant was collected followed by centrifugation at 100,000g for 4h in a 0.85/1.2M sucrose gradient. Synaptosomes were collected from the interface, diluted, and centrifuged at 18,000g for 20 min to collect a synaptosome rich pellet. All steps were performed on ice or at 4°C settings.

### FASP in-solution digestion of proteins

Filter-aided Sample preparation (FASP) was performed to digest the samples. 10 µg of synaptosomes from each sample was incubated with 75 µl reducing agent (2% SDS, 100 mM TRIS, 1.33 mM TCEP) at 55°C for 1h at 900 rpm. Followed by incubation with MMTS for 15 min at RT. Next, samples were transferred to YM-30 filters (Microcon, Millipore) and 200 µl 8 M urea in 100 mM TRIS (pH 8.8) was added. Samples were washed with this solution five times by spinning at 14,000g for 10 min each time, followed by washing four times with 50 mM NH_4_HCO_3_. Finally, the samples were incubated with 100 µl of trypsin overnight in a humidified chamber at 37°C. Digested peptides were eluted from the filter with 0.1% acetic acid. The samples were dried using a SpeedVac and stored at -20°C.

### Mass spectrometry based analysis

Samples were loaded onto an Ultimate 3000 LC system (Dionex, Thermo Scientific) as described (Gonzalez-Lozano et al., 2020; Hondius et al., 2021; van der Spek et al., 2021). A generally used hippocampal synaptosomes library created in-house with MaxQuant Software was used to annotate proteins. Spectra were annotated against the Uniprot mouse reference database.

Data quality control and statistical analysis were performed by using the downstream analysis pipeline for quantitative proteomics (MS-DAP version 0.2.6.3, Koopmans et al., under review). Outliers were removed when the variation among replicates was too large observed by deviating distribution plots, or in case of disturbed protein detection observed by altered retention time plots. This led to a final sample size of n=8 for the control (FC/CTL fear conditioning with saline) condition and n=8 for CHX treated samples (FC/CHX fear conditioning with cycloheximide).

### Statistical data analysis

Peptide abundance values were normalized and the MSqRob algorithm was used for peptide-level statistical analysis. The threshold for significance was set at q<0.05. For the Synaptic Gene Ontology database SynGO (1.1; (Koopmans et al., 2019) analysis of synaptic proteins all detected proteins were used as background input for analysis. Significance was determined by FDR correction. Cell type enrichment analysis was performed based on published proteomics data by (Sharma et al., 2015), where we annotated proteins to a certain cell type when the expression was >2 fold higher for a specific cell type compared to the cell type with the second highest expression. Data visualisation and statistics were performed in R studio (RStudio Team (2020). RStudio: Integrated Development Environment for R. RStudio, PBC, Boston, MA URL http://www.rstudio.com/, version 1.3.1093,) and GraphPad Prism (GraphPad Prism version 9.1.2. for Windows, GraphPad Software, San Diego, California USA, www.graphpad.com). Enriched gene lists were analysed and visualized by BiNGO for molecular function enrichments, as described above, but using the set of all detectable proteins **(Suppl. Table 7)** as the ‘background’ for the enrichment.

## Results

### Acute seizure activity alters *in vivo* ribosome occupancy of transcripts in astrocytes

A fundamental aspect of astrocyte function in the CNS is sensing and responding to neuronal activity (Charles et al., 1991; Perea et al., 2009; Schipke and Kettenmann, 2004). Indeed astrocytes respond transcriptionally, morphologically and functionally to the presence of neurons and neuronal activity (Goudriaan et al., 2014; Hasel et al., 2017; Hrvatin et al., 2018) but whether changes in their translatome also occur remains mostly unexplored. Therefore, we set out to determine if a robust induction of neural activity would acutely alter the profile of mRNAs bound to ribosomes in astrocytes *in vivo*.

We induced seizures in Astrocyte-TRAP mice, which express GFP-tagged ribosomes in all astrocytes **(Fig. 1a**, and (Doyle et al., 2008)), using PTZ, and harvested brains after 8-10 minutes from the seizing mice and saline injected controls **(Fig. 1b)**. We focused on a single, short time interval with the intention to isolate the translational response independent of PTZ-induced transcription. We collected both transcripts bound to astrocyte ribosomes via the TRAP method, as well as a parallel total RNAseq from each sample, and sequenced all samples to a depth of 28-43 million reads. Multidimensional scaling analysis shows that samples cluster on one axis strongly by whether they are from TRAP or RNAseq, and another axis by whether the TRAP samples come from PTZ or saline treated mice, confirming reproducibility of the method, and indicating a robust translational response is occurring on astrocyte ribosomes **(Fig. 1c)**. We next compared the TRAPseq samples to the RNAseq samples and confirmed that TRAP defined astrocyte translated transcripts **(Fig. 1d, Suppl Table 1)**, and this profile was significantly enriched for transcripts enriched by prior TRAP studies (Dougherty et al., 2012), immunopanning (Zhang et al., 2014), and nuclear sorting based assessments of *in vivo* astrocyte gene expression (Reddy et al., 2017) (all p < 10E^-16^ by Fisher’s Exact Test). Thus, we confidently measured ribosome-bound mRNAs from astrocytes in stimulated and unstimulated animals.

**Figure 1:**
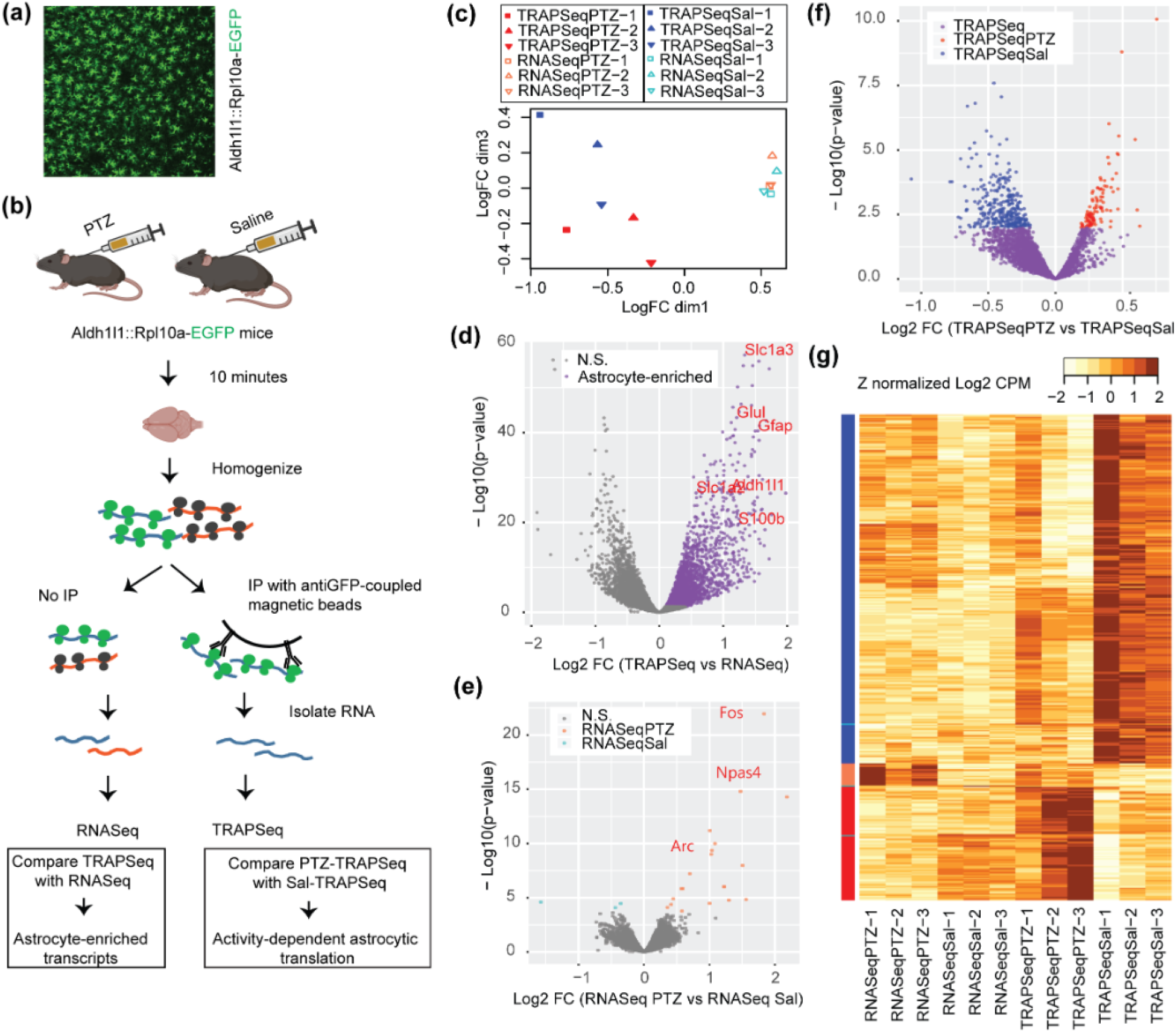
Seizure rapidly alters the translational profile of astrocytes. a) Representative image of an Astrocyte-TRAP mouse showing extensive expression of the GFP-tagged ribosome construct in astrocytes across the cortex. b) Schematic of immunoprecipitating astrocyte-specific mRNA, with and without seizure induction. c) Multidimensional scaling of twelve samples shows astrocyte TRAPseq samples are clearly separated from RNAseq samples, and that stimulated and unstimulated TRAPseq samples are also distinguished in this unsupervised clustering approach. d) Volcano plot of comparison of all TRAP samples to all RNAseq samples defines transcripts enriched on astrocyte ribosomes (purple) and these include known markers of astrocytes (red genes) as expected. e) Volcano plot comparing RNAseq from stimulated and unstimulated animals identifies the subset of transcripts responding rapidly to seizure induction (orange and blue transcripts, FDR<.1). Some of the immediate early genes are marked. f) Volcano plot comparing TRAPseq from stimulated and unstimulated animals identifies the subset of the TRAP enriched transcripts (purple) that were upregulated (red) or downregulated (blue) by seizure (FDR < 0.1). g) A heatmap of Z-normalized data for all transcriptionally upregulated (demarked by orange bar on left) or downregulated (cyan bars) transcripts, and astrocyte translationally downregulated (blue bar) or upregulated (red bar) transcripts across all conditions.

Next, we leveraged this data to determine whether transcripts alter their ribosome occupancy in response to this stimulation. First, we directly compared the PTZ and saline RNAseq data to test our assumption that using a short interval (8-10 minutes from injection to sacrifice) would allow us to exclude most of the transcriptional response, as it is slower. The assumption largely held, though there were genes that did show a significant transcriptional response, in this time window **(Fig. 1e, Suppl Table 2)**; however, these were very few (23, at FDR < 0.1), and unsurprisingly these were largely immediate early genes (e.g. *Npas4*, Log2FC = 2.2, FDR < 2E^-11^, *Fos* Log2FC = 1.8, FDR < 1.3E^-18^, *Arc*, Log2FC = 1.02, FDR < 1.6E^-6^) known to be part of the neuronal transcriptional response to synaptic activity. We next contrasted this with the number of transcripts with altered ribosome occupancy by comparing PTZ-TRAP to Saline-TRAP **(Fig. 1f)**. Because TRAP is an enrichment, but not perfect purification of astrocyte RNA, we conservatively filtered these to only pursue those transcripts that were also enriched by TRAPseq over RNAseq, as well as removed those that showed a transcriptional response, resulting in 417 transcripts (FDR < 0.1,**Suppl Table 3**, also see **Fig. 1g** and **Suppl. Table 4** for all transcripts and comparisons). Most of these showed only minimal changes in translation as we limited PTZ action to 8-10 minutes to focus exclusively on acute responses. For instance, only four transcripts exhibited a fold change ∼1.5. Yet, using qPCR, we could independently confirm these nominal changes for a majority of the candidates tested **(Fig. S1)**. We confirmed 5 out of 5 upregulated (*Mapk8ip2, Sptan1, Sptbn1, Sptb*, and *Ppp1r9a*) and 1 out of 3 downregulated (*CD248*, but not *Islr* and *Mdb3*, although the latter two still showed the expected trend) candidates, and clearly found PTZ treatment as the factor driving these translational changes (p < 0.05). Thus, overall, we discovered that astrocytes do have a robust translational response to stimulation *in vivo*, with at least 417 of the 3,410 astrocyte-enriched transcripts altering their ribosome occupancy, and with a larger proportion decreased (n = 315) rather than increased (n = 102) occupancy (**Fig. 1e**). Thus, a robust stimulation of neuronal activity induces immediate changes in astrocyte ribosome occupancy.

We then applied pathway analyses to these data to infer potential functions for the astrocytic translational response after seizure. First, examining the downregulated genes, the gestalt view provided by this analysis suggests an immediate decrease in ribosomal occupancy of transcripts involved in a variety of metabolic pathways **(Fig. 2a)**. For example, ribosomal transcripts (p < 3.03E^-7^, hypergeometric test, B-H corrected) are altered, as are key transcripts utilized by mitochondria (p < 8.37E^-14^), including numerous members of the electron transport chain. This suggests that robust activity in neurons triggers an immediate shift of protein production in astrocytes for metabolic functions. Next, we examined the upregulated genes and found robust increases in ribosome occupancy for transcripts encoding the cytoskeleton (p < 1.16E^-10^), notably actin binding proteins (p < 1.27E^-8^), as well as a variety of motor proteins (p < 9.68E^-6)^, and ion transporting proteins (p < 1.19E^-5^) **(Fig. 2b) (Suppl Table 5)**. The translational upregulation of cytoskeleton family transcripts suggests that an increase in neuronal activity poises the astrocyte for increased perisynaptic process motility. This supports a previous finding demonstrating that an increase in calcium induces astrocyte process motility (Molotkov et al., 2013), and that LTP induces PAP motility and retraction from the synapse (Henneberger et al., 2020; Perez-Alvarez et al., 2014). Furthermore, the overrepresentation of transcripts in ion transport suggest a homeostatic response to excessive neurotransmitter release induced by the seizure.

**Figure 2:**
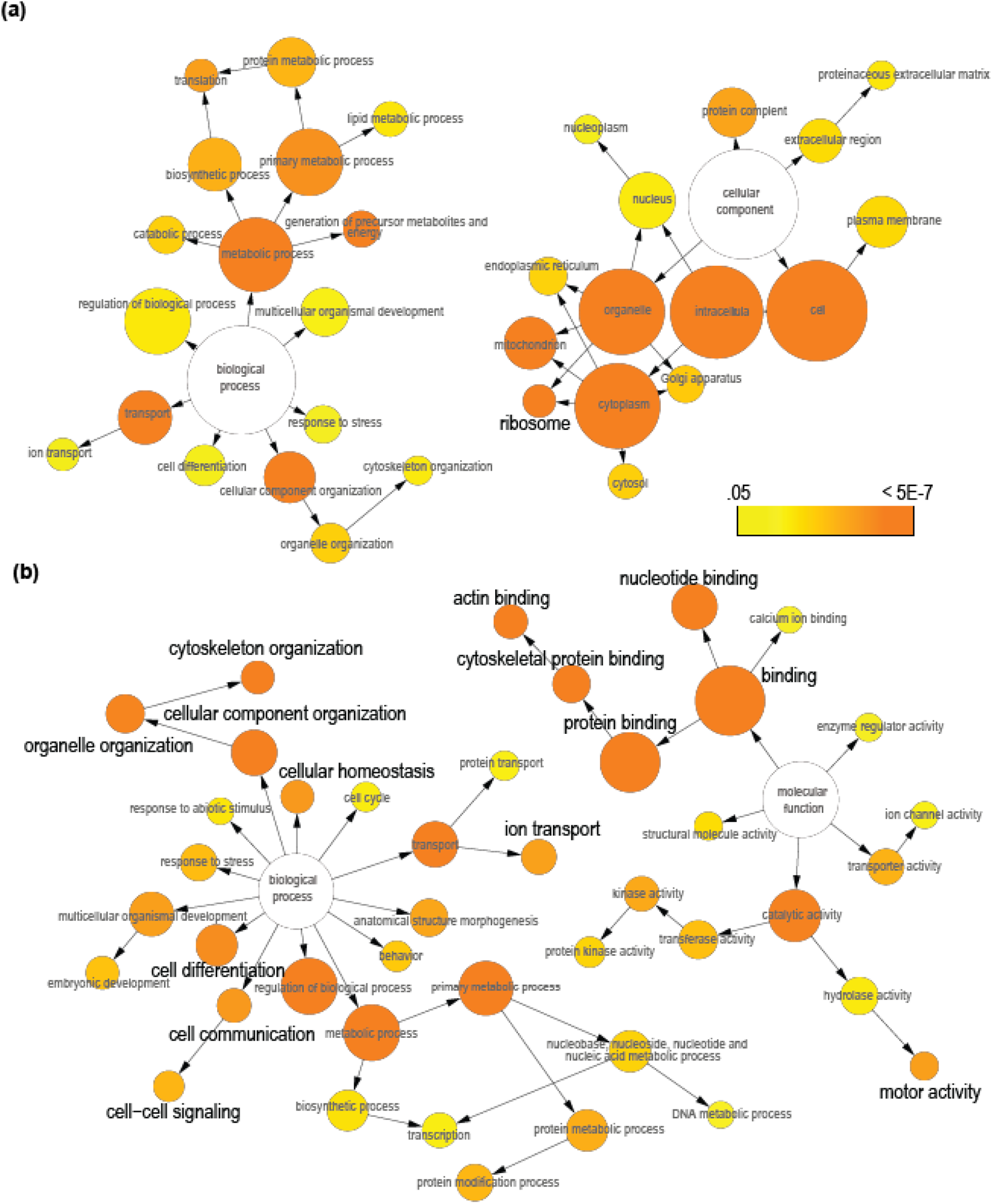
Translationally regulated genes show enrichment in specific Gene Ontology categories. a) Exploratory GO pathway analysis of downregulated genes identifies trends for transcripts in key metabolic processes (e.g. mitochondrial and ribosomal transcripts) showing changes in ribosome occupancy. b) Exploratory GO pathway analysis of upregulated genes indicates transcripts related to cytoskeleton/actin binding, motor function, ion transport, and cell-cell signaling as showing robust increases in ribosome occupancy. *Grey fonts: p < 0*.*05 for categorical enrichment compared to whole genome. Black fonts: p < 0*.*05 for categorical enrichment compared to only astrocyte-enriched genes. Color scale indicates significance for hypergeometric test. Category size is scaled to the number of genes. Arrows represent parent-child relationships in GO terms*.

Transcript specific regulation of translation is often mediated by the interaction of RNA-binding proteins (RBP) with specific sequences or RNA secondary structures, most often found in the 5’ and 3’ untranslated regions (UTRs). We therefore tested the hypothesis that the regulated genes shared particular sequence motifs. First, we examined the broad features of the 5’ and 3’ UTRs and found that the upregulated transcripts generally had UTRs with a longer length and a higher GC content (implying more stable secondary structures) than the downregulated transcripts **(Fig. S2a-d)**. Emboldened to consider the UTRs as being responsible for some of the transcript specific regulation, we then tested the hypothesis that there are specific motifs shared across transcripts. We utilized the AME tool within MEME suite (McLeay and Bailey, 2010) to screen for known vertebrate RBP target motifs in the upregulated and downregulated transcripts, and compared their abundance. At FDR < 0.1, we found dozens of potential regulatory motifs differing between the sets, which could be grouped into families based on their sequence similarity **(Fig. S2e,f)**. This suggests that the regulation of translation of these transcripts is mediated by sequences found in the UTR elements.

Finally, we sought to identify signaling pathways that might be upstream of these translationally regulated genes. We leveraged the COMPBIO tool for contextual language processing to mine Pubmed with the lists of translationally regulated genes and identify and cluster themes related to their gene products (https://gtac-compbio.wustl.edu/). COMPBIO reproduced aspects of the GO analysis, including themes related to ribosomes, cytoskeletal components, and mitochondria. Beyond what was found by GO, it highlighted a surprising connection to neurodegenerative diseases driven by the Huntington-related genes *Htt* and *Smcr8* as well as some forms of intellectual disability driven by the neurofibromatosis genes *Nf1* and *Nf2*, and the Cornelia de Lange syndrome genes *Smc3* and *Nipbl*. Indeed, the upregulated transcripts were significantly enriched in both recently empirically identified *de novo* autism genes (Fisher’s Exact Test, p < 0.0003)(Satterstrom et al., 2020), and a curated list of syndromic ASD/ID genes (p < 0.05) (Abrahams et al., 2013). Finally, this analysis also highlighted Ras and MAPK kinase signaling **(Fig. S3)**. These well studied intracellular signaling pathways are known to be downstream of neuronal activity-induced neurotrophic factors, such as BDNF. BDNF is a known regulator of translation and synaptic plasticity in neurons (Leal et al., 2014), and astrocytes are also known to express BDNF receptors and respond to BDNF (Aroeira et al., 2015; Colombo et al., 2012; Wang et al., 1998). For example, astrocytes have previously been reported to increase nitric oxide production in response to BDNF (Colombo et al., 2012), and indeed, we see increased ribosome occupancy of the key synthetase gene (*Nos1*) following stimulation **(Suppl Table 3)**. Others have detected an increase in global translation in astrocytes following BDNF stimulation in co-culture with neurons (Müller et al., 2015). We therefore turned to an imaging-based analysis of translation to assay this and other molecular signals that might directly induce a translational response in astrocytes.

### Astrocytes respond translationally to cues of neuronal activity in acute slices, including at distal processes

While our TRAP experiment demonstrates that a seizure-inducing stimulus can alter ribosome occupancy of transcripts, we cannot directly image new translation this way. Furthermore, stimulated neurons produce a variety of cues that could induce translation, such as excess glutamate and extracellular potassium changes, and it is unclear which of these the astrocytes are responding to. Therefore, we assessed how neuronal activity alters astrocyte translation acutely by exposing acute brain slices to pharmacological manipulations that trigger or mimic aspects of neuronal activity and measuring global translation in sparsely labeled astrocytes **(Fig. 3a)**. We first investigated whether BDNF stimulated translation of astrocyte transcripts in acute slices via immunofluorescent quantification of short-pulse puromycin (PMY) incorporation. Puromycin incorporates into the extending peptide chain on the translating ribosome and thus tags ongoing translation with an epitope that can be detected with immunofluorescence. BDNF induced a robust increase in puromycylation in astrocytes **(Fig. 3b,c)**, which is consistent with increased translation, as preincubation with another inhibitor of translation, anisomycin, blocked any PMY signal. BDNF transcription and secretion is regulated by neuronal activity (Balkowiec and Katz, 2000; Dieni et al., 2012; Zafra et al., 1990) and thus we hypothesized that globally altering neuronal firing regulates astrocyte translation. Therefore, we next depolarized the slices by elevating potassium using 5 mM KCl, a manipulation known to depolarize neurons and increase action potential frequency (refs). KCl alone also robustly increased puromycin incorporation in astrocytes, indicating that increased neuronal firing stimulates nascent protein synthesis in astrocytes **(Fig. 3b)**. Consistent with previous reports (Song and Gunnarson, 2012), we noted an increase in cell size after treatment with KCl **(Fig. S4**) suggesting that water permeability is also increased in astrocytes. However, even after normalizing the PMY intensity to cell size, quantified by astrocyte-GFP area, we still detected a robust increase in translation **(Fig. 3c)**, which argues against the possibility that increased cell size explains the greater PMY intensity. We next asked the reciprocal question of whether silencing neuronal firing results in a concomitant decrease in astrocyte translation by blocking all neuronal activity using tetrodotoxin (TTX). TTX alone significantly decreased puromycylation in astrocytes **(Fig. 3c)**, suggesting at least some of the basal level of translation in astrocytes is dependent on spontaneous activity of surrounding neurons. However, activity-independent translation also occurs in astrocytes because TTX-treated slices still contained significantly more puromycylation compared to anisomycin-treated slices **(Fig. 3c)**. Because KCl will also depolarize astrocyte membranes, we next investigated whether the induction of astrocyte translation by KCl was dependent on neuronal firing and subsequent neurotransmitter release. Therefore, we blocked neuronal firing with TTX prior to KCl stimulation. Indeed, pairing KCl treatment with TTX significantly inhibited most of the KCl induced increase in translation. However, translation was still higher than when TTX was used alone **(Fig. 3c)**. Thus, these findings suggest that Potassium-induced translation in astrocytes has components that are both dependent and independent of neuronal activity.

**Figure 3:**
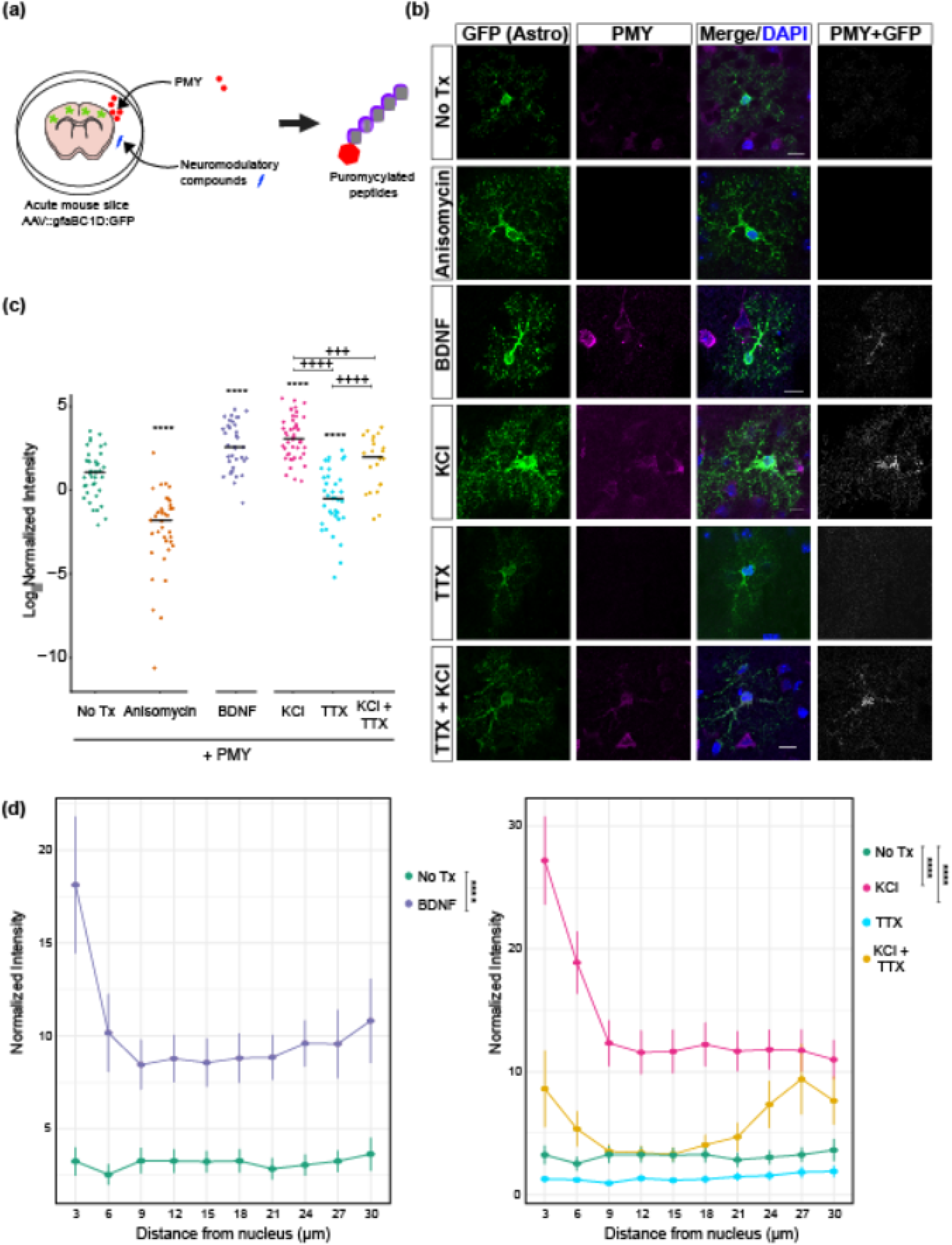
Astrocyte translation is modulated by BDNF and potassium. a) A schematic diagram showing an *ex vivo* translation assay. Acute brain slices were treated with test compounds, and the resulting translational changes were quantified using puromycin (PMY), which tags nascent peptides and serves as an epitope in the subsequent immunofluorescence staining experiment. b) Representative confocal images of P21 cortical astrocytes after puromycylation. Astrocytes were labeled with AAV9:GFP (green, see methods) and incubated for 10 minutes with PMY and indicated pharmacological manipulations. Immunostaining for GFP (green) and PMY (magenta) was performed. Puro+GFP channel indicates colocalized pixels of PMY and GFP, and was enhanced for publication. Scale bars = 10 µm. c) Quantification of PMY intensity in GFP astrocytes. Normalized intensity was calculated by dividing PMY intensity by GFP area (pixels). ANOVA was performed to determine effect of condition, F(5,211) = 53.389, p < 2.2E^-16^. Post-hoc pairwise t-tests were performed. Asterisks indicate comparison to No Treatment (Tx) (PMY only), plus signs indicate comparisons within KCl and TTX conditions. ^+++^p < 0.005, ^****/++++^p < 0.001. N_cells_(condition) = 40(No Tx), 37(Anisomycin), 36(BDNF), 44(KCl), 37(TTX), 23(KCl + TTX). (d) Quantification of PMY intensity at increasing distance from the cell soma from the data in (c). Normalized intensity was calculated by dividing PMY intensity by GFP area (pixels). A linear mixed model was performed to account for random variation from each cell / distance. Post-hoc Tukey’s test is represented with asterisks. **** p<0.0001. The cells used in this quantification are the same as in (c).

Astrocytes have been shown to have localized translation at their perisynaptic (Sakers et al., 2017) and vascular contacts (Boulay et al., 2017). We hypothesized that sensing chemical changes at the synapse stimulates localized translation in astrocytes, in an activity dependent manner. To test this, we used the images from Fig 4A-C, and quantified the abundance of puromycin signal in concentric rings drawn at increasing radial distances from the cell soma. To account for the difference in astrocyte territory size within each ring, we normalized the puromycin intensity levels to the GFP area. While BDNF and KCl strongly increased translation in the cell soma, there was a considerable increase in puromycin signal to the distal processes (**Fig. 3d**). Given the short incubation and subsequent washout of puromycin, this distal labeling is consistent with localized translation. Interestingly, while TTX reduced translation throughout the astrocyte, the combination of KCl and TTX substantially reduced translation in the soma and proximal processes yet not in the distal branches (**Fig. 3d**). These data point towards a model in which nascent translation in subcellular regions of the astrocyte respond differently to cues of neuronal activity.

**Figure 4:**
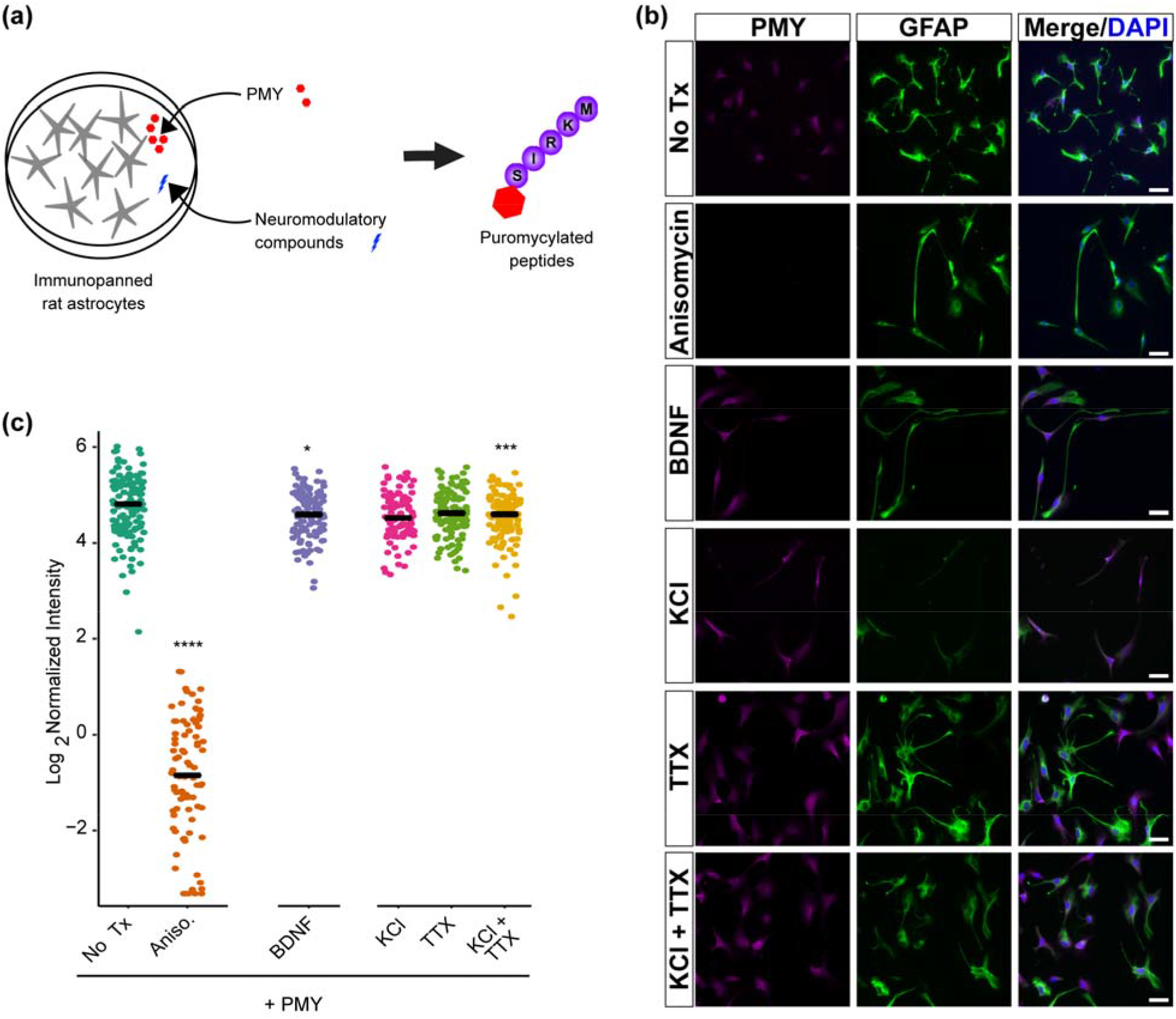
Impact of BDNF and KCl on astrocyte translation requires neurons. a) A schematic diagram of an assay for astrocyte-specific translation. Purified astrocytes are treated with test compounds, and the resulting translational response is measured using puromycin (PMY), which tags nascent peptides and is visualized in the subsequent immunofluorescence staining experiment. b) Representative fields of immunopanned astrocytes, stained for puromycin and GFAP. Scale bar = 50 µm. c) Quantification of PMY intensity in astrocytes. Mean intensity (signal/area) was calculated for individual cells. Pairwise t-tests were performed compared to No Tx. ****p < 0.001, ***p < 0.005, *p < 0.05. N_cells_(condition) = 129 (No Tx, i.e. PMY only), 87 (Anisomycin), 97 (KCl), 120 (TTX), 104 (BDNF), 111 (KCl+TTX).

### Activity-dependent astrocyte translation is non-cell autonomous

The TTX experiments above provided evidence that the presence of neurons and neuronal activity was largely required for the KCl response in astrocyte translation. We further tested this hypothesis by separating astrocytes from neurons and assessing their translational response to the same panel of chemical modulators **(Fig. 4a)**. We utilized a primary culture system in which mature astrocytes are immunopanned from dissociated rat cortices (Foo et al., 2011; Zhang et al., 2016). In this system, astrocytes exhibit morphological, physiological, and transcriptional profiles similar to those *in vivo* and thus make for a more direct comparison to our *ex vivo* data than traditionally cultured astrocytes. As before, we found that 1 mM anisomycin was sufficient to block puromycin signal in this system **(Fig. 4b,c)**. However, we found no other manipulation resulted in a significant upregulation on puromycin signal *in vitro*. Indeed, in contrast to our slice findings, we found a modest but significant downregulation of translation when astrocytes were treated with BDNF (**Fig 4c)**. This finding is reinforced by previous work using comparable methods, supporting the idea that astrocyte translational regulation is mediated by neuronal firing and/or neuronal contact (Müller et al., 2015). Further, we found another modest yet significant decrease in translation from the combination of KCl and TTX, but no effect of either TTX or KCl alone **(Fig. 4b,c)**. Together, these data suggest activity-dependent signaling from neurons induces astrocyte translation.

### Activity-dependent astrocyte translation alters the perisynaptic proteome under physiological conditions

Classic studies blocking new translation in the hippocampus during fear conditioning conclusively demonstrate that translation is required for memory consolidation (Debiec et al., 2002). We show in our experiments above that astrocytes can respond to manipulated cues of neuronal activity with a translational response. However, they don’t show whether they occur with physiological manipulations, nor the resulting changes in protein level. We therefore next set up an experiment to determine whether local translation is important for alterations of the peri-synaptic astrocytic proteome upon physiological stimulations that induce memory consolidation. We first replicated that cycloheximide before fear conditioning blocked hippocampus dependent memory formation in mice (**Fig. S5**). We then treated mice with cycloheximide or saline, fear conditioned both groups, and conducted unbiased proteomics on synaptic fractions from the dorsal hippocampus collected four hours later as done in (Rao-Ruiz et al., 2015) (**Fig. 5a**).

**Figure 5:**
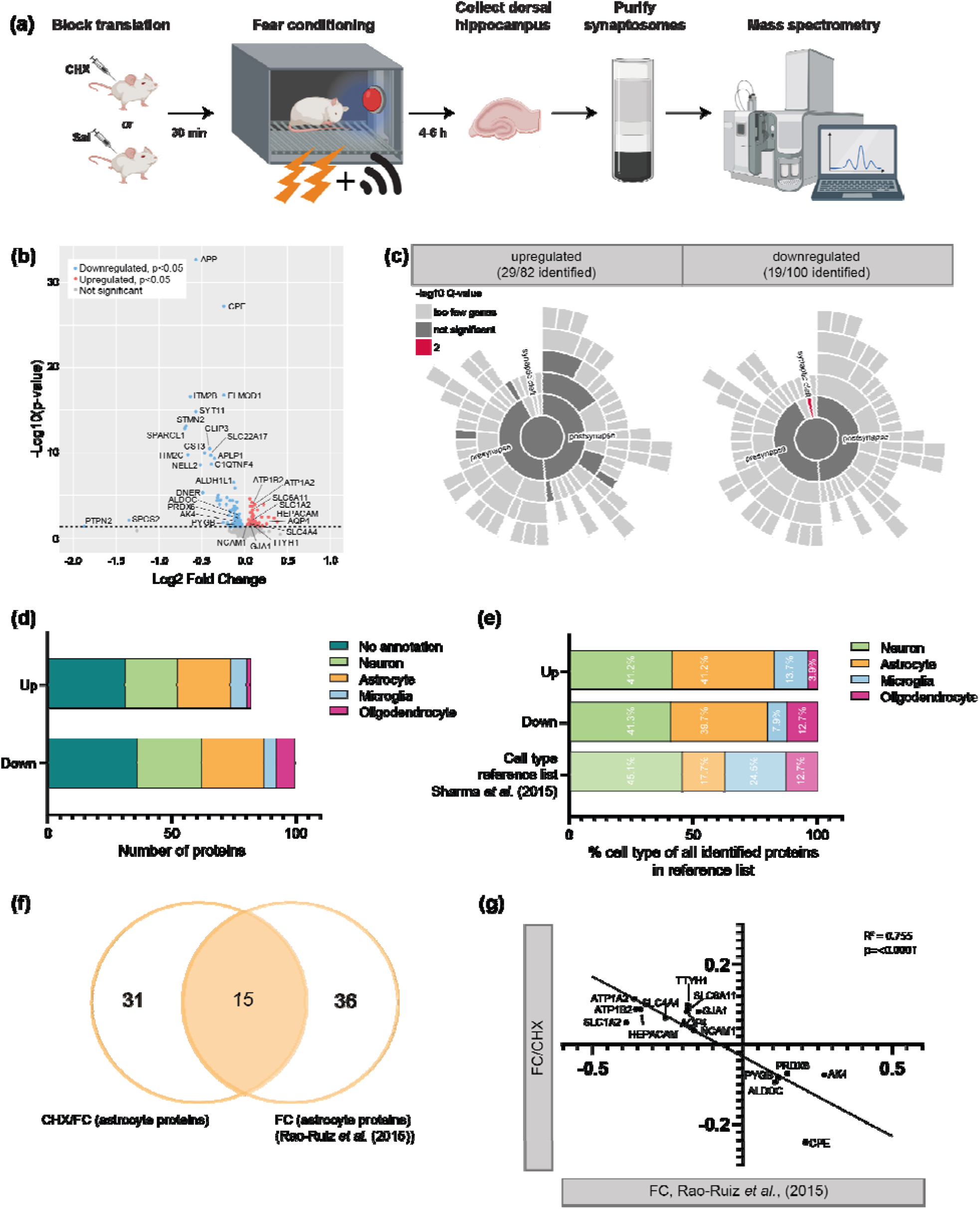
The astrocytic peri-synaptic proteome is altered when translation is inhibited by cycloheximide treatment. a) Experimental design of blocked translation by cycloheximide before fear conditioning followed by proteomic analysis on hippocampal synaptosomes. b) Volcano plot depicting proteomics data with expression changes in cycloheximide-treated mice (FC/CHX) compared to non-treated controls (FC/CTL) (X-axis, log2), and statistical significance (Y-axis). Downregulated, significant (p<0.05) proteins are shown by individual red dots, upregulated proteins by blue dots. Non-significant observed proteins in grey. The threshold of significance is represented by the dashed line. c) Sunburst plots showing the annotation in synaptic compartments using SynGO. Color coding is based on FDR-corrected p-values (q-values). d) Cell enrichment analysis based on cell type specific proteomics data by Sharma et al., (2015). Data is provided for all upregulated and downregulated proteins separated. Each bar is representing a different cell type, as depicted. e) Fractions of annotated cell types for proteins that are regulated by cycloheximide treatment, compared to the fractions of cell types identified in the reference list for cell type enrichment analysis by Sharma *et al*. (2015). The percentages are based on all identified regulated proteins with a cell type annotation, divided by “upregulated” and “downregulated”. f) A Venn-diagram showing similarities between astrocyte proteins regulated by FC/CHX and proteins regulated by FC only (Rao-Ruiz *et al*. (2015)). g) Correlation plot showing a high inverse correlation for changes in astrocyte protein levels caused by CHX (FC/CHX – FC/CTL, x-axis,) versus FC (Rao-Ruiz et al., 2015, Y-axis)

Across all samples, >3300 proteins and >19000 peptides were detected, with more than 15000 of these being identified in all samples (supplementary PDF). The data was reproducible across all replicates as shown by the analysis of abundance distributions, retention times and coefficient of variation computations (supplementary PDF). With this, we concluded that the data was of high quality and reliable to use for analysis.

A total of 182 proteins were significantly regulated (p<0.05) in the cycloheximide treated group (FC/CHX) compared to saline controls (FC/CTL). One hundred of these proteins were found to be downregulated, and 82 to be upregulated (**Fig. 5b**). A list of all detected proteins is present in supplementary table S7. A gene ontology analysis using BINGO was performed on the regulated proteins compared to astrocyte expressed genes, and identified coherent changes in proteins involved in ion regulation, as well as glycogen and glutathione modulation (**Fig. S6**)

We first determined the regulation of proteins that were of synaptic origin by using the Synaptic Gene Ontology database SynGO (Koopmans et al., 2019) Merely a small fraction of 29 out of 82 upregulated proteins and 19 out of 100 downregulated proteins was identified in the SynGO database (**Fig. 5c**). No significant enrichment was observed for a specific cellular component of the synapse, except for the “synaptic cleft” among the detected downregulated proteins (p=1.83e-3 after FDR correction). This involved the proteins APOE, SPACRL1, C1QL2 and C1QL3, which are all secreted proteins and of which at least the first two mentioned have been shown to be highly expressed by astrocytes (Sharma et al., 2015). To further explore the probable cellular origin of the regulated proteins, we performed a cell type specific enrichment analysis based on high-resolution proteomics data on single cell-types of Sharma *et al*. (Sharma et al., 2015); **Fig. 5d**). A large proportion of the regulated proteins are enriched in neurons (p=<0.0001), as expected. However, an equally sized proportion of regulated proteins is enriched in astrocytes (Fisher’s exact test, p=<0.0001). The observed fraction of astrocyte enriched proteins was more than expected by chance, based on the percentage of astrocytic proteins present in the reference list (**Fig. 5e**) unlike the minor fraction also found for microglia and oligodendrocyte enriched proteins. These findings are in line with previous observations that PAP proteins can be detected in synaptosomes(Carney et al., 2014; Chicurel et al., 1993; Rao-Ruiz et al., 2015; Shavit et al., 2011), and indicate that local translation affects the proteome of both the neuronal as well as the astrocytic elements of the activated synapse.

To obtain further insight into the regulation of locally translated PAP proteins by synaptic activity during memory consolidation, we compared our data to the earlier published dataset by Rao-Ruiz *et al*. (Rao-Ruiz et al., 2015) in which time dependent changes in the synaptic proteome, including 51 astrocyte enriched proteins, after contextual fear conditioning were identified. Interestingly, 15 of the 46 CHX-regulated astrocyte proteins were found to be regulated by contextual fear conditioning alone (**Fig. 5f)**. A strong inverse correlation was found for the direction of regulation of astrocyte proteins between the two data sets (Pearson R^2^=0.755, p=<0.0001; **Fig. 5g**), with the interesting observation that most of the astrocyte proteins were found upregulated by CHX (FC/CHX versus FC/CTL) while downregulated by fear conditioning alone (Rao-Ruiz dataset). A similar observation was done for the directional regulation of neuronal proteins although a much smaller overlap was found between neuronal proteins regulated by CHX or fear conditioning alone (**Fig. S7**). Interestingly, for a large portion of the astrocyte proteins regulated by both fear conditioning and FC/CHX, a peri-synaptic localization has previously been described: SLC1A2 (Chaudhry et al., 1995; Cholet et al., 2002; Minelli et al., 2001), SLC6A11 (Melone et al., 2015; Minelli et al., 1996), AQP4 (Nagelhus et al., 2004; Nielsen et al., 1997), ATP1A2 (Cholet et al., 2002), and PYGB (Richter et al., 1996).

Taken together, these data indicate that CHX treatment before shock is affecting local translation in astrocyte peri-synaptic processes, including proteins that could function in the formation of memory upon physiological stimulation.

## Discussion

Proper nervous system development and function requires precise spatiotemporal translation of proteins. Recently, there has been a focus on identifying transcriptional programs correlated with important developmental milestones (Kalish et al., 2018) and neuronal activity (Hrvatin et al., 2018). Neuronal activity can also induce such transcription in a cell-autonomous fashion, which has been shown to be important for synaptic plasticity and is classically mediated through activation of factors such as CREB (Yap and Greenberg, 2018). While CREB also induces activity-dependent transcriptional changes in astrocytes (Hasel et al., 2017; Pardo et al., 2017), neuronal activity-regulated translation in astrocytes has been mostly overlooked. Here, we show in acute slices that overall astrocyte translation changes after treatment with neuronal activity modulators but occurs only in the presence of neurons. Further, we profiled the changes in ribosome occupancy in astrocytes acutely after induction of seizure by TRAPseq and found a robust translational response, with hundreds of transcripts showing regulation. Pathway analyses highlight independent roles for upregulated and downregulated genes, and analyses of sequence features indicate that some of this regulation may be driven by shared motifs in UTR sequences. Literature mining tools suggested regulation by BDNF, and some relationship to genes implicated in neurodegeneration and neurodevelopmental disorder. Indeed, several genes involved in ASD, appear to show acute regulation of translation in astrocytes in response to neuronal activity.

The current study does have some limitations that should be mentioned. First, TRAP is a method for enrichment, but not perfect purification. Therefore, we conservatively limited our analysis for translationally regulated transcripts to those that were significantly enriched in astrocytes. It is possible that with additional strategies for enrichment, we might be able to identify even more transcripts that are changing ribosome occupancy. However, even with the current criteria we were still able to identify hundreds of transcripts. It is also worth mentioning that while assessment of ribosome occupancy does indicate regulation of translation, it is not necessarily identical to measurement of actual nascent protein production. Indeed, increased signal in TRAP might reflect an increase in stalled ribosomes on a given transcript rather than a recruitment of new ribosomes and increased translation. Thus, one should be cautious about inferring direction of changes in protein levels from changes in TRAP alone. Indeed, there was significant (Fisher Exact Test, OR 2.3, p < 0.0054), but not overwhelming overlap with our later proteomic studies of new protein translation and perisynaptic translation.

Nonetheless, it is clear from this study that astrocytes have a robust and transcript-specific translational response to neuronal activity. We found that both BDNF and KCl can induce translation in astrocytes. In agreement with our findings, a study using FUNCAT(Dieterich et al., 2010), an analogous method to puromycylation, found similar results of BDNF *in vitro* on astrocyte translation(Müller et al., 2015). Interestingly, KCl stimulation – a manipulaiton that can both trigger firing and modeling the high K+ concentrations that occur transiently after neuronal firing - has components that are both dependent on and independent of neuronal firing, as shown by TTX block. This may mean that there are discrete transcripts regulated by each neuronal signal, perhaps secondary to distinct secondary messenger events. This is supported by the discovery of motifs in subsets of the transcripts. We found motifs for over a dozen different RBPs in each transcript list – more than could be readily pursued experimentally here. Further, as many of these motifs are quite similar across RBPs, it can be a challenge to identify which, if any, of the known RBPs might be playing a role in this particular context. Nonetheless, it is possible that a RBP molecular code is defined by discrete signaling pathways which upon activation ultimately alter translation of specific transcripts downstream of each RBP. On the other hand, there could be a more non-specific mechanism where a variety of RBPs have redundant roles and work in a larger ensemble. For example, the set of RBPs identified as enriched in the upregulated transcripts is remarkably similar to the motifs found in our recent study of Celf6 binding targets (Rieger et al., 2020), even though the actual transcripts are distinct. As Celf6 and related Celf proteins have a role in forming RNA granules for mRNA storage and/or decay, and these granules often contain numerous species of RBPs and mRNAs interacting non-specifically through phase separation mediated by disordered domains, this suggests that transcripts enriched in binding for this selection of proteins may be more likely sequestered into a common granule awaiting release and translation following neuronal activation. What is common to both studies is that there was a GC bias in both lists, and thus there could be shared preference for GC binding RBPs in both cases. More sophisticated microscopy to track individual mRNAs, and a targeted proteomics study of some of these proteins in the context of astrocytes, might be able to elucidate which RBPs play a role in the regulation described here. This could eventually help test whether distinct signals regulate specific transcripts or whether there is a more coordinated action across RBPs.

It is also interesting to consider the relationship between global translation regulation and the potential for local translation regulation at peripheral processes and endfeet (Boulay et al., 2017; Sakers et al., 2017). We detected robust activity-dependent translation in astrocyte distal processes after a brief pulse of puromycin and subsequent washout and fixation to prevent sustained nascent peptide labeling. Strikingly, the addition of TTX to KCl treated slices blunts the effect of KCl in the soma and proximal processes, but not in the distal processes. Without knowing the location of synapses relative to the astrocyte territory, we cannot claim that this localized translation is perisynaptic. Furthermore, mass spectrometry-based proteomics of synaptic fractions after physiologically stimulating the dorsal hippocampus revealed a requirement for new translation for 46 proteins that are enriched in astrocytes, and many have prior evidence of local translation(Sakers et al., 2017). While methods do not yet exist to inhibit just local translation without impacting the rest of the cell, these results strongly suggest local translation in astrocytes plays a role in dynamically adapting the perisynaptic astrocyte processes to the demands associated with learning at synapses. Likewise, it is intriguing that there is a highly significant overlap (p < 10E^-16^, Fisher’s Exact Test) between the transcripts identified as PTZ stimulated, particularly those that are downregulated, and those previously described as bound to astrocyte ribosomes in perisynaptic fractions (Sakers et al., 2017). This suggests that some of these transcripts may be stalled on ribosomes in the periphery awaiting some signal secondary to neuronal activation, at which point they are rapidly translated and the RNA is subsequently degraded, similar to what has been reported for Arc in neurons (Farris et al., 2014). This could provide a substrate for local and activity dependent production of certain proteins in specific processes of a given astrocyte. This would of course require a highly localized signal to activate such translational activity in a process-specific manner. Candidates could be calcium transients in microdomains, or regions of elevated kinase activity downstream of G protein-coupled receptors or receptor tyrosine kinase cell surface receptors such as those activated by BDNF. Our findings here motivate future studies aimed at understanding the potential for subcellular localization of activity dependent translation in astrocytes.

Reexamining our puromycylation studies reveal that some combinations of stimuli (e.g., KCl and TTX) could preferentially skew translation in one compartment compared to another (**Fig 3d**) consistent with a cue-specific regulation of local translation. Furthermore, mass spectrometry-based proteomics of synaptic fractions after physiologically stimulating the dorsal hippocampus revealed a requirement for new translation for the levels of at least 182 proteins, a significant fraction of which are expressed in astrocytes, and have prior evidence of activity dependent changes in perisynaptic abundance (**Fig 5**) (Rao-Ruiz et al., 2015). For instance, contextual fear memory-induced upregulation of the astrocyte proteins ALDOC, PYGB, PDX6, CPE and AK4 was found to be partially dependent on translation. Reduced expression of many of these proteins have been associated with impaired memory (Ji et al., 2017; Mathieu et al., 2016; Opii et al., 2008; Phasuk et al., 2021). Remarkable, also the contextual fear memory-induced downregulation of a large number of astrocyte proteins (**Fig. 5g**, Rao-Ruiz et al, 2015) was found to be blunted by a translation block. This included established PAP proteins involved in neurotransmitter uptake (SLC1A2, SLC6A11) and ion balance (ATP1A2) that serve in balancing the tone of neuronal activity during and after the cellular- and network-changes involved in learning. It remains to be determined how the downregulation of these proteins is dependent on translation but may involve diverse mechanisms; e.g. protein degradation, reduced PAP localization (Murphy-Royal et al., 2015) or activity-induced PAP retraction (Henneberger et al., 2020; Perez-Alvarez et al., 2014).

## Supporting information

Supplemental Pipeline PDF

## Acknowledgements

We thank R. Barve, R. Head and the COMPBIO team for their support, as well as Katie McCullough, Stuart Fass, and Mike Vasek. This work was supported by the NIH (5R21DA038458, 1R01NS102272, T32GM008151, and 1K99AG061231) and the NSF (DGE-1745038). MSJK was supported by ZonMw (7330508160). RK was supported by the WUSTL BioSURF program. Key technical resources were supported by NIH through the CTSA (UL1 TR000448) and the Siteman Cancer center (P30 CA91842). Viral vectors were packaged at the Hope Center Viral Vectors Core at Washington University, and fear conditioning equipment was provided by the Animal Behavior Core of Washington University.

## Supplemental Material

**Supplemental Table 1: TRAP enriched transcripts**

A list of 3,409 transcripts enriched by TRAP in any one of three comparisons: (TRAPseq^All^ vs RNAseq^All^), or (TRAPseq^PTZ^ vs RNAseq^PTZ^), or (TRAPseq^Sal^ vs RNAseq^Sal^). Table includes columns for Ensembl gene IDs, Ensembl gene name, full gene name, Log2 Fold Change (Log2FC), EdgeR p-value and FDR corrected p-value for each of these comparisons.

**Supplemental Table 2: Transcripts that respond to PTZ transcriptionally**

A list of 23 genes altered transcriptionally by PTZ (RNAseq^PTZ^ vs RNAseq^Sal^). Table includes columns for Ensembl gene IDs, Ensembl gene name, full gene name, Log2 Fold Change (Log2FC), EdgeR p-value and FDR corrected p-value for this comparison.

**Supplemental Table 3: Transcripts that respond to PTZ translationally**

A list of transcripts with upregulated (n = 102) or downregulated (n = 315) ribosomal occupancy in response to PTZ (TRAPseq^PTZ^ vs TRAPseq^Sal^), filtered to remove those responding transcriptionally (Supplemental Table 2), and to only include those enriched in TRAP (Supplemental Table 1). Table includes columns for Ensembl gene IDs, Ensembl gene name, full gene name, Log2 Fold Change (Log2FC), EdgeR p-value and FDR corrected p-value for this comparison.

**Supplemental Table 4: Mean expression and results of all comparisons across all expressed genes**

A list of 11,593 genes that were measurably expressed in the experiment (CPM > 5 in at least 3 out of 12 samples), with columns for Ensembl gene IDs, Ensembl gene name, full gene name, the p-value, FDR, and Log2FC results of any EdgeR comparisons described above, and mean expression for each sample type in CPM. Also included are columns (TRUE/FALSE) for whether the gene was considered TRAP enriched (Supplemental Table 1) or transcriptionally responsive (Supplemental Table 2).

**Supplemental Table 5: Full results tables for BiNGO pathway analysis.**

Four sheets with full Cytoscape/BiNGO outputs using GOSlims pathways. Upregulated and downregulated genes from Supplemental Table 3 were tested using a reference set of either the whole mouse genome or filtering the reference to include only the astrocyte TRAP enriched genes (Supplemental Table 1 – “astro back” sheets).

**Supplemental Table 6: Selected results from COMPBIO analysis**

Genes found within each theme, along with overall theme score, and the score for each individual gene within a theme as produced by COMPBIO analysis.

**Supplemental Table 7: All regulated (and/or detected) proteins by proteomics analysis**

A list of 3615 proteins detected by proteomics analysis, along with differential expression log fold change values, p-values, and FDR corrected p-values.

**Supplemental PDF: Full Output of MS-DAP analysis of proteomics data.**

Full R code and output for proteomics data using the Mass Spect. Downstream Analysis Pipeline. https://github.com/ftwkoopmans/msdap/

**Supplemental Figure S1:**
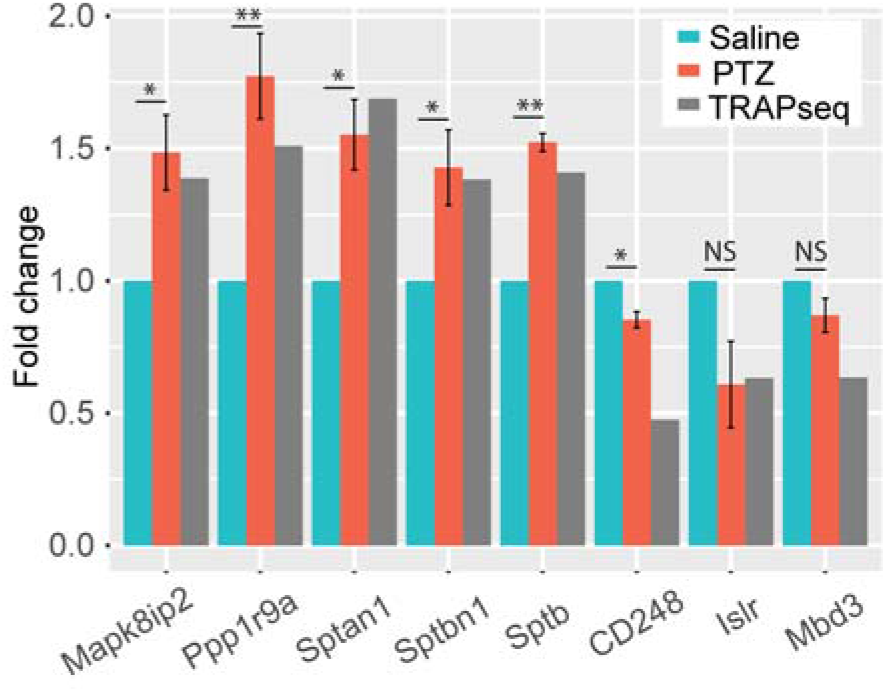
qPCR validates translational changes in independent samples. PTZ- and saline-treated mice were subjected to TRAP followed by qPCR, and fold changes calculated as 2^-DeltaDeltaCt^. Mixed linear model followed by likelihood ratio test was used to test DeltaCt as a function of PTZ treatment. Corresponding TRAPseq results are included for comparison. N = 3 mice per treatment. *, p-value ≤ 0.05; **, p-value ≤ 0.01

**Supplemental Figure S2:**
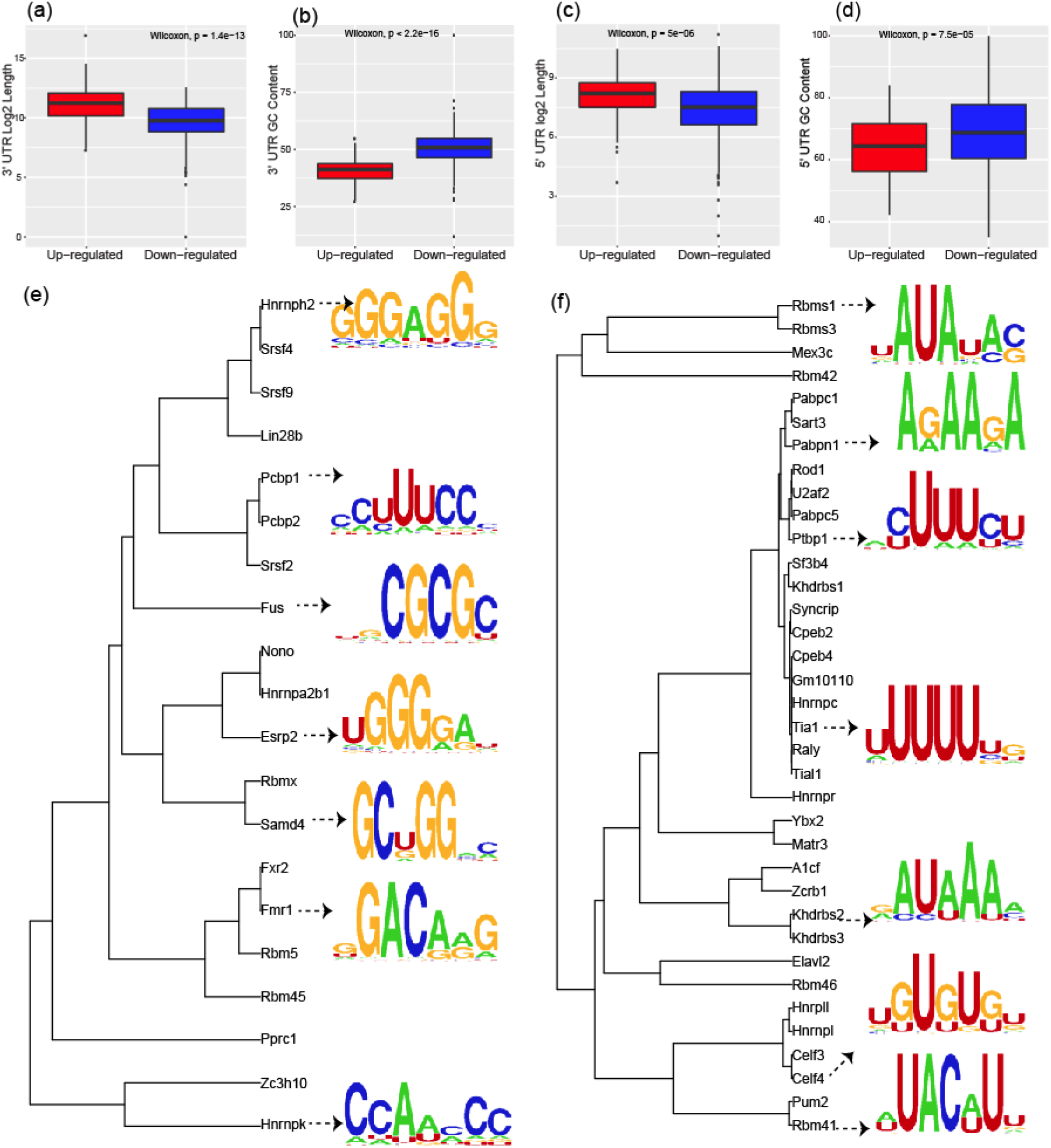
Translationally regulated transcripts show distinct sequence features and recurrent motifs. a-d) Translationally upregulated transcripts have significantly longer 3’UTR and 5’UTR (a and c) and lower GC contents in their 3’UTR and 5’UTR (b and d). e) Analysis of Motif Enrichment(AME) identifies dozens of RBPs with motifs that are significantly more common in 3’UTRs of the down-regulated transcript list. As many motifs are highly similar across RBPs, these have been clustered based on Euclidian distance in families of related factors. Examples of individual motifs for prototypical family members shown. *Motifs from RNAcompete data* (Ray et al., 2013) *http://hugheslab.ccbr.utoronto.ca/supplementary-data/RNAcompete_eukarya/Experiment_reports/RNAcompete_report_index.html* f) AME analysis identifies dozens of RBPs with motifs that are significantly more common in 3’UTRs of the up-regulated transcript list.

**Supplemental Figure S3:**
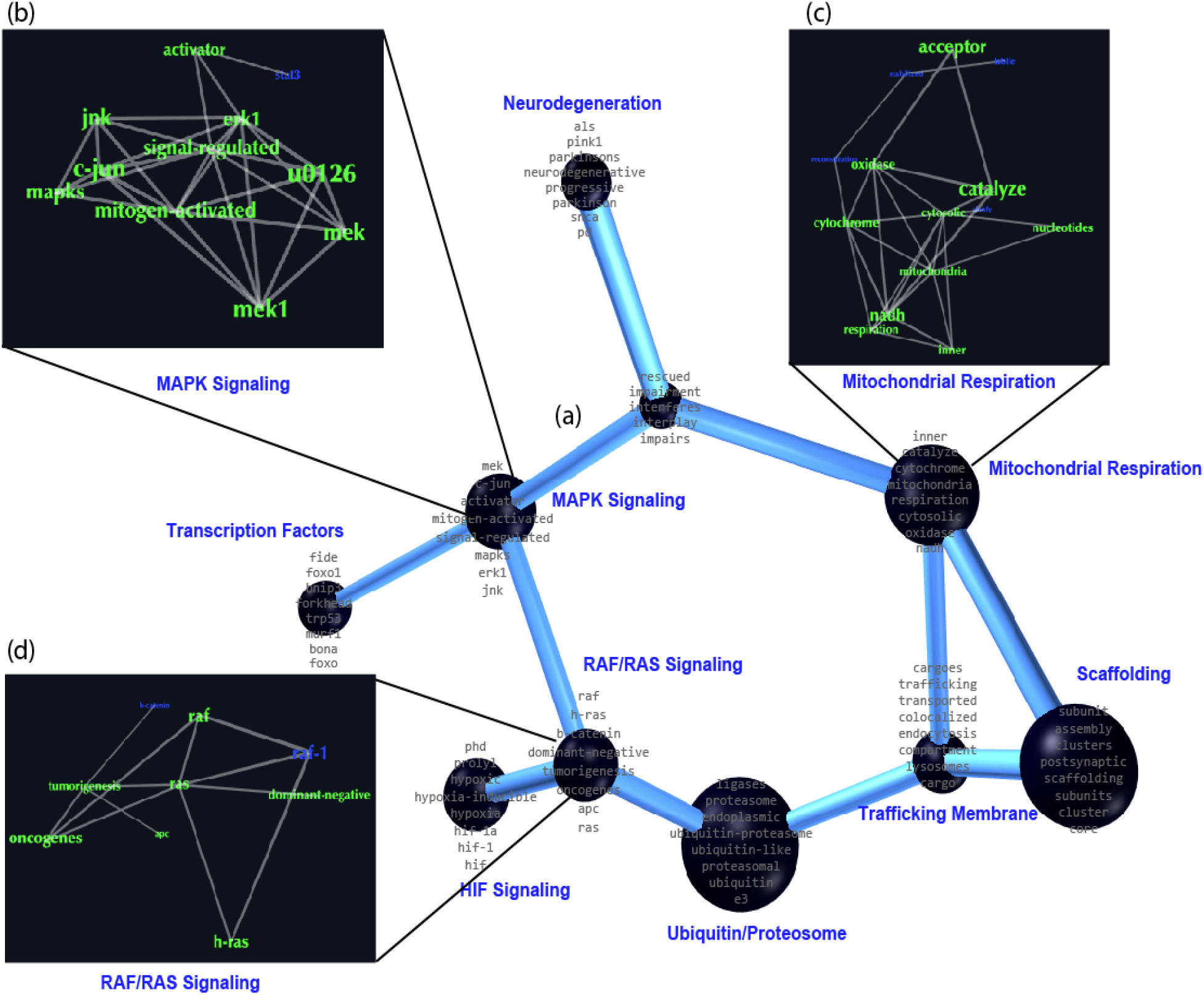
Literature mining through language processing highlights potential regulatory pathways for translationally regulated transcripts. a) Selected output of literature mining tool COMPBIO highlighting themes defined by terms co-occurring with the translationally regulated genes and with each other. Co-occurring terms are grouped into themes (nodes) that are connected by shared genes (edges). b-d) Relationships linking co-occurring terms within three example themes are shown. Supplemental table 6 includes the lists of genes connected with each theme, along with the scores as determined by permutation testing of similar sized lists of genes.

**Supplemental Figure S4:**
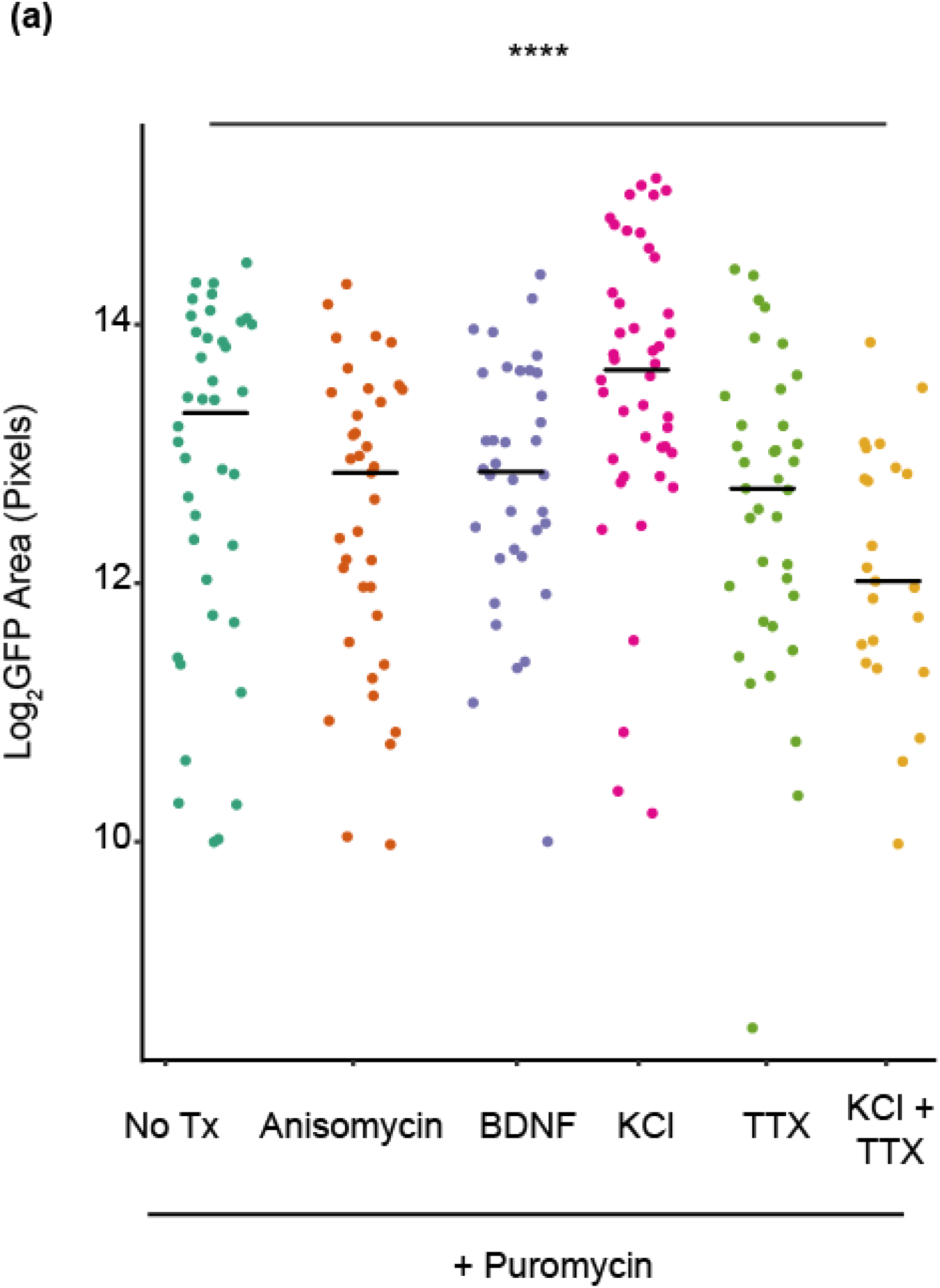
KCl depolarization alters size of astrocytes. a) Quantification of GFP pixel area of cells from Figure 4a. ANOVA was performed to determine the effect of conditions, F(5,211) = 5.7716, p = 5.13E^-5^. TukeyHSD post-hoc analyses reveal a significant decrease of astrocyte size with: Anisomycin treatment (p < 0.001), BDNF treatment (p = 0.02), TTX treatment (p = 0.004) and a significant increase of astrocyte size with KCl treatment (p < 0.001).

**Supplemental Figure S5:**
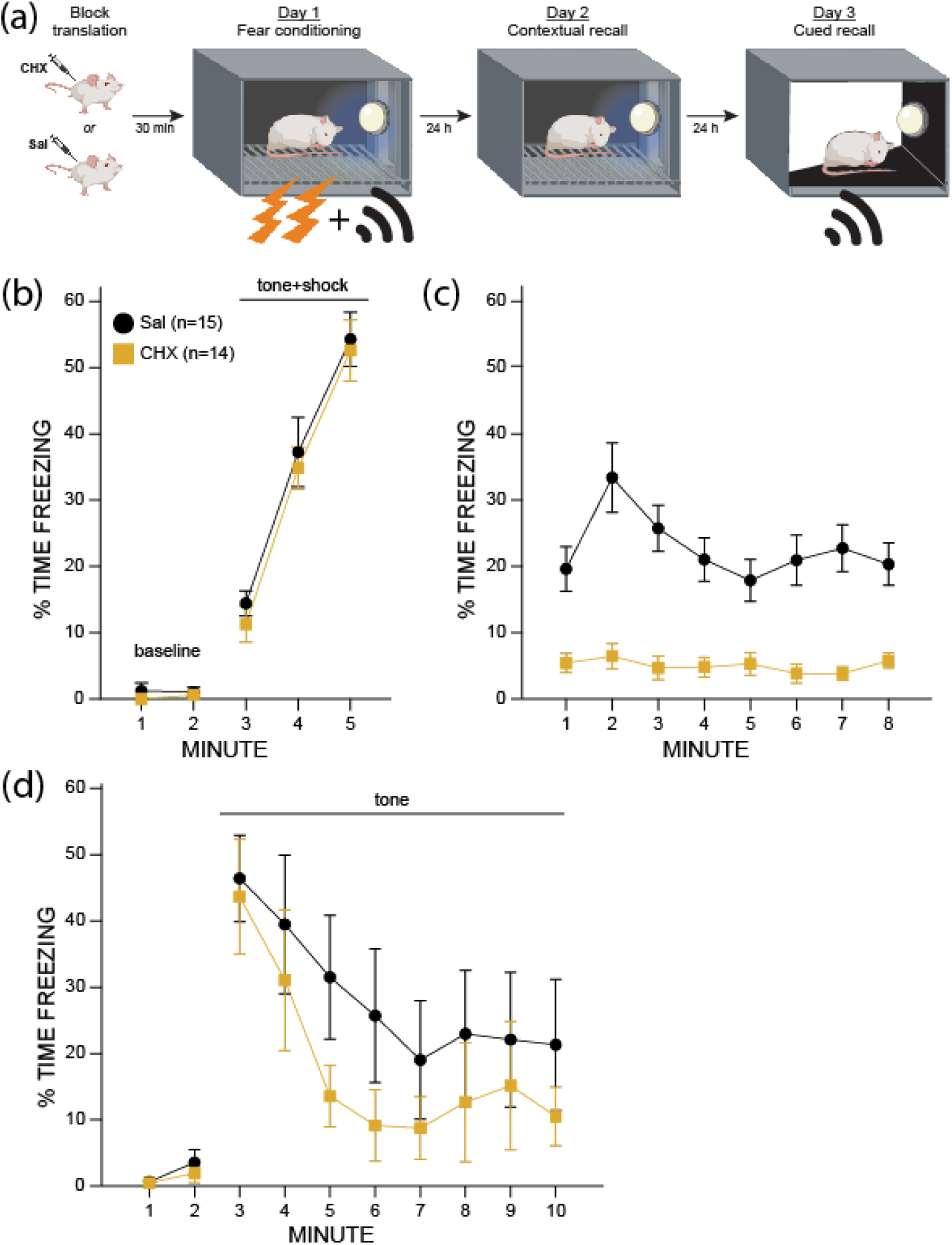
Translational inhibition blocks contextual fear recall. a) Schematic of fear conditioning paradigm with either cycloheximide or saline treatment. Day 1, mice are presented with paired foot shocks and tone. Day 2, mice are placed in the same context without any other stimulus. Day 3, mice are placed in a novel context and presented with the tone. Yellow squares represent cycloheximide-treated mice (n = 14, 7M/7F), black circles signify saline-treated animals (n = 15, 8F/7M). b) Percent of time mice freeze during the first day of fear conditioning. Tone was paired with foot shock at minutes 3, 4, and 5. Minutes 1 and 2 are baseline measures. c) Percent of time mice spent freezing in the contextual fear recall phase. CHX mice freeze less than saline: main effect of drug F(1,25)=45.434, p<0.0001. There was no effect of sex or interaction between drug and sex. d) Percent time freezing in a novel context. Tone was presented minutes 3 through 10. Minutes 1-2 indicate baseline freezing in the new environment.

**Supplemental figure S6:**
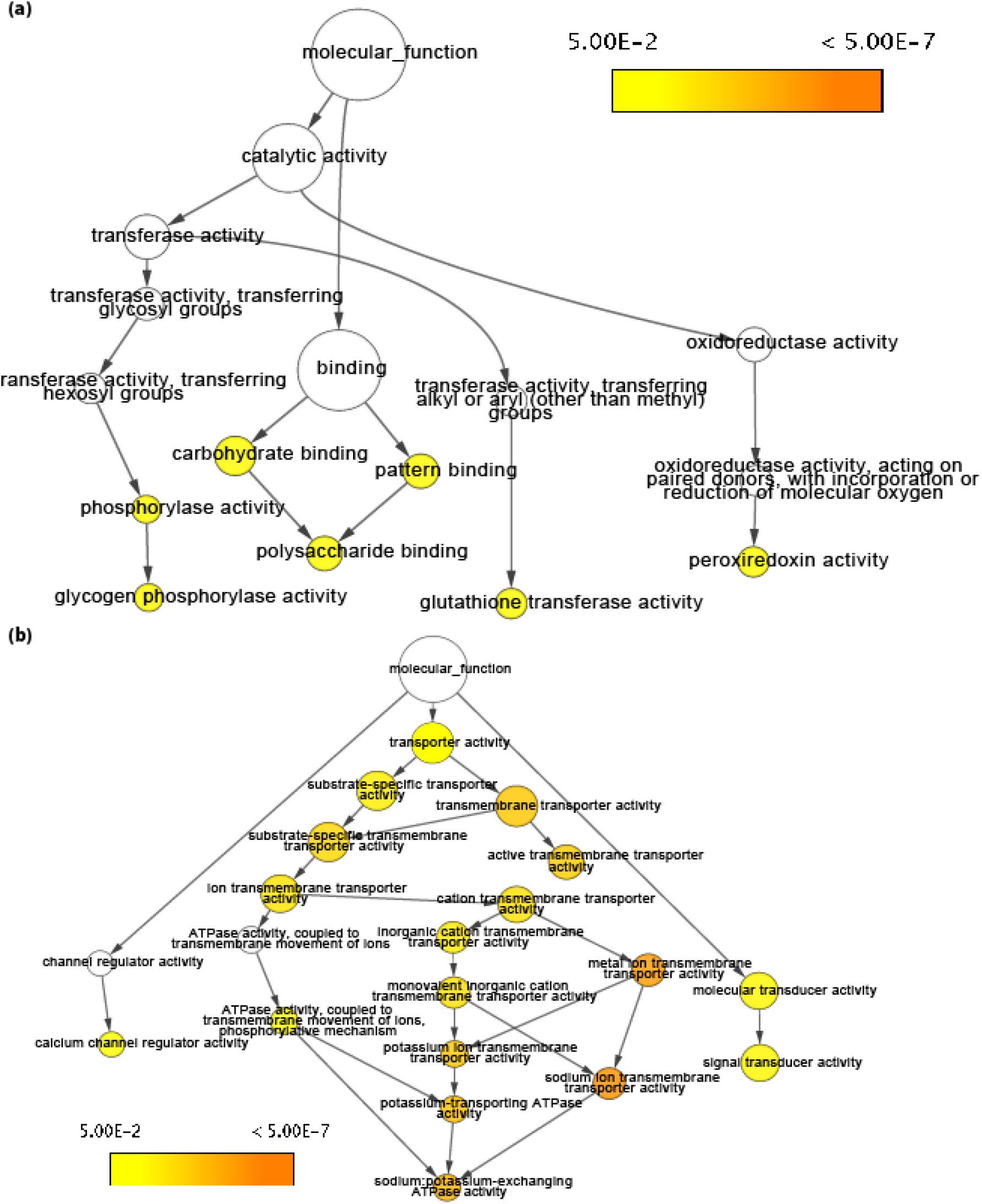
BinGO analysis reveals molecular functions enriched in proteins regulated by CHX treatment following fear conditioning. a) GO pathway analysis of downregulated proteins identifies categories related to glycogen, carbohydrate metabolism and gluthionine transferase. b) GO pathway analysis of upregulated proteins indicates proteins related to ATPase exchange, ion transport, and signal transduction are significantly enriched. *p < 0.05 for categorical enrichment compared to the full set of detectable proteins. Color scale indicates significance for hypergeometric test, corrected for multiple testing by Benjamin-Hochberg. Category size is scaled to the number of genes. Arrows represent parent-child relationships in GO terms*.

**Supplemental figure S7:**
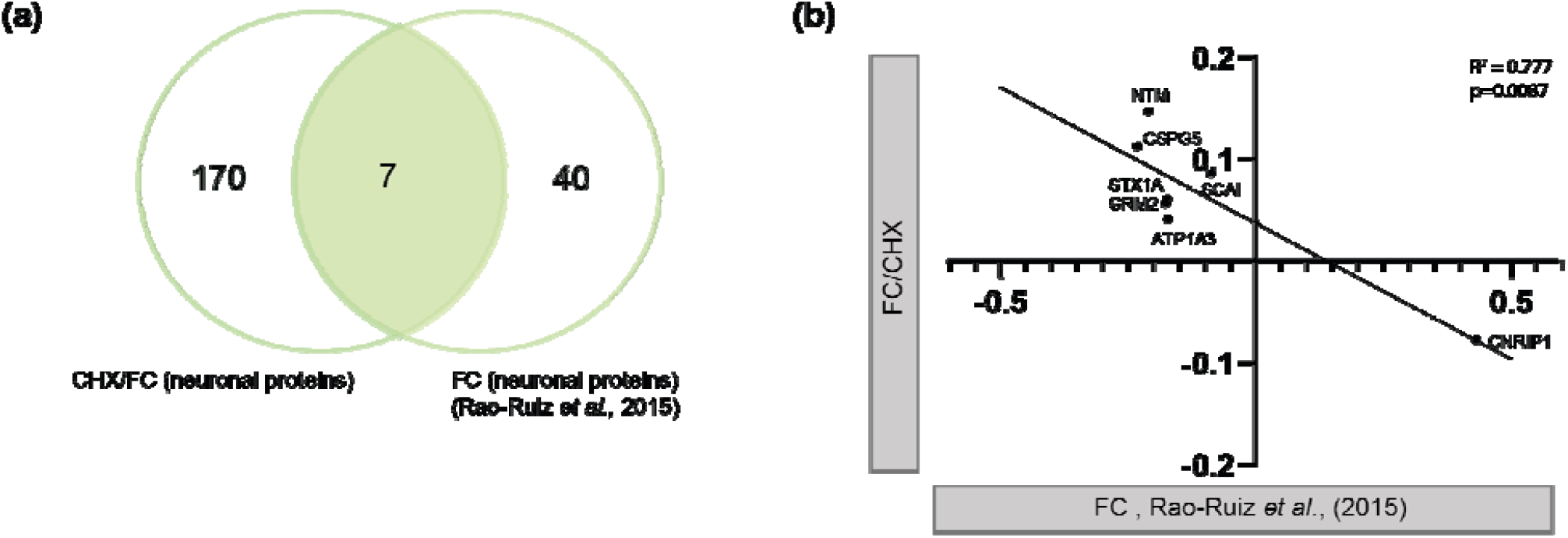
Inhibition of translation by cycloheximide alters the uniquely neuronal proteins observed in synaptosomes. (a) A Venn-diagram showing similarities between neuronal proteins regulated by FC/CHX and proteins regulated by FC only(Rao-Ruiz et al., 2015). **(b)** Correlation plot showing a high inverse correlation for changes in astrocyte protein levels caused by CHX (FC/CHX – FC/CTL, x-axis) versus FC only (Rao-Ruiz *et al*., 2015, Y-axis).

## References

Abrahams, B.S., Arking, D.E., Campbell, D.B., Mefford, H.C., Morrow, E.M., Weiss, L.A., Menashe, I., Wadkins, T., Banerjee-Basu, S., and Packer, A. (2013). SFARI Gene 2.0: a community-driven knowledgebase for the autism spectrum disorders (ASDs). Mol. Autism 4, 36.

Agranoff, B.W., Davis, R.E., and Brink, J.J. (1966). Chemical studies on memory fixation in goldfish. Brain Res. 1, 303–309.

Aroeira, R.I., Sebastião, A.M., and Valente, C.A. (2015). BDNF, via truncated TrkB receptor, modulates GlyT1 and GlyT2 in astrocytes. Glia 63, 2181–2197.

Balkowiec, A., and Katz, D.M. (2000). Activity-dependent release of endogenous brain-derived neurotrophic factor from primary sensory neurons detected by ELISA in situ. J. Neurosci. Off. J. Soc. Neurosci. 20, 7417–7423.

Bolger, A.M., Lohse, M., and Usadel, B. (2014). Trimmomatic: a flexible trimmer for Illumina sequence data. Bioinformatics 30, 2114–2120.

Boulay, A.-C., Saubaméa, B., Adam, N., Chasseigneaux, S., Mazaré, N., Gilbert, A., Bahin, M., Bastianelli, L., Blugeon, C., Perrin, S., et al. (2017). Translation in astrocyte distal processes sets molecular heterogeneity at the gliovascular interface. Cell Discov. 3, 17005.

Cajigas, I.J., Tushev, G., Will, T.J., Dieck, S. tom, Fuerst, N., and Schuman, E.M. (2012). The Local Transcriptome in the Synaptic Neuropil Revealed by Deep Sequencing and High-Resolution Imaging. Neuron 74, 453–466.

Carney, K.E., Milanese, M., van Nierop, P., Li, K.W., Oliet, S.H.R., Smit, A.B., Bonanno, G., and Verheijen, M.H.G. (2014). Proteomic Analysis of Gliosomes from Mouse Brain: Identification and Investigation of Glial Membrane Proteins. J. Proteome Res. 13, 5918–5927.

Chai, H., Diaz-Castro, B., Shigetomi, E., Monte, E., Octeau, J.C., Yu, X., Cohn, W., Rajendran, P.S., Vondriska, T.M., Whitelegge, J.P., et al. (2017). Neural Circuit-Specialized Astrocytes: Transcriptomic, Proteomic, Morphological, and Functional Evidence. Neuron 95, 531-549.e9.

Charles, A.C., Merrill, J.E., Dirksen, E.R., and Sandersont, M.J. (1991). Intercellular signaling in glial cells: Calcium waves and oscillations in response to mechanical stimulation and glutamate. Neuron 6, 983– 992.

Chaudhry, F.A., Lehre, K.P., van Lookeren Campagne, M., Ottersen, O.P., Danbolt, N.C., and Storm-Mathisen, J. (1995). Glutamate transporters in glial plasma membranes: highly differentiated localizations revealed by quantitative ultrastructural immunocytochemistry. Neuron 15, 711–720.

Chen, P.B., Kawaguchi, R., Blum, C., Achiro, J.M., Coppola, G., O’Dell, T.J., and Martin, K.C. (2017). Mapping Gene Expression in Excitatory Neurons during Hippocampal Late-Phase Long-Term Potentiation. Front. Mol. Neurosci. 10, 39.

Chicurel, M.E., Terrian, D.M., and Potter, H. (1993). mRNA at the synapse: analysis of a synaptosomal preparation enriched in hippocampal dendritic spines. J. Neurosci. Off. J. Soc. Neurosci. 13, 4054–4063.

Cho, J., Yu, N.-K., Choi, J.-H., Sim, S.-E., Kang, S.J., Kwak, C., Lee, S.-W., Kim, J., Choi, D.I., Kim, V.N., et al. (2015). Multiple repressive mechanisms in the hippocampus during memory formation. Science 350, 82–87.

Cholet, N., Pellerin, L., Magistretti, P.J., and Hamel, E. (2002). Similar perisynaptic glial localization for the Na+,K+-ATPase alpha 2 subunit and the glutamate transporters GLAST and GLT-1 in the rat somatosensory cortex. Cereb. Cortex N. Y. N 1991 12, 515–525.

Colombo, E., Cordiglieri, C., Melli, G., Newcombe, J., Krumbholz, M., Parada, L.F., Medico, E., Hohlfeld, R., Meinl, E., and Farina, C. (2012). Stimulation of the neurotrophin receptor TrkB on astrocytes drives nitric oxide production and neurodegeneration. J. Exp. Med. 209, 521–535.

Dalal, J.S., Yang, C., Sapkota, D., Lake, A.M., O’Brien, D.R., and Dougherty, J.D. (2017). Quantitative Nucleotide Level Analysis of Regulation of Translation in Response to Depolarization of Cultured Neural Cells. Front. Mol. Neurosci. 10, 9.

Debiec, J., LeDoux, J.E., and Nader, K. (2002). Cellular and systems reconsolidation in the hippocampus. Neuron 36, 527–538.

Dieni, S., Matsumoto, T., Dekkers, M., Rauskolb, S., Ionescu, M.S., Deogracias, R., Gundelfinger, E.D., Kojima, M., Nestel, S., Frotscher, M., et al. (2012). BDNF and its pro-peptide are stored in presynaptic dense core vesicles in brain neurons. J. Cell Biol. 196, 775–788.

Dieterich, D.C., Hodas, J.J.L., Gouzer, G., Shadrin, I.Y., Ngo, J.T., Triller, A., Tirrell, D.A., and Schuman, E.M. (2010). In situ visualization and dynamics of newly synthesized proteins in rat hippocampal neurons. Nat. Neurosci. 13, 897–905.

Dobin, A., Davis, C.A., Schlesinger, F., Drenkow, J., Zaleski, C., Jha, S., Batut, P., Chaisson, M., and Gingeras, T.R. (2013). STAR: ultrafast universal RNA-seq aligner. Bioinforma. Oxf. Engl. 29, 15–21.

Dougherty, J.D., Fomchenko, E.I., Akuffo, A.A., Schmidt, E., Helmy, K.Y., Bazzoli, E., Brennan, C.W., Holland, E.C., and Milosevic, A. (2012). Candidate pathways for promoting differentiation or quiescence of oligodendrocyte progenitor-like cells in glioma. Cancer Res. 72, 4856–4868.

Doyle, J.P., Dougherty, J.D., Heiman, M., Schmidt, E.F., Stevens, T.R., Ma, G., Bupp, S., Shrestha, P., Shah, R.D., Doughty, M.L., et al. (2008). Application of a translational profiling approach for the comparative analysis of CNS cell types. Cell 135, 749–762.

Farris, S., Lewandowski, G., Cox, C.D., and Steward, O. (2014). Selective Localization of Arc mRNA in Dendrites Involves Activity-and Translation-Dependent mRNA Degradation. J. Neurosci. 34, 4481–4493.

Flexner, J.B., Flexner, L.B., and Stellar, E. (1963). Memory in Mice as Affected by Intracerebral Puromycin. Science 141, 57–59.

Foo, L.C., Allen, N.J., Bushong, E.A., Ventura, P.B., Chung, W.-S., Zhou, L., Cahoy, J.D., Daneman, R., Zong, H., Ellisman, M.H., et al. (2011). Development of a method for the purification and culture of rodent astrocytes. Neuron 71, 799–811.

Gonzalez-Lozano, M.A., Koopmans, F., Sullivan, P.F., Protze, J., Krause, G., Verhage, M., Li, K.W., Liu, F., and Smit, A.B. (2020). Stitching the synapse: Cross-linking mass spectrometry into resolving synaptic protein interactions. Sci. Adv. 6, eaax5783.

Goudriaan, A., Camargo, N., Carney, K., Oliet, S.H.R., Smit, A.B., and Verheijen, M.H.G. (2014). Novel cell separation method for molecular analysis of neuron-astrocyte co-cultures. Front. Cell. Neurosci. 8.

Hafner, A.-S., Donlin-Asp, P.G., Leitch, B., Herzog, E., and Schuman, E.M. (2019). Local protein synthesis is a ubiquitous feature of neuronal pre-and postsynaptic compartments. Science 364, eaau3644.

Hardingham, G.E., Arnold, F.J.L., and Bading, H. (2001). Nuclear calcium signaling controls CREB-mediated gene expression triggered by synaptic activity. Nat. Neurosci. 4, 261.

Hasel, P., Dando, O., Jiwaji, Z., Baxter, P., Todd, A.C., Heron, S., Márkus, N.M., McQueen, J., Hampton, D.W., Torvell, M., et al. (2017). Neurons and neuronal activity control gene expression in astrocytes to regulate their development and metabolism. Nat. Commun. 8, 15132.

Heiman, M., Schaefer, A., Gong, S., Peterson, J.D., Day, M., Ramsey, K.E., Suárez-Fariñas, M., Schwarz, C., Stephan, D.A., Surmeier, D.J., et al. (2008). A translational profiling approach for the molecular characterization of CNS cell types. Cell 135, 738–748.

Henneberger, C., Bard, L., Panatier, A., Reynolds, J.P., Kopach, O., Medvedev, N.I., Minge, D., Herde, M.K., Anders, S., Kraev, I., et al. (2020). LTP Induction Boosts Glutamate Spillover by Driving Withdrawal of Perisynaptic Astroglia. Neuron 108, 919-936.e11.

Hondius, D.C., Koopmans, F., Leistner, C., Pita-Illobre, D., Peferoen-Baert, R.M., Marbus, F., Paliukhovich, I., Li, K.W., Rozemuller, A.J.M., Hoozemans, J.J.M., et al. (2021). The proteome of granulovacuolar degeneration and neurofibrillary tangles in Alzheimer’s disease. Acta Neuropathol. (Berl.) 141, 341–358.

Hrvatin, S., Hochbaum, D.R., Nagy, M.A., Cicconet, M., Robertson, K., Cheadle, L., Zilionis, R., Ratner, A., Borges-Monroy, R., Klein, A.M., et al. (2018). Single-cell analysis of experience-dependent transcriptomic states in the mouse visual cortex. Nat. Neurosci. 21, 120–129.

Ji, L., Wu, H.-T., Qin, X.-Y., and Lan, R. (2017). Dissecting carboxypeptidase E: properties, functions and pathophysiological roles in disease. Endocr. Connect. 6, R18–R38.

Kalish, B.T., Cheadle, L., Hrvatin, S., Nagy, M.A., Rivera, S., Crow, M., Gillis, J., Kirchner, R., and Greenberg, M.E. (2018). Single-cell transcriptomics of the developing lateral geniculate nucleus reveals insights into circuit assembly and refinement. Proc. Natl. Acad. Sci. 201717871.

Kang, H., and Schuman, E.M. (1996). A requirement for local protein synthesis in neurotrophin-induced hippocampal synaptic plasticity. Science 273, 1402–1406.

Koopmans, F., van Nierop, P., Andres-Alonso, M., Byrnes, A., Cijsouw, T., Coba, M.P., Cornelisse, L.N., Farrell, R.J., Goldschmidt, H.L., Howrigan, D.P., et al. (2019). SynGO: An Evidence-Based, Expert-Curated Knowledge Base for the Synapse. Neuron 103, 217-234.e4.

Langmead, B., and Salzberg, S.L. (2012). Fast gapped-read alignment with Bowtie 2. Nat. Methods 9, 357–359.

Leal, G., Comprido, D., and Duarte, C.B. (2014). BDNF-induced local protein synthesis and synaptic plasticity. Neuropharmacology 76, Part C, 639–656.

Li, J., Khankan, R.R., Caneda, C., Godoy, M.I., Haney, M.S., Krawczyk, M.C., Bassik, M.C., Sloan, S.A., and Zhang, Y. (2019). Astrocyte-to-astrocyte contact and a positive feedback loop of growth factor signaling regulate astrocyte maturation. Glia 67, 1571–1597.

Maere, S., Heymans, K., and Kuiper, M. (2005). BiNGO: a Cytoscape plugin to assess overrepresentation of gene ontology categories in biological networks. Bioinforma. Oxf. Engl. 21, 3448–3449.

Mathieu, C., Duval, R., Cocaign, A., Petit, E., Bui, L.-C., Haddad, I., Vinh, J., Etchebest, C., Dupret, J.-M., and Rodrigues-Lima, F. (2016). An Isozyme-specific Redox Switch in Human Brain Glycogen Phosphorylase Modulates Its Allosteric Activation by AMP * ♦. J. Biol. Chem. 291, 23842–23853.

McLeay, R.C., and Bailey, T.L. (2010). Motif Enrichment Analysis: a unified framework and an evaluation on ChIP data. BMC Bioinformatics 11, 165.

Melone, M., Ciappelloni, S., and Conti, F. (2015). A quantitative analysis of cellular and synaptic localization of GAT-1 and GAT-3 in rat neocortex. Brain Struct. Funct. 220, 885–897.

Minelli, A., DeBiasi, S., Brecha, N.C., Vitellaro Zuccarello, L., and Conti, F. (1996). GAT-3, a High-Affinity GABA Plasma Membrane Transporter, Is Localized to Astrocytic Processes, and It Is Not Confined to the Vicinity of GABAergic Synapses in the Cerebral Cortex. J. Neurosci. 16, 6255–6264.

Minelli, A., Barbaresi, P., Reimer, R.J., Edwards, R.H., and Conti, F. (2001). The glial glutamate transporter GLT-1 is localized both in the vicinity of and at distance from axon terminals in the rat cerebral cortex. Neuroscience 108, 51–59.

Molotkov, D., Zobova, S., Arcas, J.M., and Khiroug, L. (2013). Calcium-induced outgrowth of astrocytic peripheral processes requires actin binding by Profilin-1. Cell Calcium 53, 338–348.

Müller, A., Stellmacher, A., Freitag, C.E., Landgraf, P., and Dieterich, D.C. (2015). Monitoring Astrocytic Proteome Dynamics by Cell Type-Specific Protein Labeling. PloS One 10, e0145451.

Murphy-Royal, C., Dupuis, J.P., Varela, J.A., Panatier, A., Pinson, B., Baufreton, J., Groc, L., and Oliet, S.H.R. (2015). Surface diffusion of astrocytic glutamate transporters shapes synaptic transmission. Nat. Neurosci. 18, 219–226.

Nagelhus, E.A., Mathiisen, T.M., and Ottersen, O.P. (2004). Aquaporin-4 in the central nervous system: cellular and subcellular distribution and coexpression with KIR4.1. Neuroscience 129, 905–913.

Nielsen, S., Nagelhus, E.A., Amiry-Moghaddam, M., Bourque, C., Agre, P., and Ottersen, O.P. (1997). Specialized membrane domains for water transport in glial cells: high-resolution immunogold cytochemistry of aquaporin-4 in rat brain. J. Neurosci. Off. J. Soc. Neurosci. 17, 171–180.

Opii, W.O., Joshi, G., Head, E., Milgram, N.W., Muggenburg, B.A., Klein, J.B., Pierce, W.M., Cotman, C.W., and Butterfield, D.A. (2008). Proteomic identification of brain proteins in the canine model of human aging following a long-term treatment with antioxidants and a program of behavioral enrichment: Relevance to Alzheimer’s disease. Neurobiol. Aging 29, 51–70.

Ouwenga, R., Lake, A.M., O’Brien, D., Mogha, A., Dani, A., and Dougherty, J.D. (2017). Transcriptomic Analysis of Ribosome-Bound mRNA in Cortical Neurites In Vivo. J. Neurosci. 37, 8688–8705.

Ouwenga, R., Lake, A.M., Aryal, S., Lagunas, T., and Dougherty, J.D. (2018). The Differences in Local Translatome across Distinct Neuron Types Is Mediated by Both Baseline Cellular Differences and Post-transcriptional Mechanisms. ENeuro 5, ENEURO.0320-18.2018.

Pardo, L., Valor, L.M., Eraso-Pichot, A., Barco, A., Golbano, A., Hardingham, G.E., Masgrau, R., and Galea, E. (2017). CREB Regulates Distinct Adaptive Transcriptional Programs in Astrocytes and Neurons. Sci. Rep. 7, 6390.

Perea, G., Navarrete, M., and Araque, A. (2009). Tripartite synapses: astrocytes process and control synaptic information. Trends Neurosci. 32, 421–431.

Perego, C., Vanoni, C., Bossi, M., Massari, S., Basudev, H., Longhi, R., and Pietrini, G. (2000). The GLT-1 and GLAST glutamate transporters are expressed on morphologically distinct astrocytes and regulated by neuronal activity in primary hippocampal cocultures. J. Neurochem. 75, 1076–1084.

Perez-Alvarez, A., Navarrete, M., Covelo, A., Martin, E.D., and Araque, A. (2014). Structural and Functional Plasticity of Astrocyte Processes and Dendritic Spine Interactions. J. Neurosci. 34, 12738– 12744.

Phasuk, S., Pairojana, T., Suresh, P., Yang, C.-H., Roytrakul, S., Huang, S.-P., Chen, C.-C., Pakaprot, N., Chompoopong, S., Nudmamud-Thanoi, S., et al. (2021). Enhanced contextual fear memory in peroxiredoxin 6 knockout mice is associated with hyperactivation of MAPK signaling pathway. Mol. Brain 14, 42.

Pilaz, L.-J., Lennox, A.L., Rouanet, J.P., and Silver, D.L. (2016). Dynamic mRNA Transport and Local Translation in Radial Glial Progenitors of the Developing Brain. Curr. Biol. CB 26, 3383–3392.

Rao-Ruiz, P., Carney, K.E., Pandya, N., van der Loo, R.J., Verheijen, M.H.G., van Nierop, P., Smit, A.B., and Spijker, S. (2015). Time-dependent changes in the mouse hippocampal synaptic membrane proteome after contextual fear conditioning. Hippocampus 25, 1250–1261.

Ray, D., Kazan, H., Cook, K.B., Weirauch, M.T., Najafabadi, H.S., Li, X., Gueroussov, S., Albu, M., Zheng, H., Yang, A., et al. (2013). A compendium of RNA-binding motifs for decoding gene regulation. Nature 499, 172–177.

Reddy, A.S., O’Brien, D., Pisat, N., Weichselbaum, C.T., Sakers, K., Lisci, M., Dalal, J.S., and Dougherty, J.D. (2017). A Comprehensive Analysis of Cell Type–Specific Nuclear RNA From Neurons and Glia of the Brain. Biol. Psychiatry 81, 252–264.

Richter, K., Hamprecht, B., and Scheich, H. (1996). Ultrastructural localization of glycogen phosphorylase predominantly in astrocytes of the gerbil brain. Glia 17, 263–273.

Rieger, M.A., King, D.M., Crosby, H., Liu, Y., Cohen, B.A., and Dougherty, J.D. (2020). CLIP and Massively Parallel Functional Analysis of CELF6 Reveal a Role in Destabilizing Synaptic Gene mRNAs through Interaction with 31 UTR Elements. Cell Rep. 33, 108531.

Robinson, M.D., McCarthy, D.J., and Smyth, G.K. (2010). edgeR: a Bioconductor package for differential expression analysis of digital gene expression data. Bioinforma. Oxf. Engl. 26, 139–140.

Sakers, K., Lake, A.M., Khazanchi, R., Ouwenga, R., Vasek, M.J., Dani, A., and Dougherty, J.D. (2017). Astrocytes locally translate transcripts in their peripheral processes. Proc. Natl. Acad. Sci. U. S. A. 114, E3830–E3838.

Sanz, E., Yang, L., Su, T., Morris, D.R., McKnight, G.S., and Amieux, P.S. (2009). Cell-type-specific isolation of ribosome-associated mRNA from complex tissues. Proc. Natl. Acad. Sci. U. S. A. 106, 13939–13944.

Satterstrom, F.K., Kosmicki, J.A., Wang, J., Breen, M.S., Rubeis, S.D., An, J.-Y., Peng, M., Collins, R.L., Grove, J., Klei, L., et al. (2018). Novel genes for autism implicate both excitatory and inhibitory cell lineages in risk. BioRxiv 484113.

Satterstrom, F.K., Kosmicki, J.A., Wang, J., Breen, M.S., Rubeis, S.D., An, J.-Y., Peng, M., Collins, R., Grove, J., Klei, L., et al. (2020). Large-Scale Exome Sequencing Study Implicates Both Developmental and Functional Changes in the Neurobiology of Autism. Cell 180, 568-584.e23.

Schipke, C.G., and Kettenmann, H. (2004). Astrocyte responses to neuronal activity. Glia 47, 226–232.

Sharma, K., Schmitt, S., Bergner, C.G., Tyanova, S., Kannaiyan, N., Manrique-Hoyos, N., Kongi, K., Cantuti, L., Hanisch, U.-K., Philips, M.-A., et al. (2015). Cell type– and brain region–resolved mouse brain proteome. Nat. Neurosci. 18, 1819–1831.

Shavit, E., Michaelson, D.M., and Chapman, J. (2011). Anatomical localization of protease-activated receptor-1 and protease-mediated neuroglial crosstalk on peri-synaptic astrocytic endfeet. J. Neurochem. 119, 460–473.

Song, Y., and Gunnarson, E. (2012). Potassium Dependent Regulation of Astrocyte Water Permeability Is Mediated by cAMP Signaling. PLOS ONE 7, e34936.

van der Spek, S.J.F., Gonzalez-Lozano, M.A., Koopmans, F., Miedema, S.S.M., Paliukhovich, I., Smit, A.B., and Li, K.W. (2021). Age-Dependent Hippocampal Proteomics in the APP/PS1 Alzheimer Mouse Model: A Comparative Analysis with Classical SWATH/DIA and directDIA Approaches. Cells 10, 1588.

Steward, O., Farris, S., Pirbhoy, P.S., Darnell, J., and Driesche, S.J.V. (2015). Localization and local translation of Arc/Arg3.1 mRNA at synapses: some observations and paradoxes. Front. Mol. Neurosci. 7.

Swanson, R.A., Liu, J., Miller, J.W., Rothstein, J.D., Farrell, K., Stein, B.A., and Longuemare, M.C. (1997). Neuronal regulation of glutamate transporter subtype expression in astrocytes. J. Neurosci. Off. J. Soc. Neurosci. 17, 932–940.

Wang, X., Lou, N., Xu, Q., Tian, G.-F., Peng, W.G., Han, X., Kang, J., Takano, T., and Nedergaard, M. (2006). Astrocytic Ca2+ signaling evoked by sensory stimulation in vivo. Nat. Neurosci. 9, 816–823.

Wang, Y., Hagel, C., Hamel, W., Müller, S., Kluwe, L., and Westphal, M. (1998). Trk A, B, and C are commonly expressed in human astrocytes and astrocytic gliomas but not by human oligodendrocytes and oligodendroglioma. Acta Neuropathol. (Berl.) 96, 357–364.

Yap, E.-L., and Greenberg, M.E. (2018). Activity-Regulated Transcription: Bridging the Gap between Neural Activity and Behavior. Neuron 100, 330–348.

Zafra, F., Hengerer, B., Leibrock, J., Thoenen, H., and Lindholm, D. (1990). Activity dependent regulation of BDNF and NGF mRNAs in the rat hippocampus is mediated by non-NMDA glutamate receptors. EMBO J. 9, 3545–3550.

Zhang, Y., Chen, K., Sloan, S.A., Bennett, M.L., Scholze, A.R., O’Keeffe, S., Phatnani, H.P., Guarnieri, P., Caneda, C., Ruderisch, N., et al. (2014). An RNA-sequencing transcriptome and splicing database of glia, neurons, and vascular cells of the cerebral cortex. J. Neurosci. Off. J. Soc. Neurosci. 34, 11929–11947.

Zhang, Y., Sloan, S.A., Clarke, L.E., Caneda, C., Plaza, C.A., Blumenthal, P.D., Vogel, H., Steinberg, G.K., Edwards, M.S.B., Li, G., et al. (2016). Purification and Characterization of Progenitor and Mature Human Astrocytes Reveals Transcriptional and Functional Differences with Mouse. Neuron 89, 37–53.

