## Supplemental Pipeline PDF for "Activity dependent translation in astrocytes dynamically alters the proteome of the perisynaptic astrocyte process"

### MS-DAP: Mass Spectrometry Downstream Analysis Pipeline

version: beta 0.1.7.2    https://github.com/ftwkoopmans/msdap/

**3 Summary of statistical results** **45**

**4 log** **46**

**5 R command history** **47**

**6 R session info** **55**

### 1 Quality control

The quality control figures in this section investigate reproducibility and global clustering of samples by visualizing:

- number of features detected in each sample
- dataset completeness
- local effects in peptide retention time per sample
- reproducibility of peptide quantification among replicates
- PCA of all samples to visualize clustering

The first set of quality control figures describes individual samples, thereafter group-level quality metrics are described and finally sample clustering highlights structure in the entire dataset.

#### 1.1 number of peptides and proteins

These plots show the number of (target) peptides that are detected per sample. For DDA, 'detected' implies the peptide has a MS/MS identification whereas peptides quantified through match-between-runs (MBR) are quantified but not 'detected'. In case of DDA, we also show the number of peptides quantified through MBR. For DIA, we refer to a peptide as 'detected' if the confidence score (for identification) is at most 0.01.

Samples in this plot are sorted by their experimental group, and then ordered and by their name within each group. This data is also available in the output table 'samples.xlsx'.

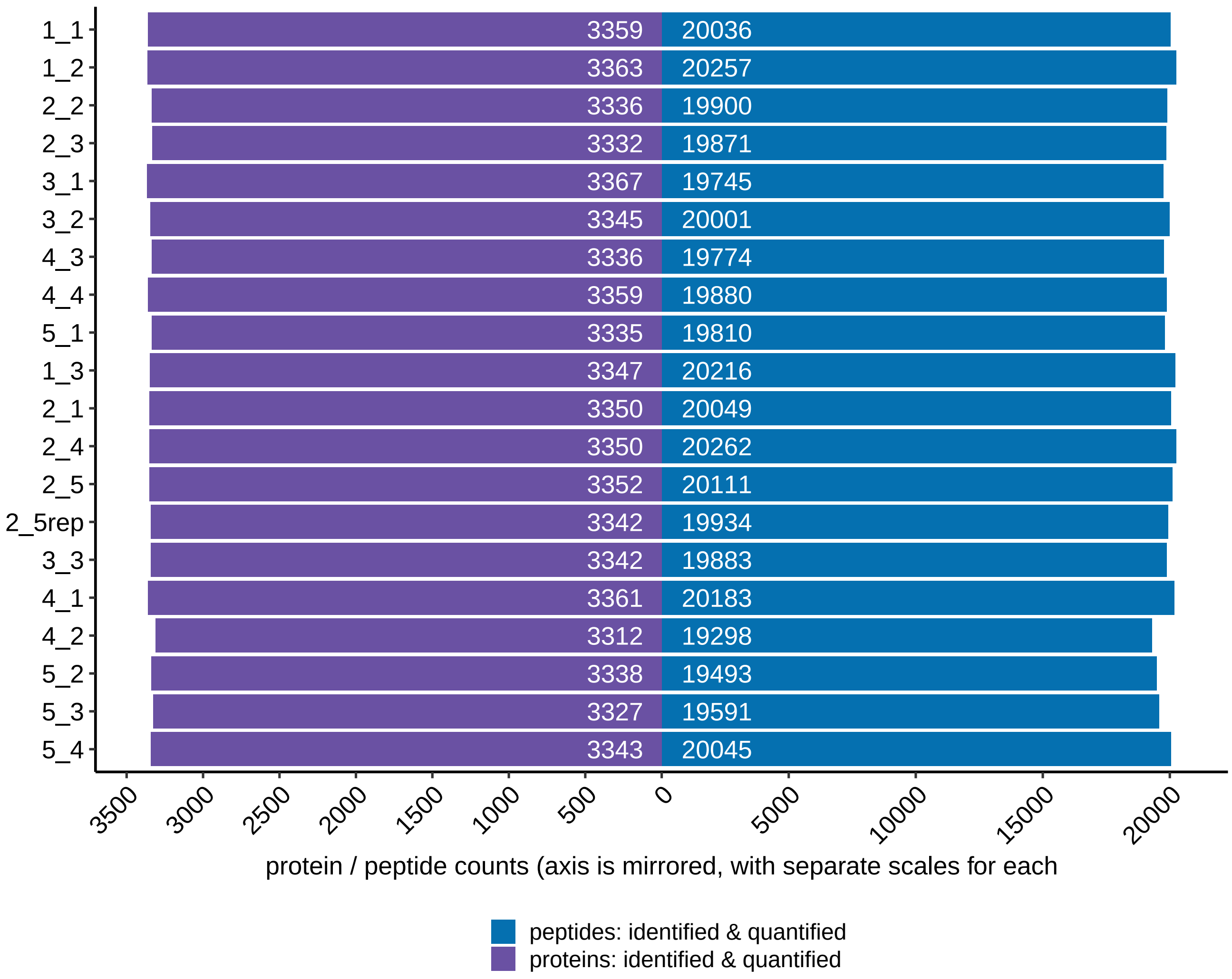

##### 1.1.1 color-coding sample metadata

The number of detected peptides in a sample, as compared to other samples within a dataset, can be used as a measure for sample quality. By color-coding individual samples for metadata that you provided as input (eg; experiment batch, sample handling order, gel lanes, etc.) we can visually inspect if these relate to the rate of successful peptide detection.

Each row in this figure shows all samples as a dot, each color-coded by the respective property shown on the y-axis, with minor vertical jitter for visual clarity. The section after this figure will further expand this summary figure into a detailed figure for each sample property.

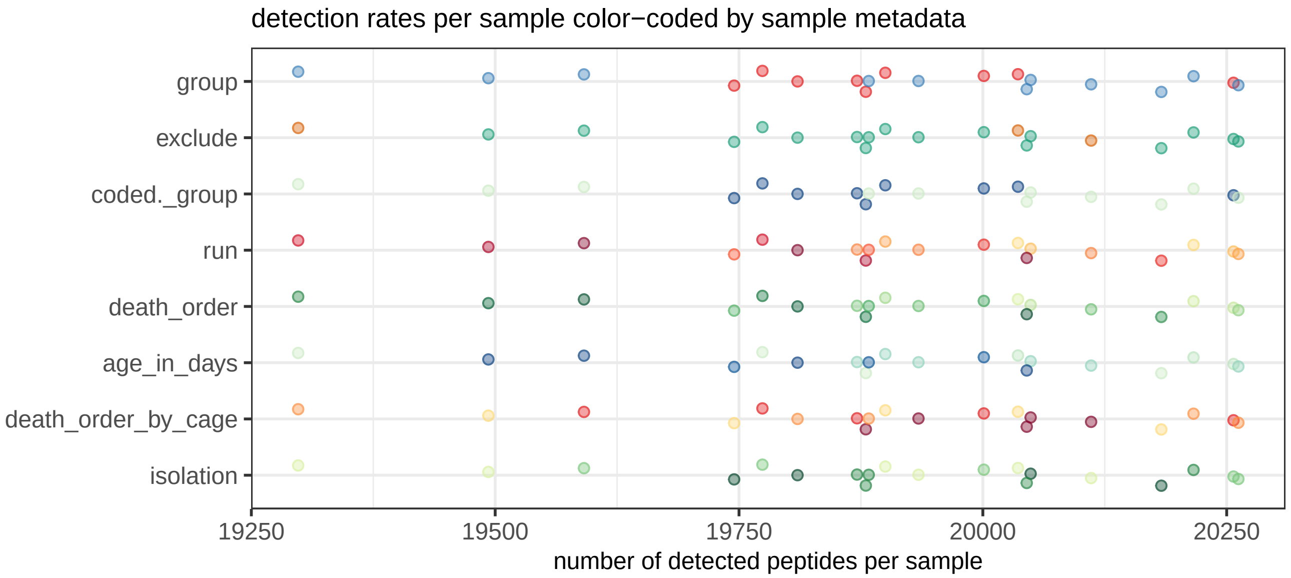

###### color-coding sample metadata, expanded

To further detail each sample property, we now split each row in the above figure into separate plots. Here, the rows detail the set of unique values for some sample metadata and the color-coded dots represent the samples annotated respectively. The vertical line indicates the median value for each row. Colors are consistent with the above plot.

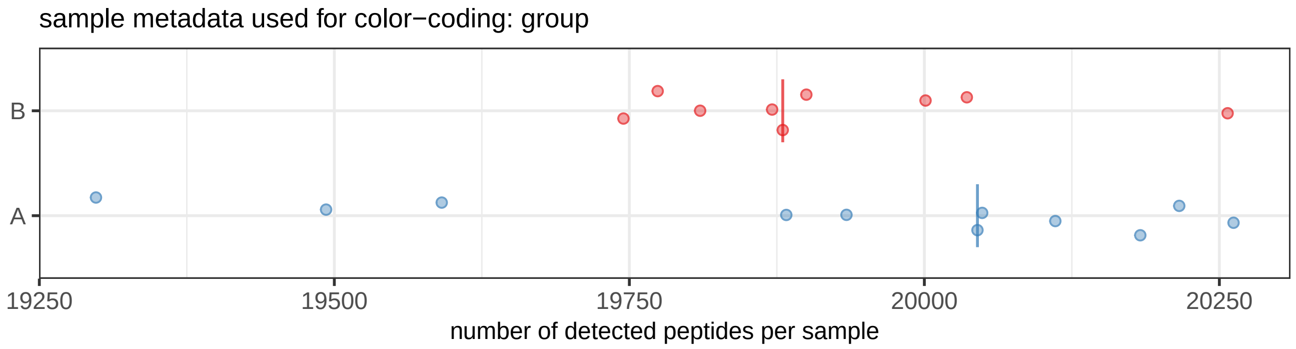

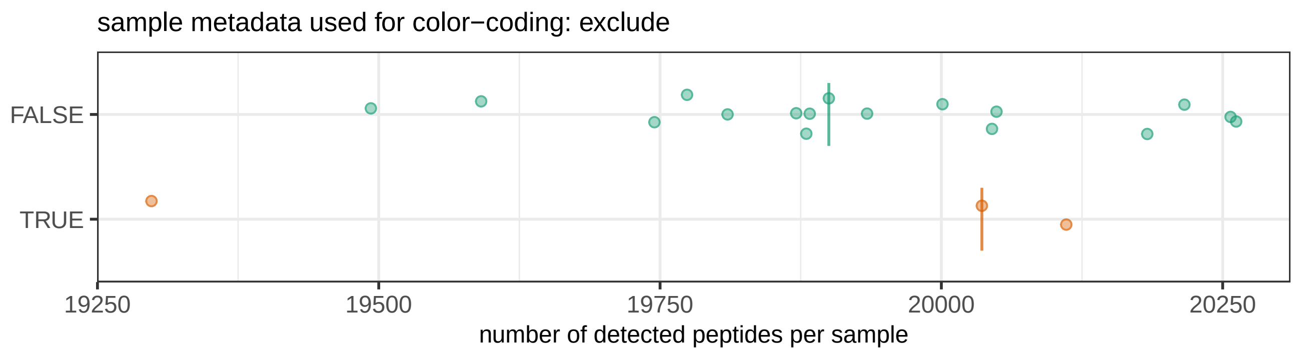
sample metadata used for color−coding: exclude
FALSE
TRUE
19250
19500
19750
20000
20250
number of detected peptides per sample

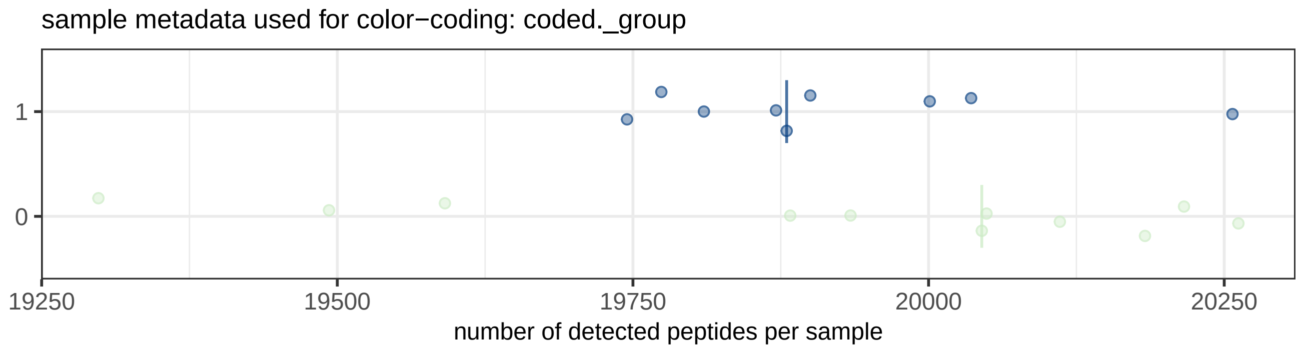
sample metadata used for color−coding: coded._group
1
0
19250
19500
19750
20000
20250
number of detected peptides per sample

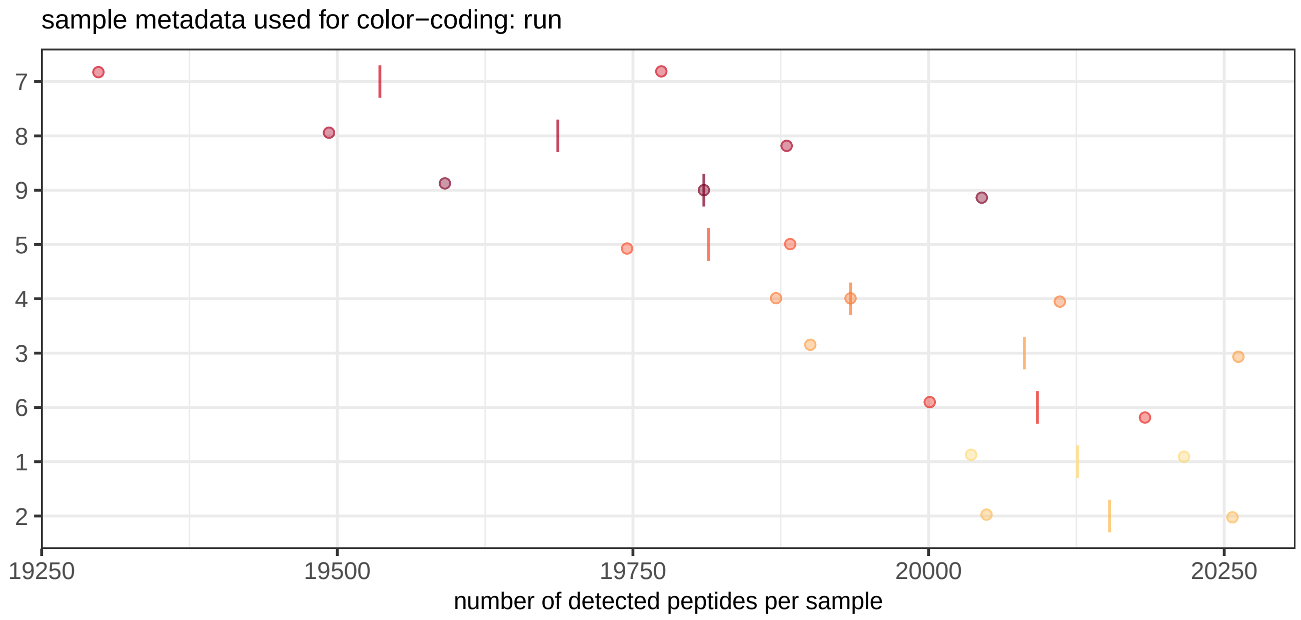
sample metadata used for color−coding: run
7
8
9
5
4
3
6
1
2
19250
19500
19750
20000
20250
number of detected peptides per sample

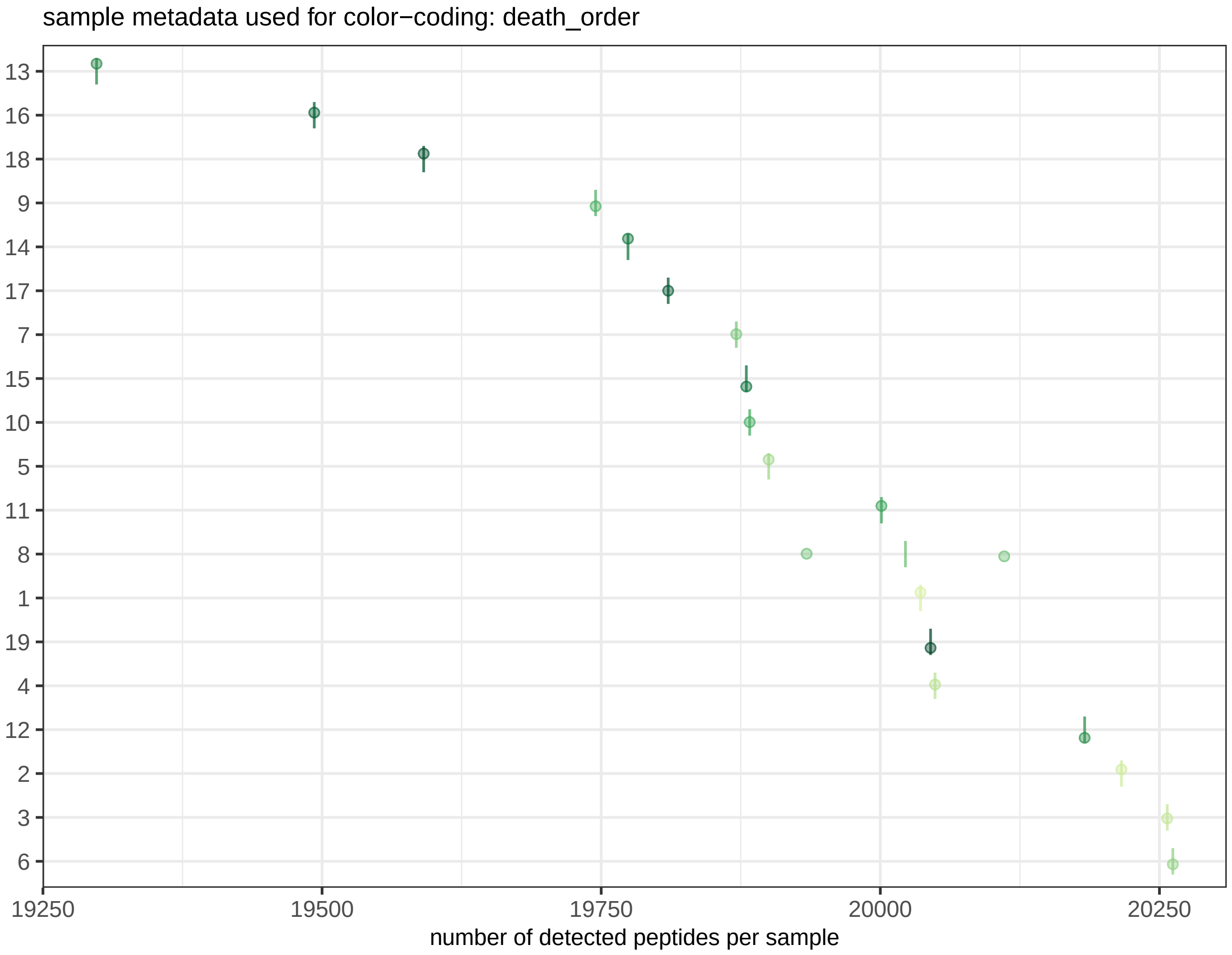

sample metadata used for color−coding: death_order
13
16
18
9
14
17
7
15
10
5
11
8
1
19
4
12
2
3
6
19250
19500
19750
20000
20250
number of detected peptides per sample

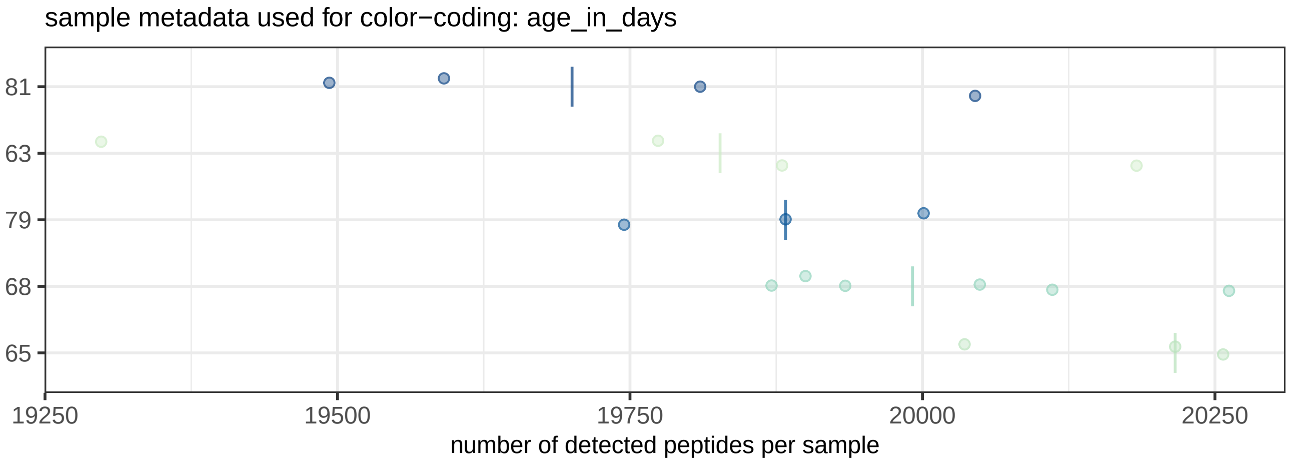

sample metadata used for color−coding: age_in_days
81
63
79
68
65
19250
19500
19750
20000
20250
number of detected peptides per sample

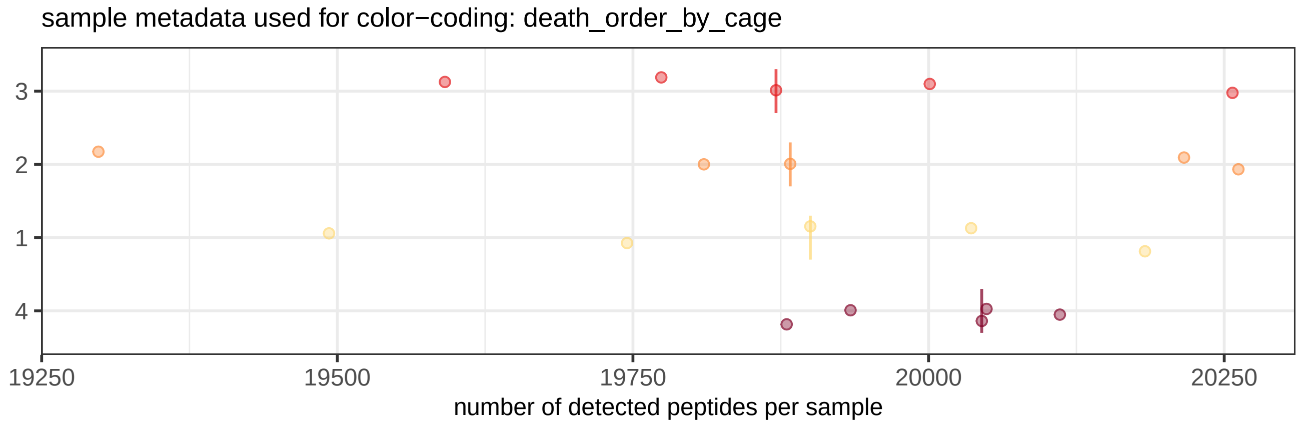
sample metadata used for color−coding: death_order_by_cage
3
2
1
4
19250
19500
19750
20000
20250
number of detected peptides per sample

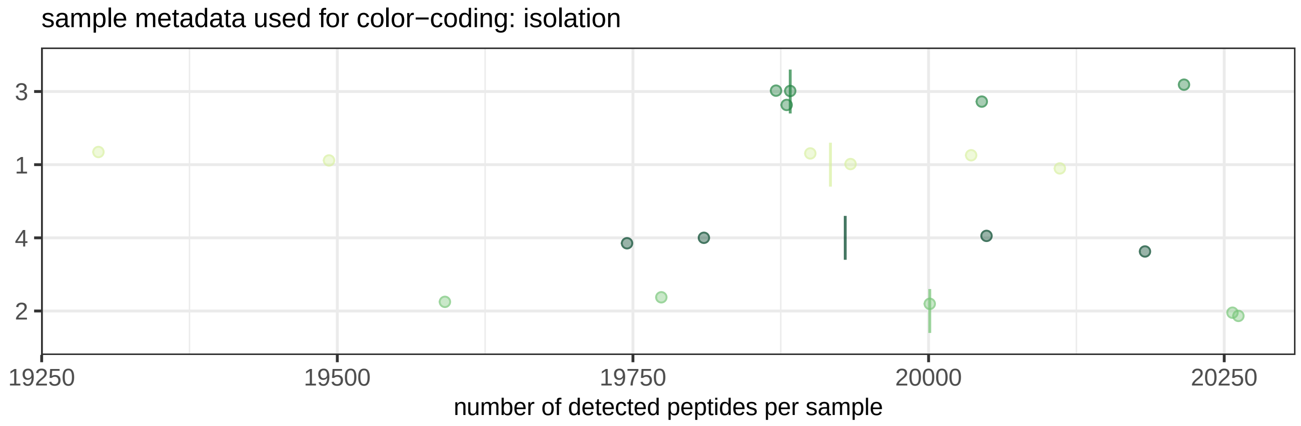
sample metadata used for color−coding: isolation
3
1
4
2
19250
19500
19750
20000
20250
number of detected peptides per sample

##### 1.2 data completeness

Visualize how many peptides are consistently identified in multiple samples. The first figure summarizes how common missing values are in the entire dataset, ideally many peptides are identified in 100% of samples and this curve slowly falls of. The second figure shows for each sample whether its peptides are also present in other samples in the dataset or whether these are unique to a (minor) subset of samples. You can use this mark of experimental consistency to compare datasets generated by similar protocols and mass-spec acquisition.

###### 1.2.1 cumulative distribution

The number of peptides passing some 'data completeness threshold' are shown at data points matching 90% and 50% of samples respectively. Samples flagged 'exclude' are not taken into account.

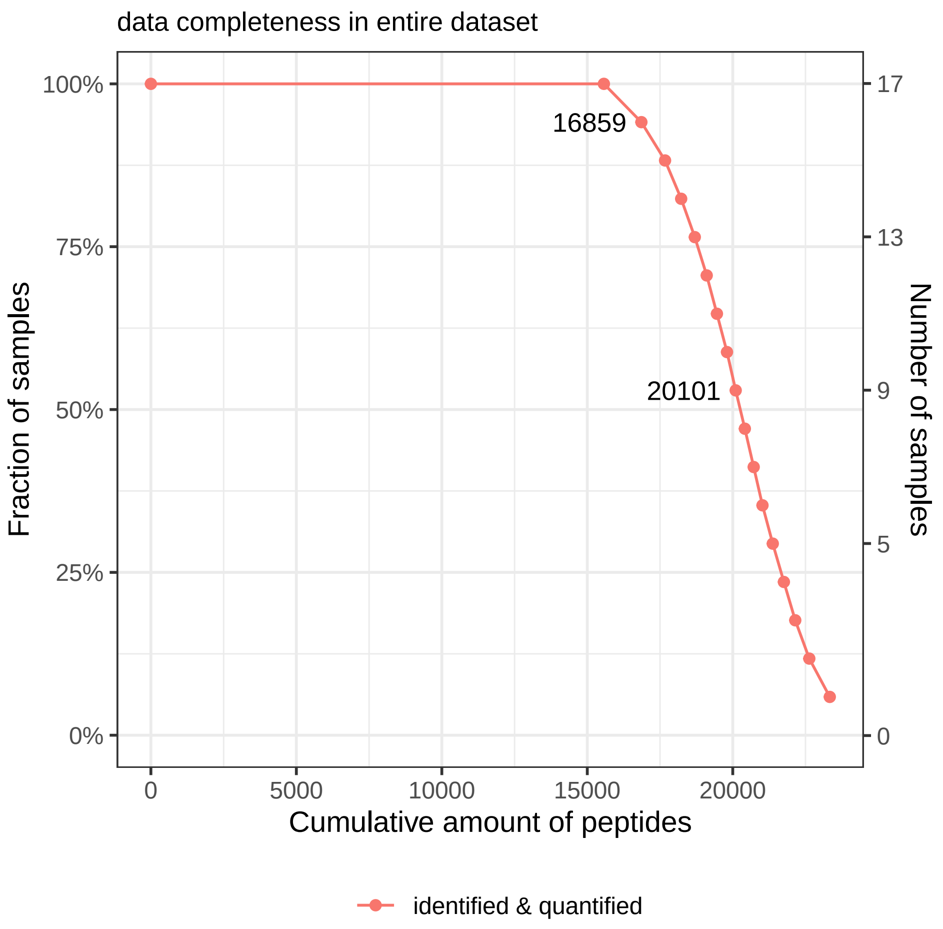

##### 1.2.2 peptide detection frequency

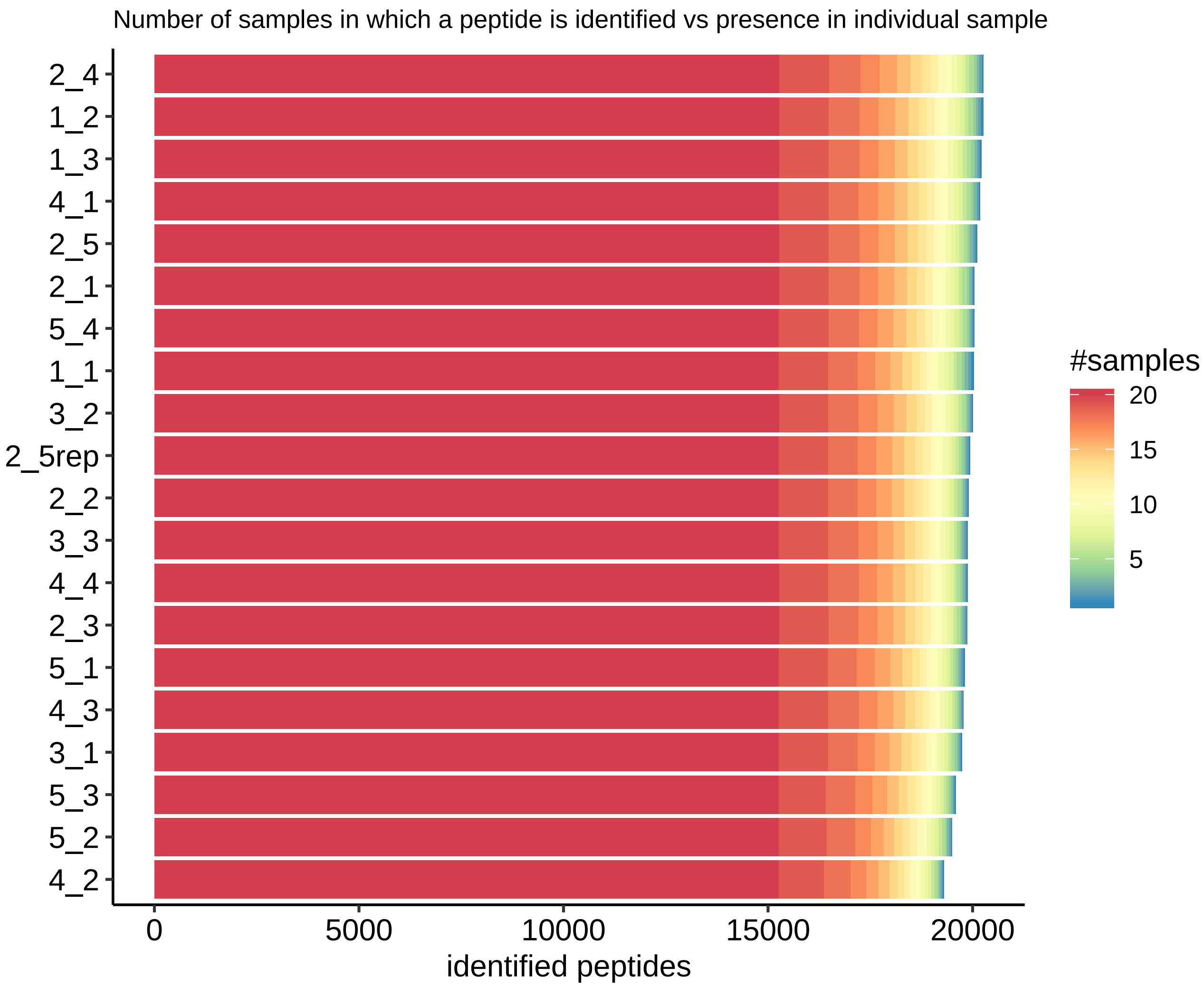

#### 1.3 abundance distributions

The figures in this subsection are used to identify unexpected mass-spec sensitivity or sample loading differencess. Peptide data is shown as provided in input files, so peptide filtering nor intensity normalization has been applied yet (for proper QC, make sure the software that generated the input data did not apply normalization prior). If the dataset is DDA, match-between-runs (MBR) peptides are included in these distributions whereas for DIA only 'detected' peptides (based on confidence score threshold) are included.

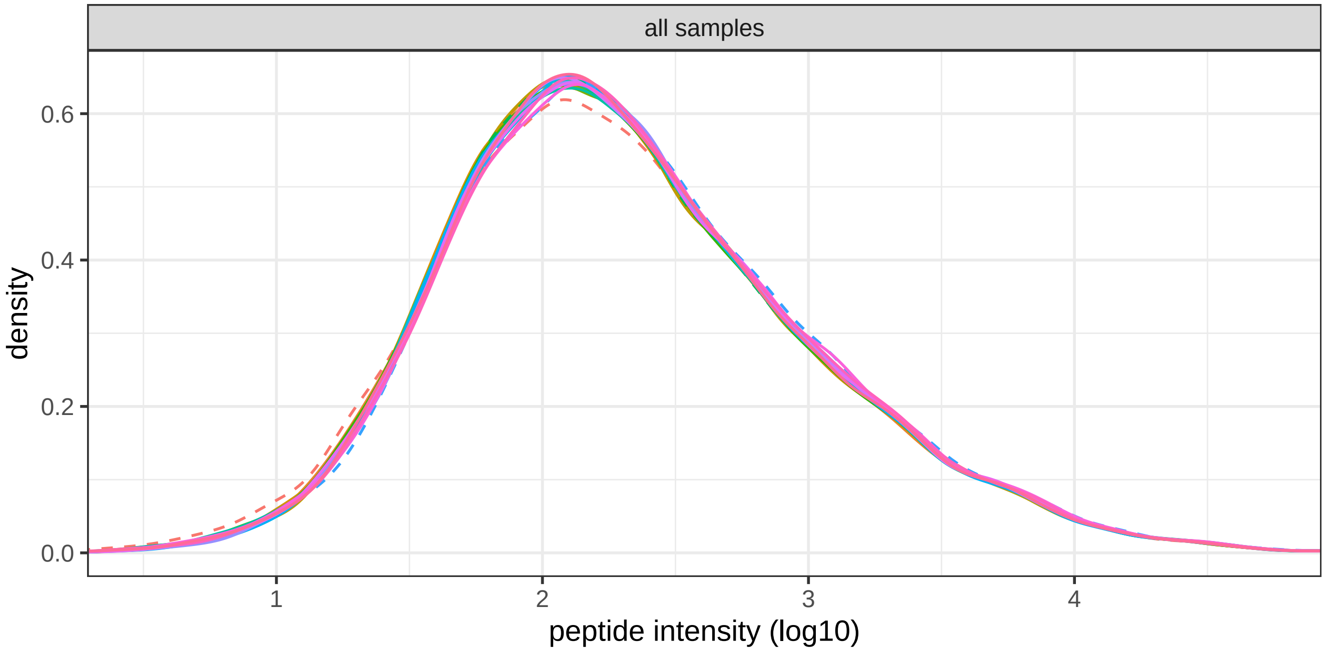

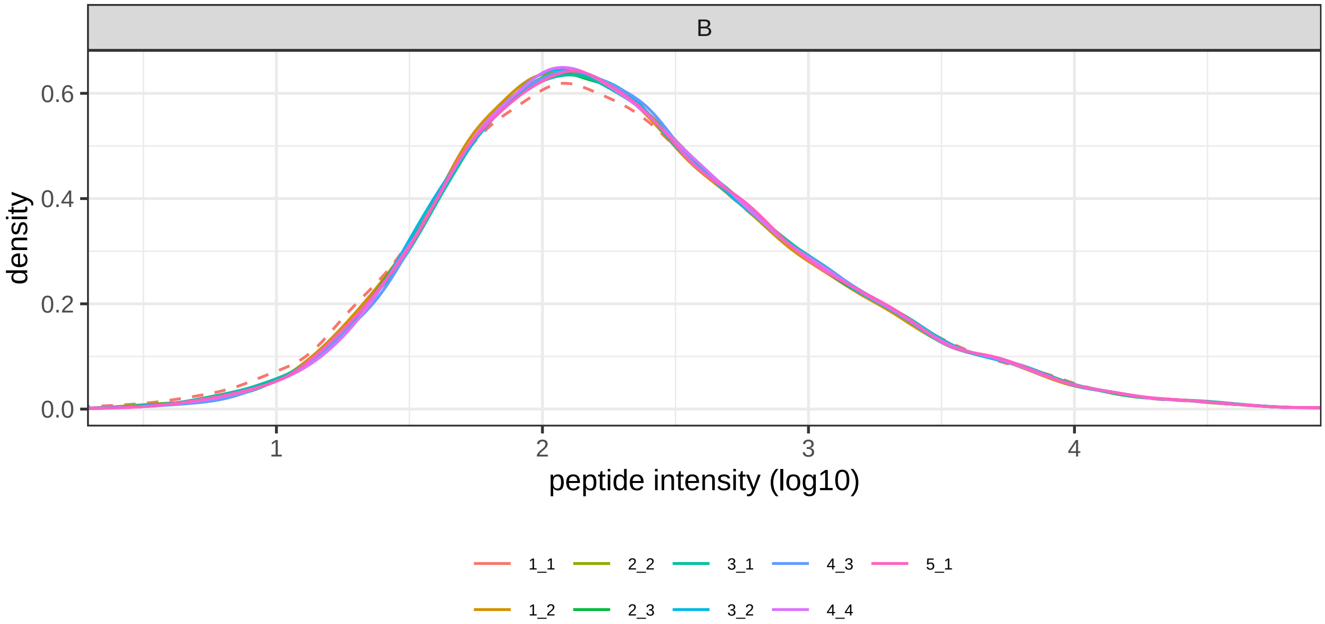

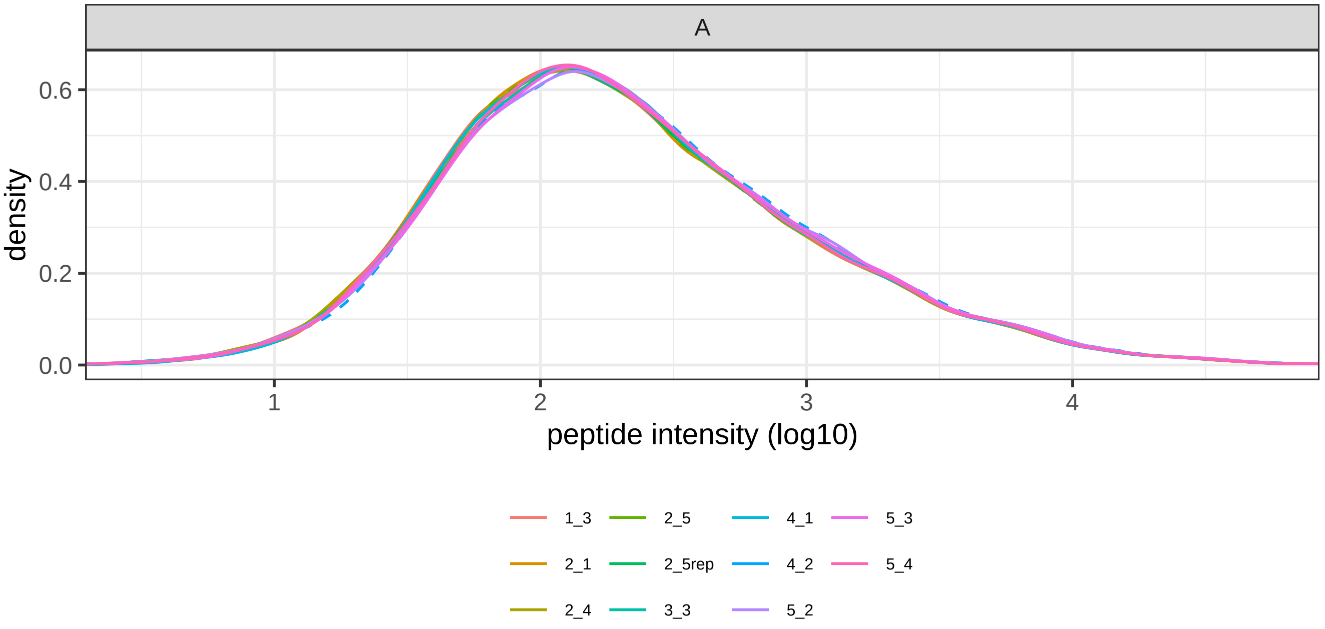
A
density
0.6
0.4
0.2
0.0
1
2
3
4
peptide intensity (log10)
1_3
2_1
2_4
2_5
2_5rep
3_3
4_1
4_2
5_2
5_3
5_4

##### 1.4 retention time

These figures allow you to identify potential problems during elution, such as a temporarily blocking column, failing ionization spray or decreasing sensitivity over time. For each sample, all peptides that are also observed in a replicate (such that there is a point of reference available) are visualized.

###### 1.4.1 retention time distributions

The density of the number of peptides eluting at each point in time. This figure presents a global view of all samples that allows for the identification of outlier samples that follow distinct elution patterns. The following section shows details for each sample. Samples marked as 'exclude' in the provided sample metadata table are visualized as dashed lines.

progress: RT plots: preparing data took 5 seconds progress: RT plots: creating plots took 2 seconds

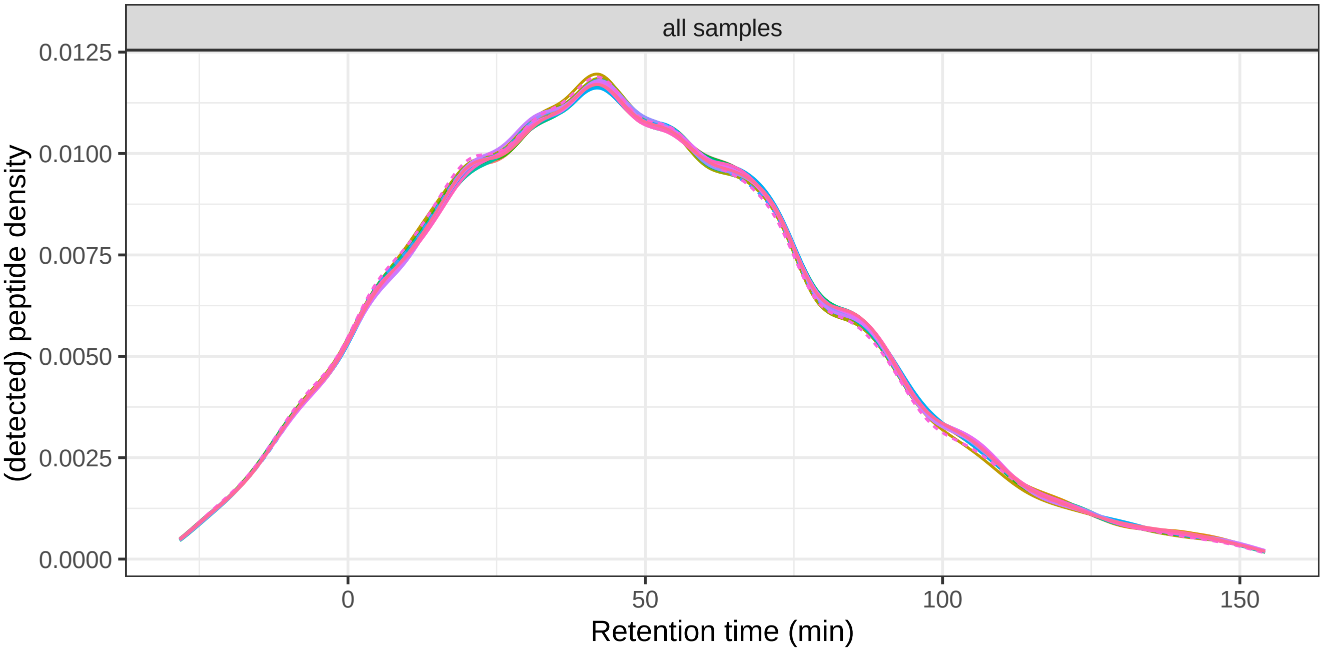

###### 1.4.2 retention time local effects

To investigate how each measurement differs from others, we visualize each sample as a 4 panel figure. First, the data is binned across the retention time dimension (x-axis). If a samples was marked as 'exclude' in the provided sample metadata, this is indicated in the plot title.

The top panel shows the number of peptides, as recognized by the software that generated input for this pipeline, over time (black line). For reference, the grey line shows the median amount over all samples.

The middle panel indicates whether peptide retention times deviate from their median over all samples (blue line). The grey area depicts the 5% and 95% quantiles, respectively. The line width corresponds to the number of peptides eluting at that time (data from first panel). Analogously, the bottom panel show the deviation in peptide abundance as compared to the median over all samples (red line).

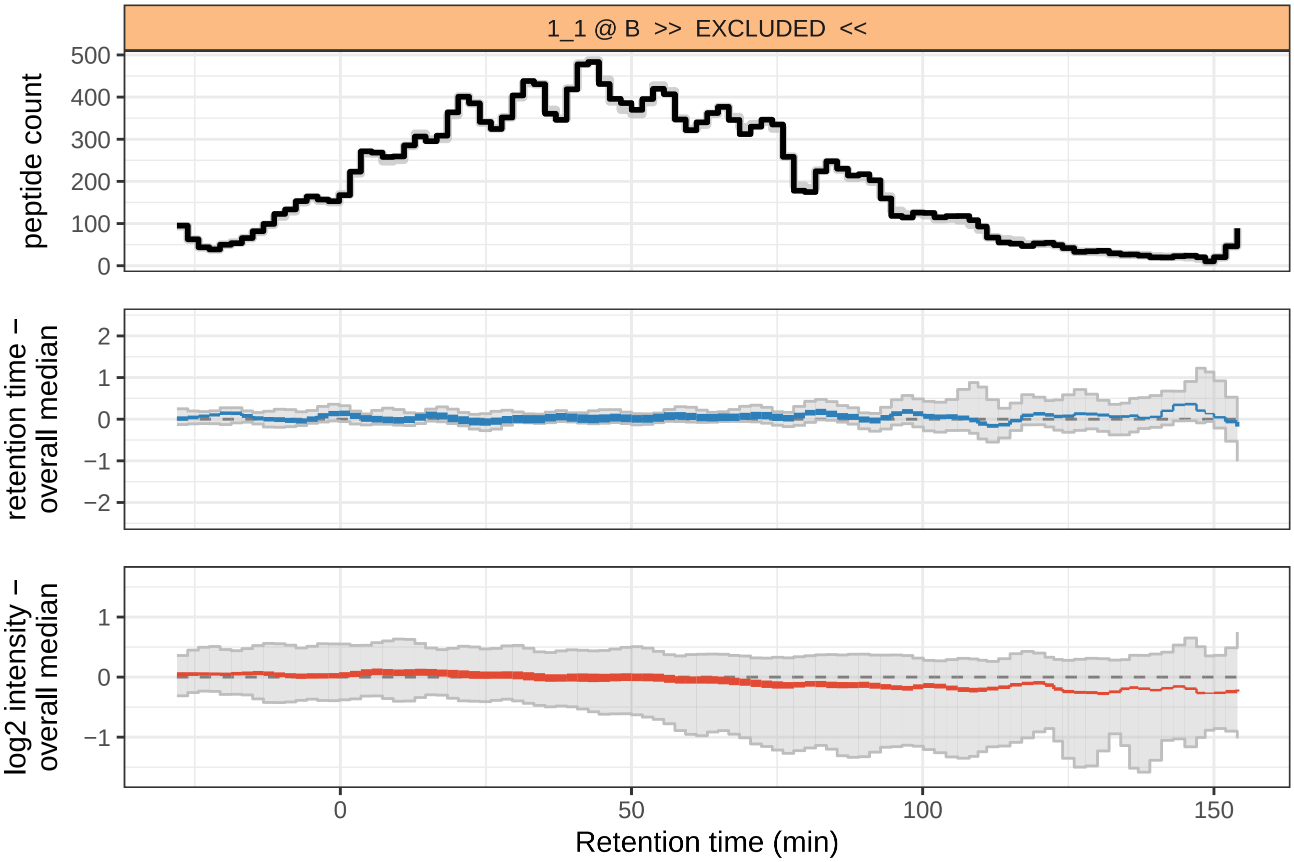
1_1 @ B >> EXCLUDED <<
peptide count
retention time – overall median
log2 intensity – overall median
Retention time (min)

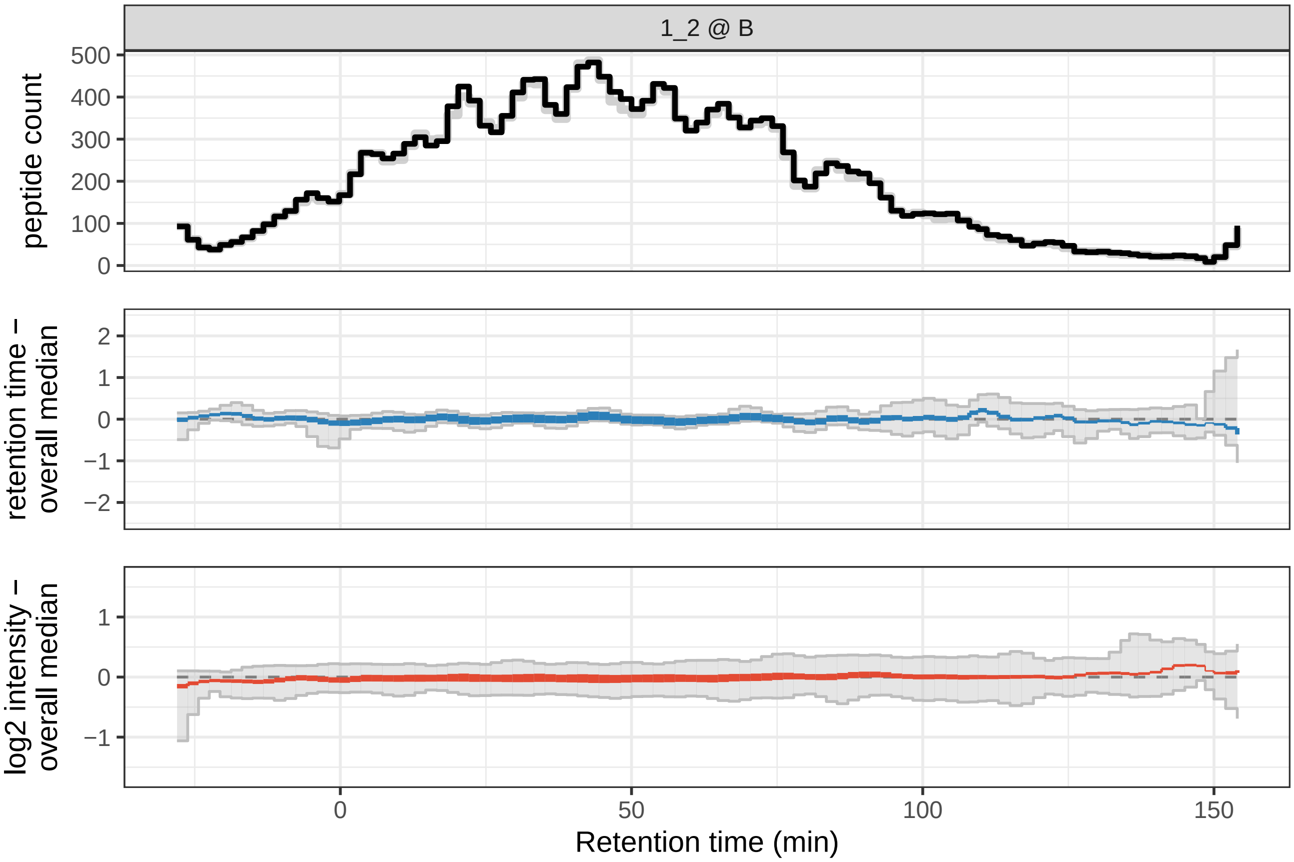
1_2 @ B
peptide count
retention time – overall median
log2 intensity – overall median
Retention time (min)

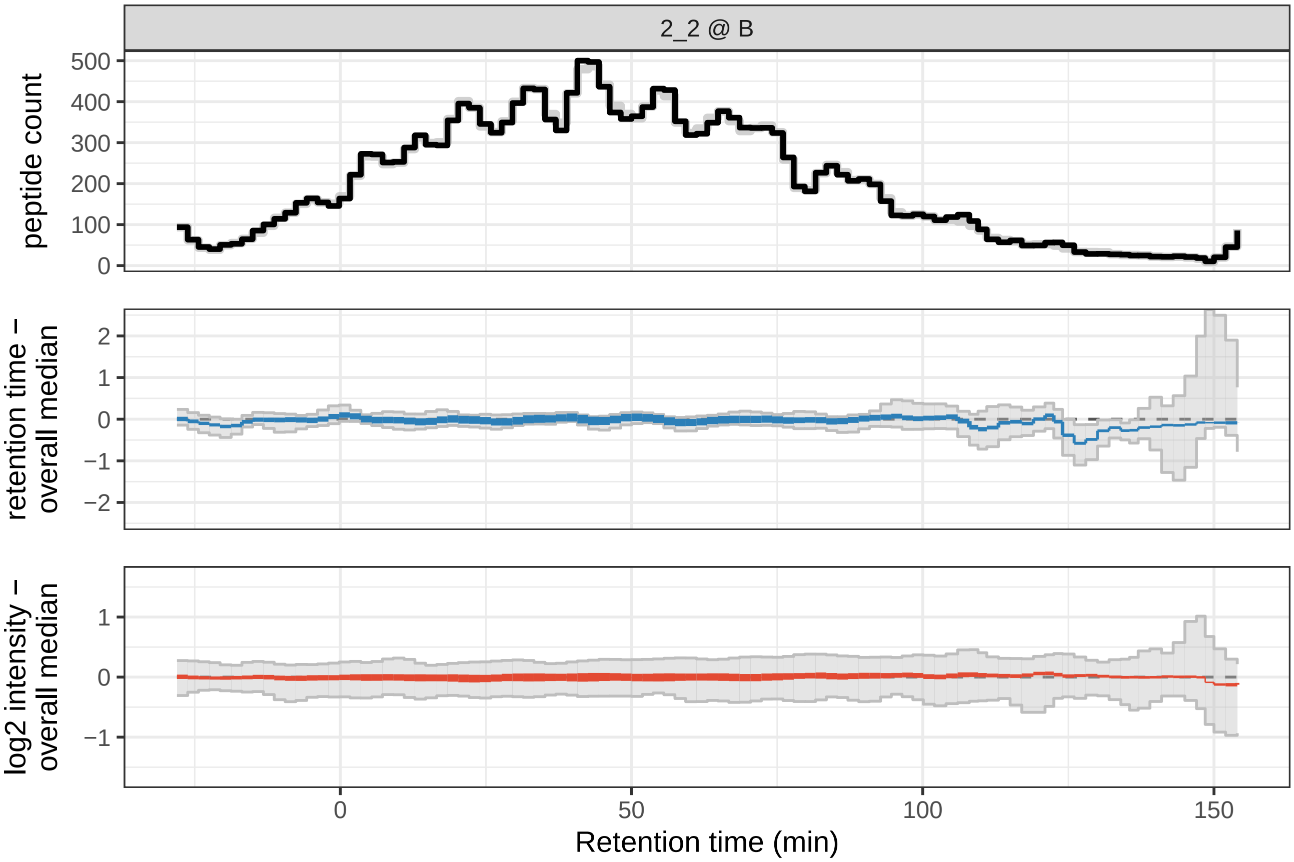

2_2 @ B
peptide count
500
400
300
200
100
0
retention time –
overall median
2
1
0
−1
−2
log2 intensity –
overall median
1
0
−1
0
50
100
150
Retention time (min)

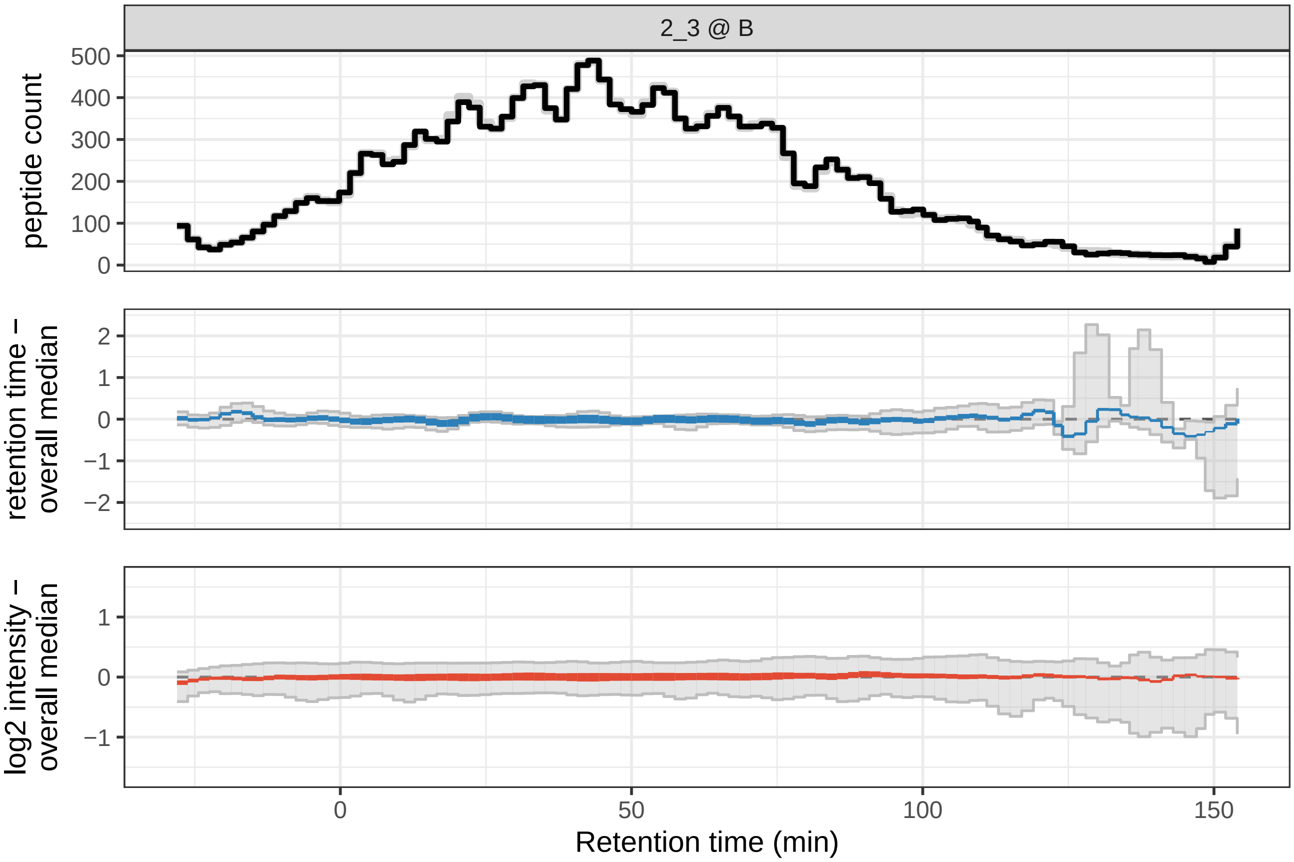

2_3 @ B
peptide count
500
400
300
200
100
0
retention time –
overall median
2
1
0
−1
−2
log2 intensity –
overall median
1
0
−1
0
50
100
150
Retention time (min)

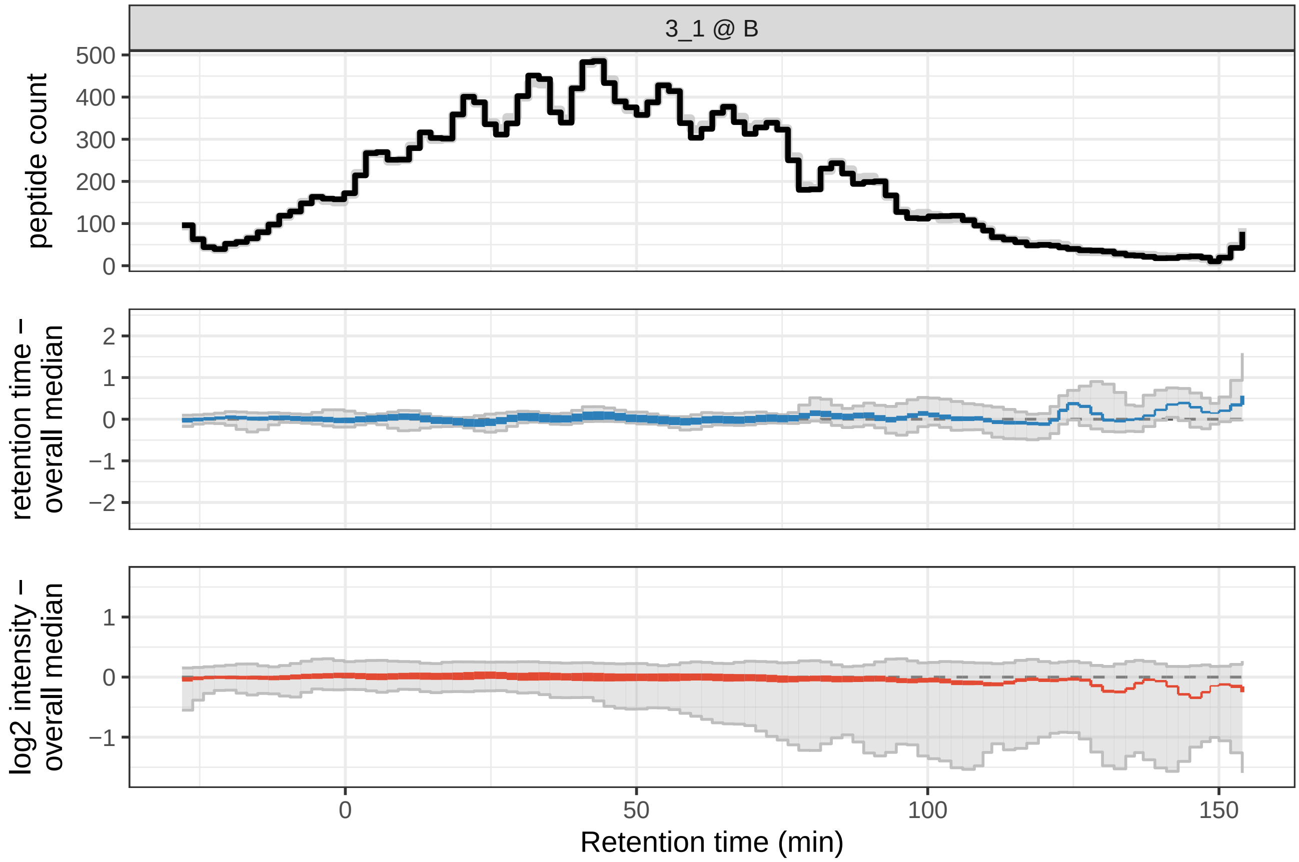
3_1 @ B
peptide count
500
400
300
200
100
0
retention time −
overall median
2
1
0
−1
−2
log2 intensity −
overall median
1
0
−1
0
50
100
150
Retention time (min)

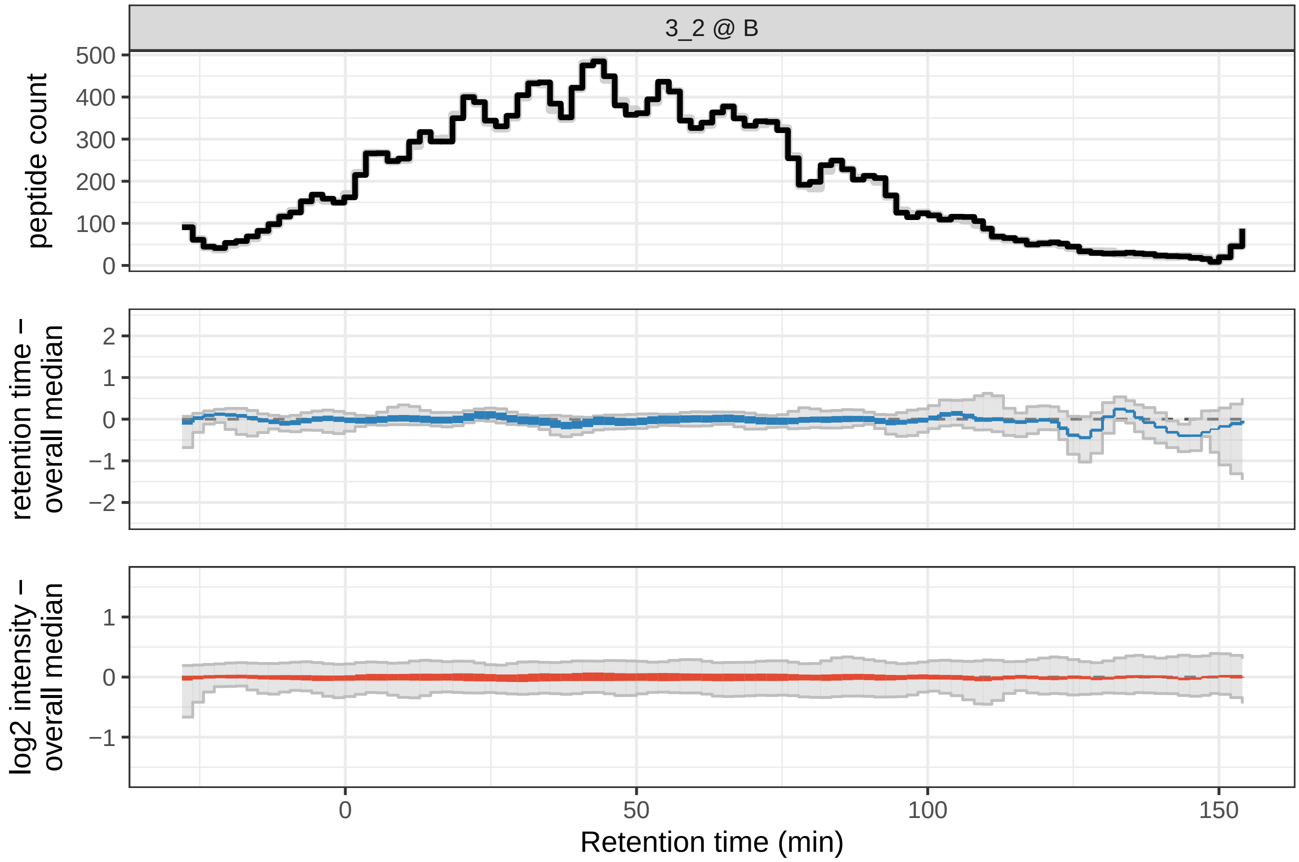
3_2 @ B
peptide count
500
400
300
200
100
0
retention time −
overall median
2
1
0
−1
−2
log2 intensity −
overall median
1
0
−1
0
50
100
150
Retention time (min)

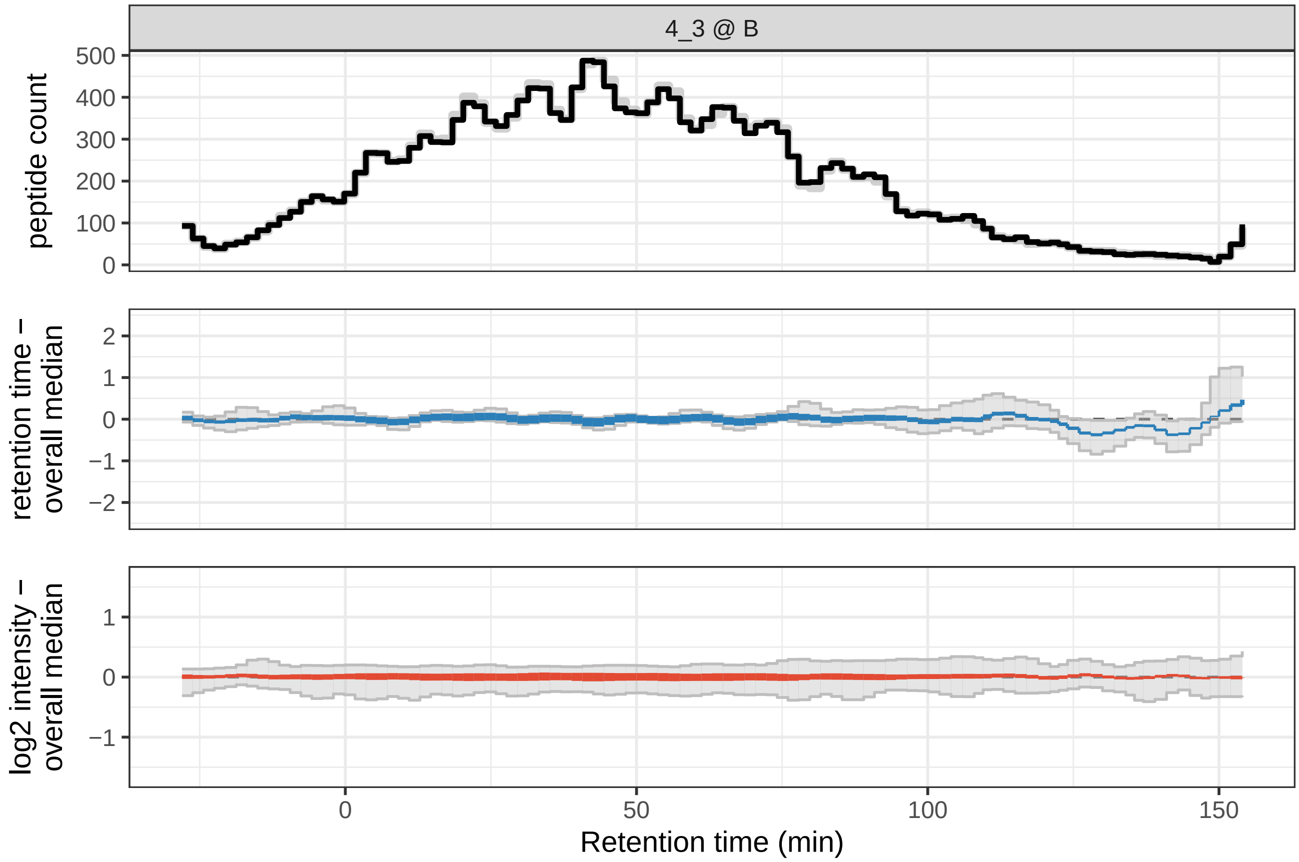
4_3 @ B
peptide count
500
400
300
200
100
0
retention time – overall median
2
1
0
−1
−2
log2 intensity – overall median
1
0
−1
0
50
100
150
Retention time (min)

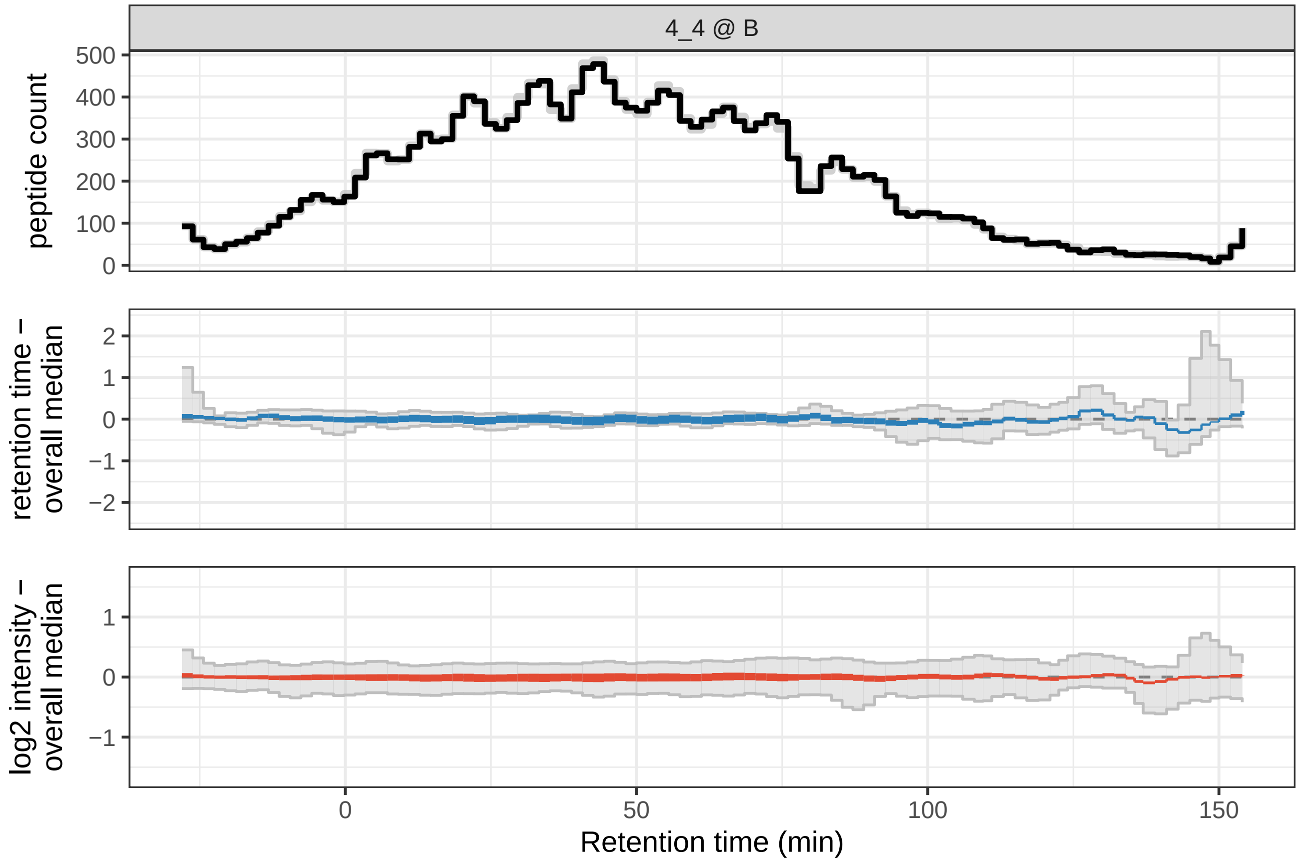
4_4 @ B
peptide count
500
400
300
200
100
0
retention time – overall median
2
1
0
−1
−2
log2 intensity – overall median
1
0
−1
0
50
100
150
Retention time (min)

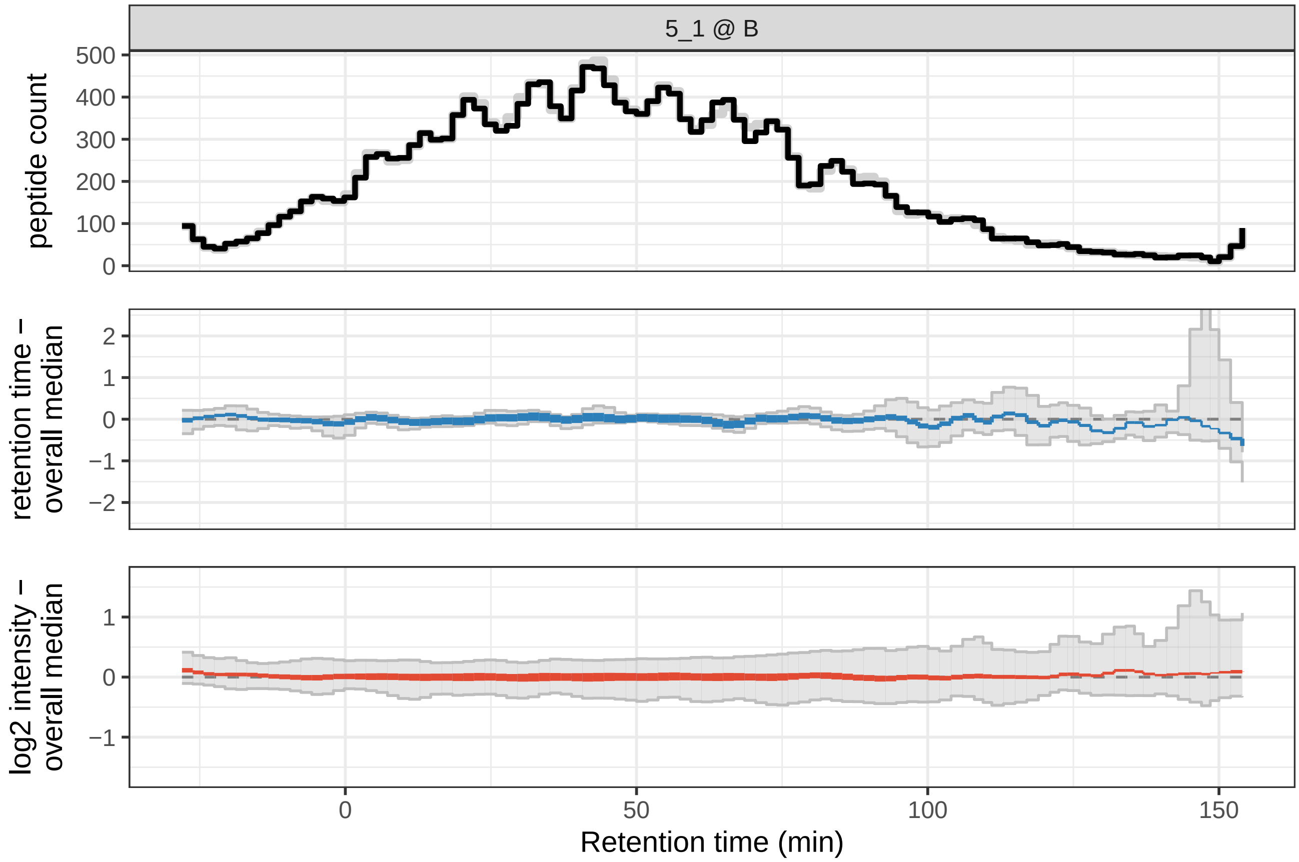

5_1 @ B
peptide count
500
400
300
200
100
0
retention time –
overall median
2
1
0
−1
−2
log2 intensity –
overall median
1
0
−1
0
50
100
150
Retention time (min)

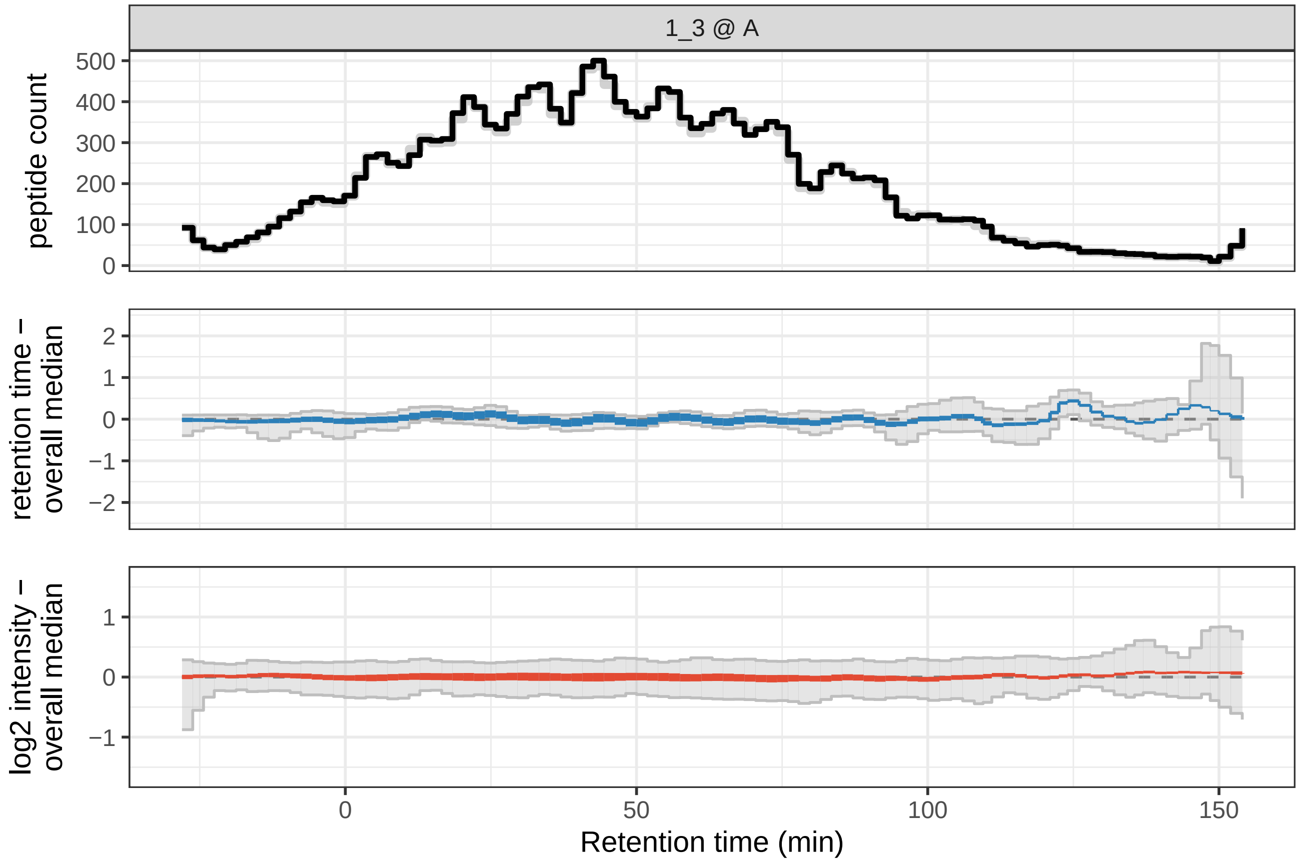

1_3 @ A
peptide count
500
400
300
200
100
0
retention time –
overall median
2
1
0
−1
−2
log2 intensity –
overall median
1
0
−1
0
50
100
150
Retention time (min)

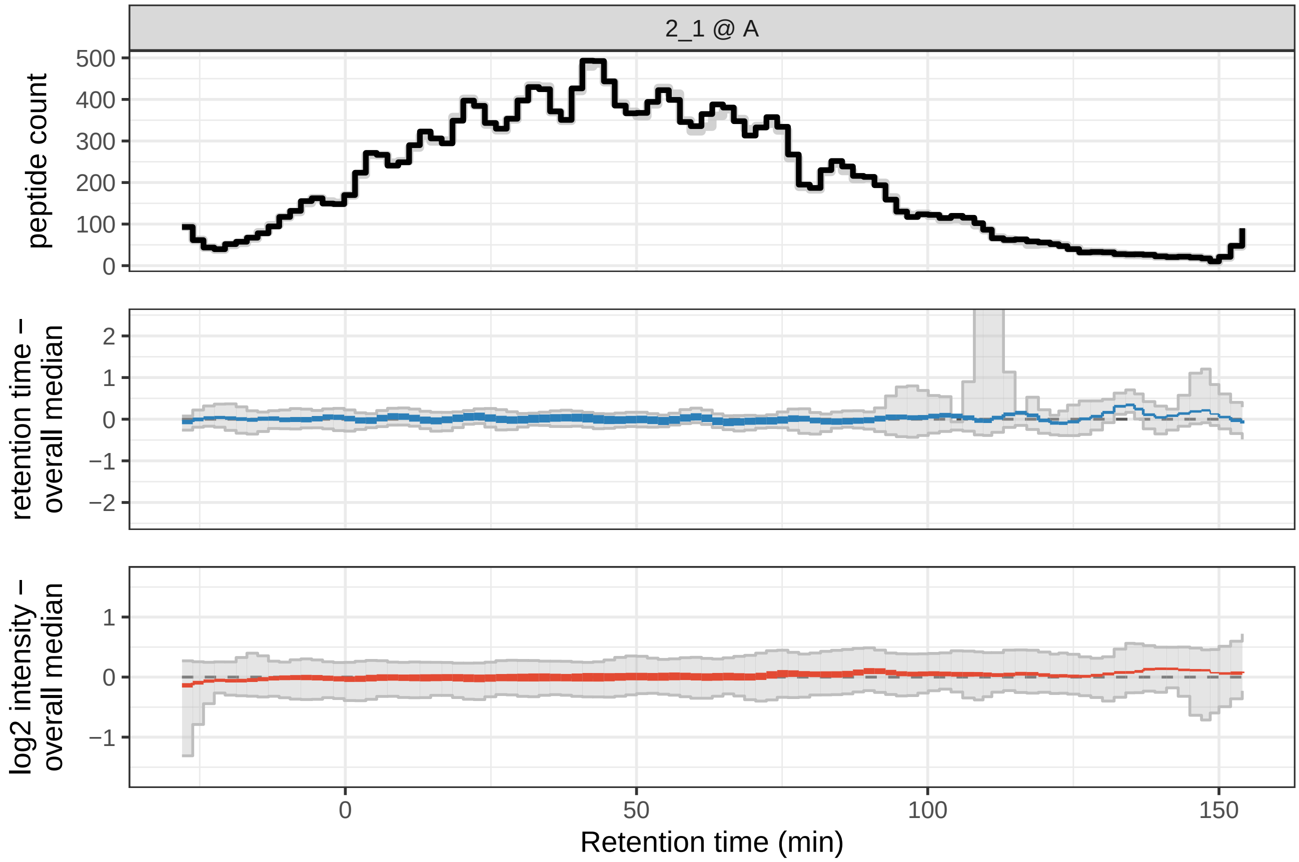
2_1 @ A
peptide count
retention time – overall median
log2 intensity – overall median
Retention time (min)

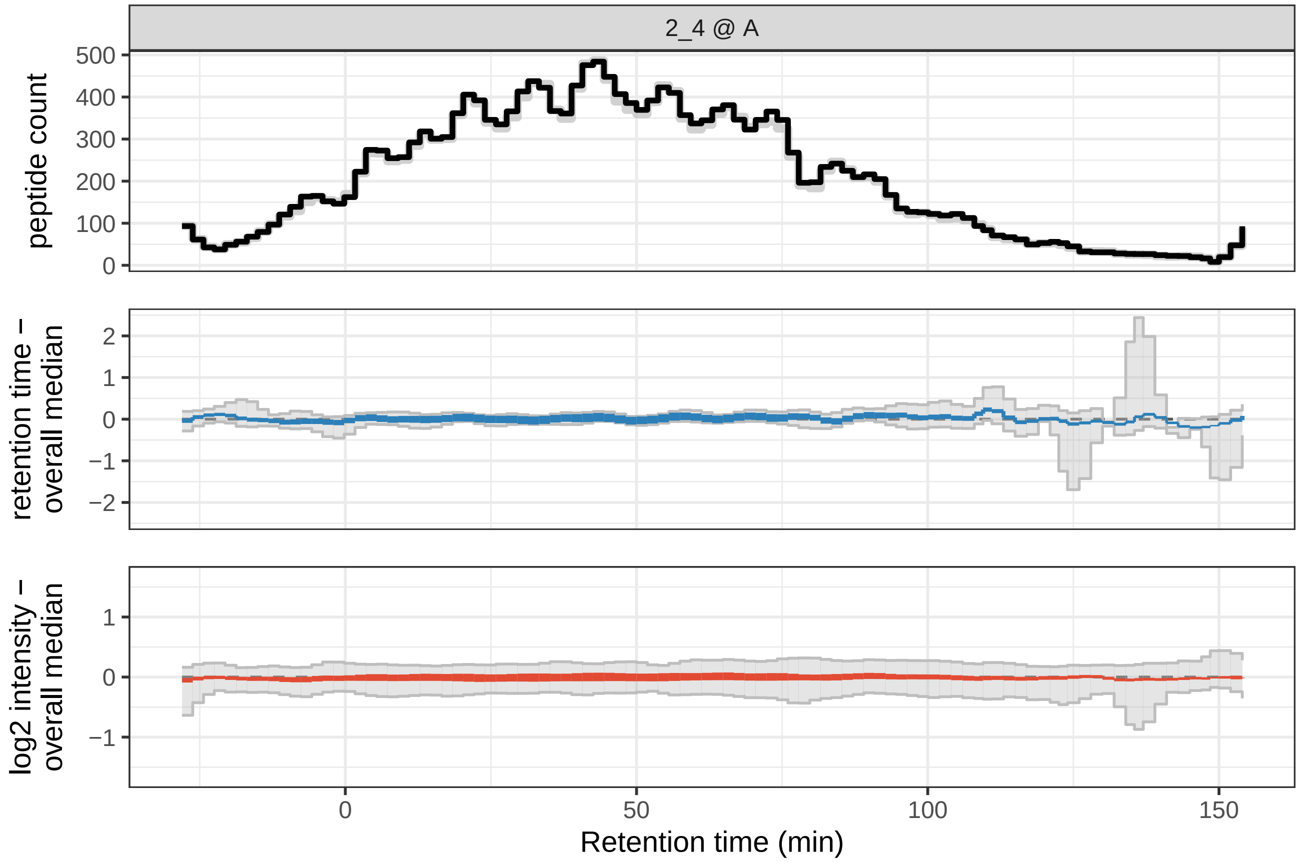
2_4 @ A
peptide count
retention time – overall median
log2 intensity – overall median
Retention time (min)

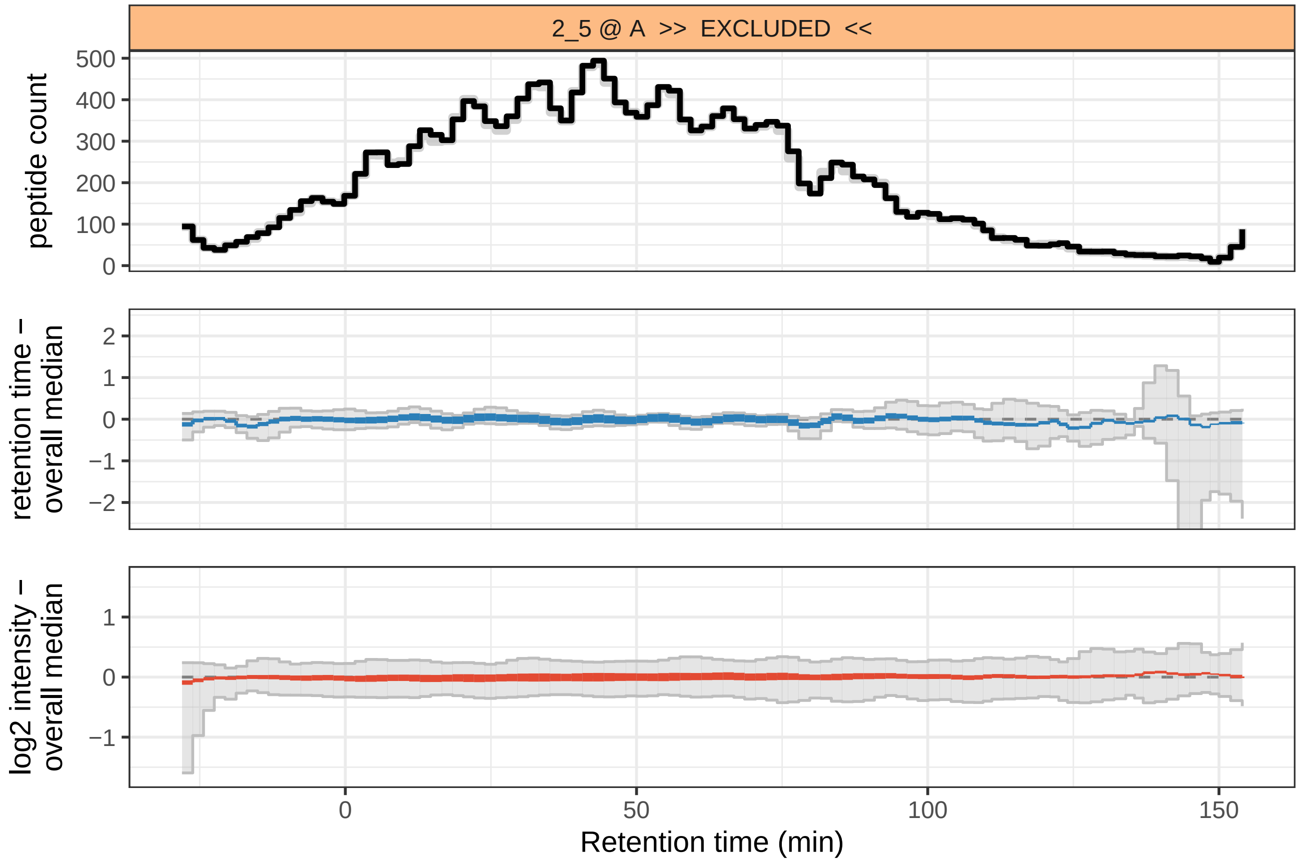

2_5 @ A >> EXCLUDED <<
peptide count
retention time – overall median
log2 intensity – overall median
Retention time (min)

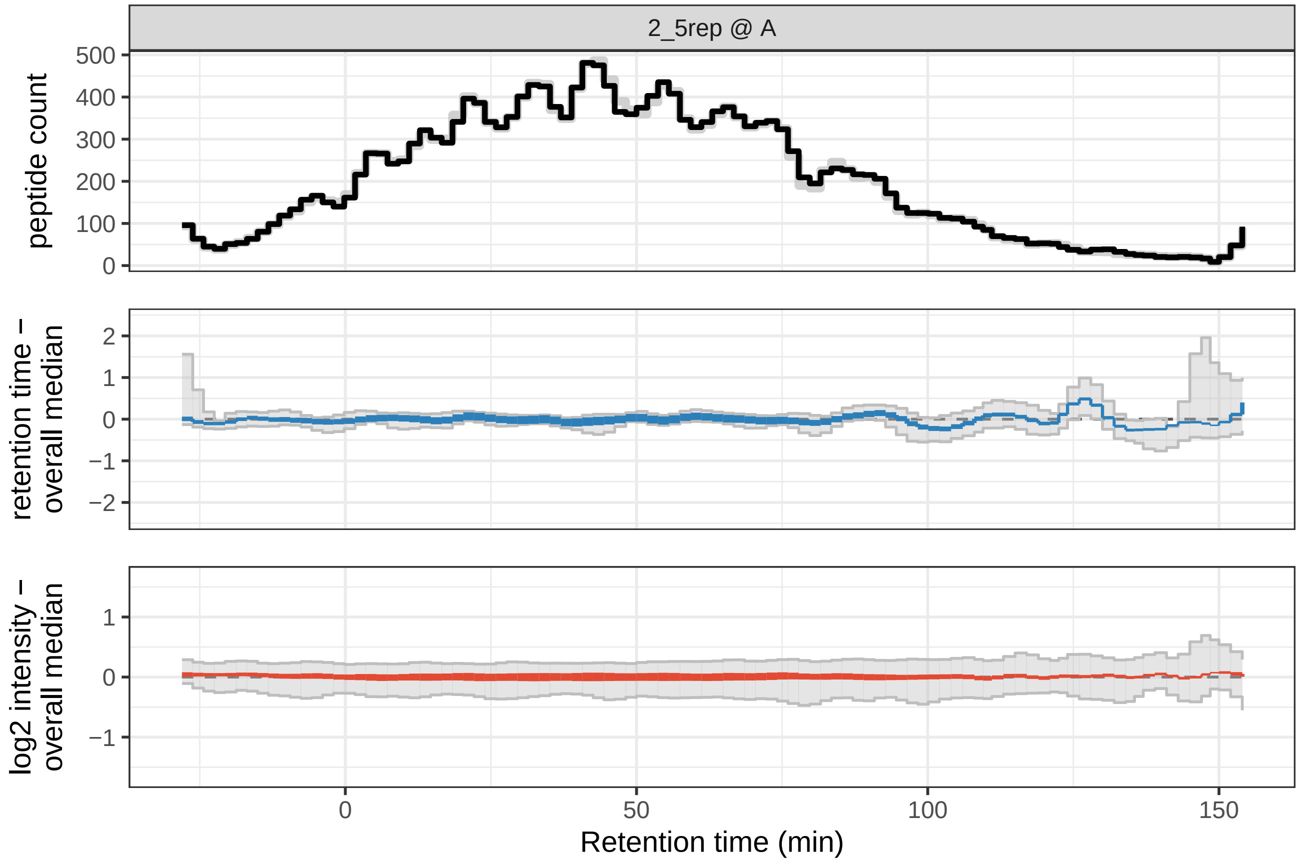

2_5rep @ A
peptide count
retention time – overall median
log2 intensity – overall median
Retention time (min)

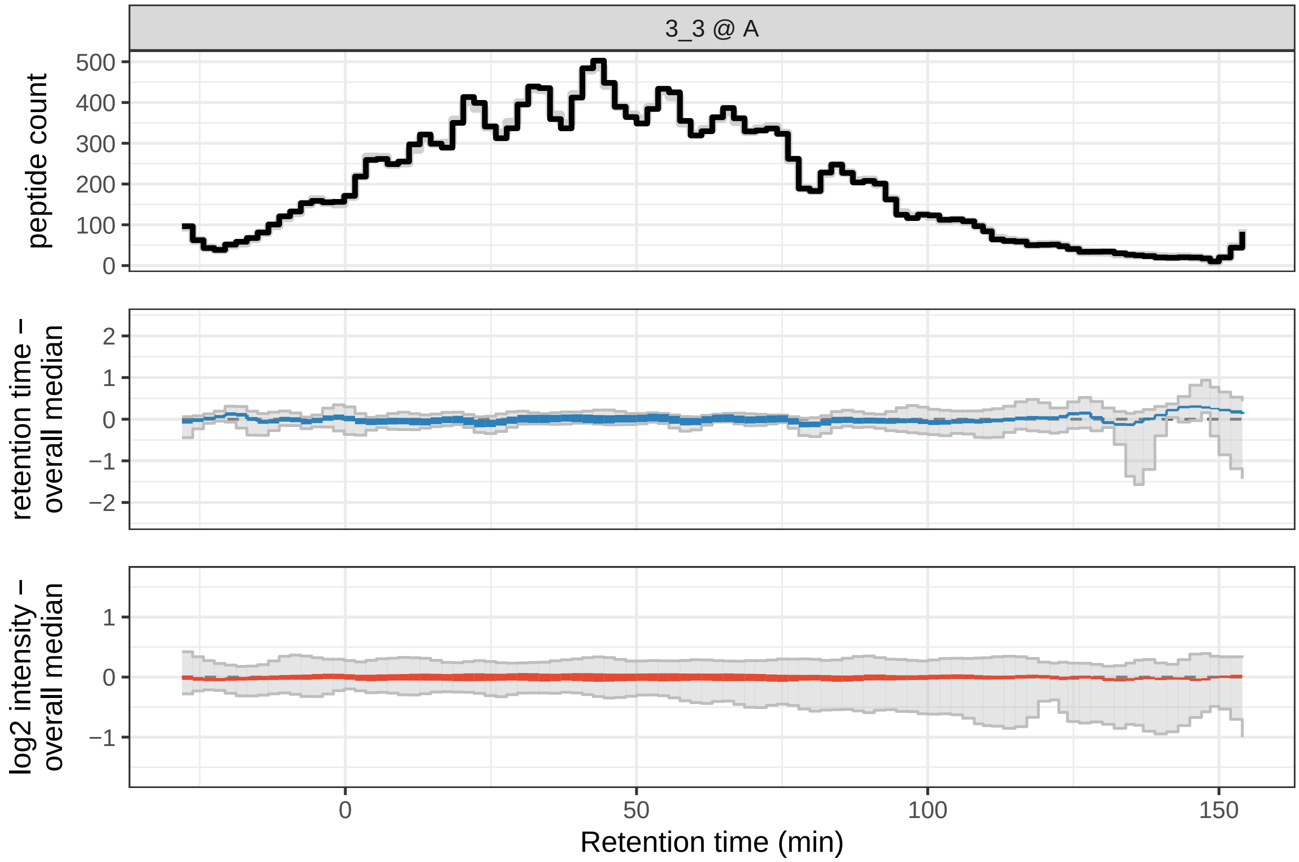
3_3 @ A
peptide count
500
400
300
200
100
0
retention time –
overall median
2
1
0
−1
−2
log2 intensity –
overall median
1
0
−1
0
50
100
150
Retention time (min)

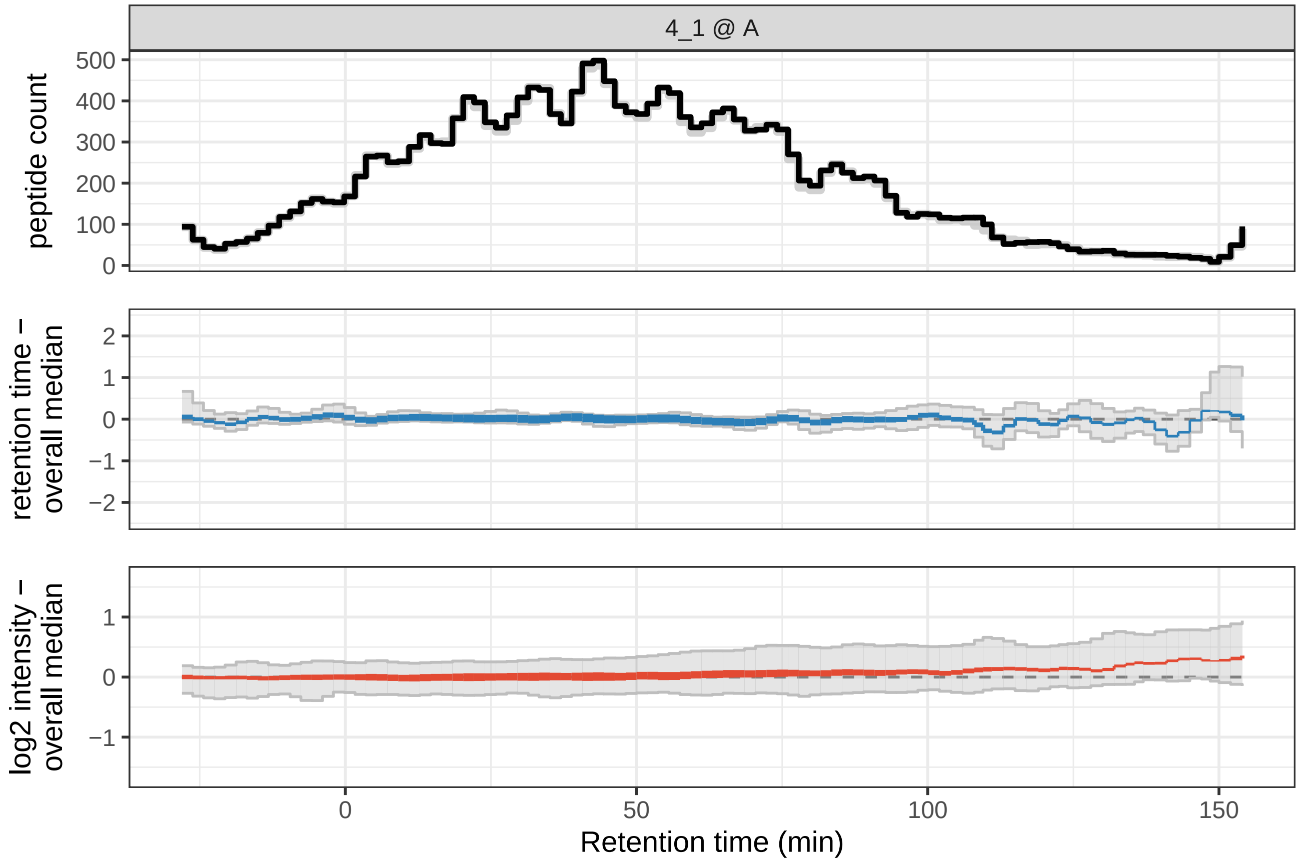
4_1 @ A
peptide count
500
400
300
200
100
0
retention time –
overall median
2
1
0
−1
−2
log2 intensity –
overall median
1
0
−1
0
50
100
150
Retention time (min)

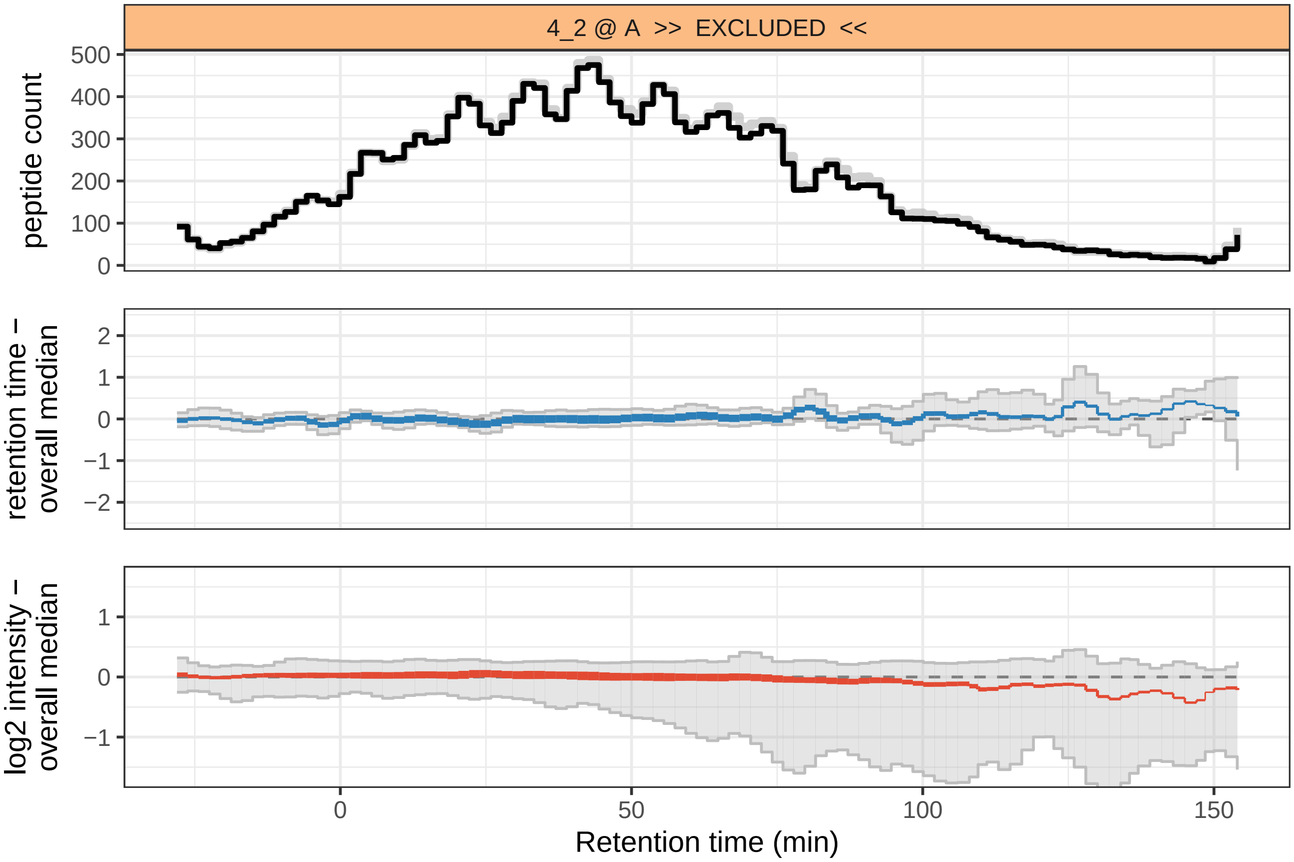
4_2 @ A >> EXCLUDED <<
peptide count
0
100
200
300
400
500
retention time –
overall median
−2
−1
0
1
2
log2 intensity –
overall median
−1
0
1
0
50
100
150
Retention time (min)

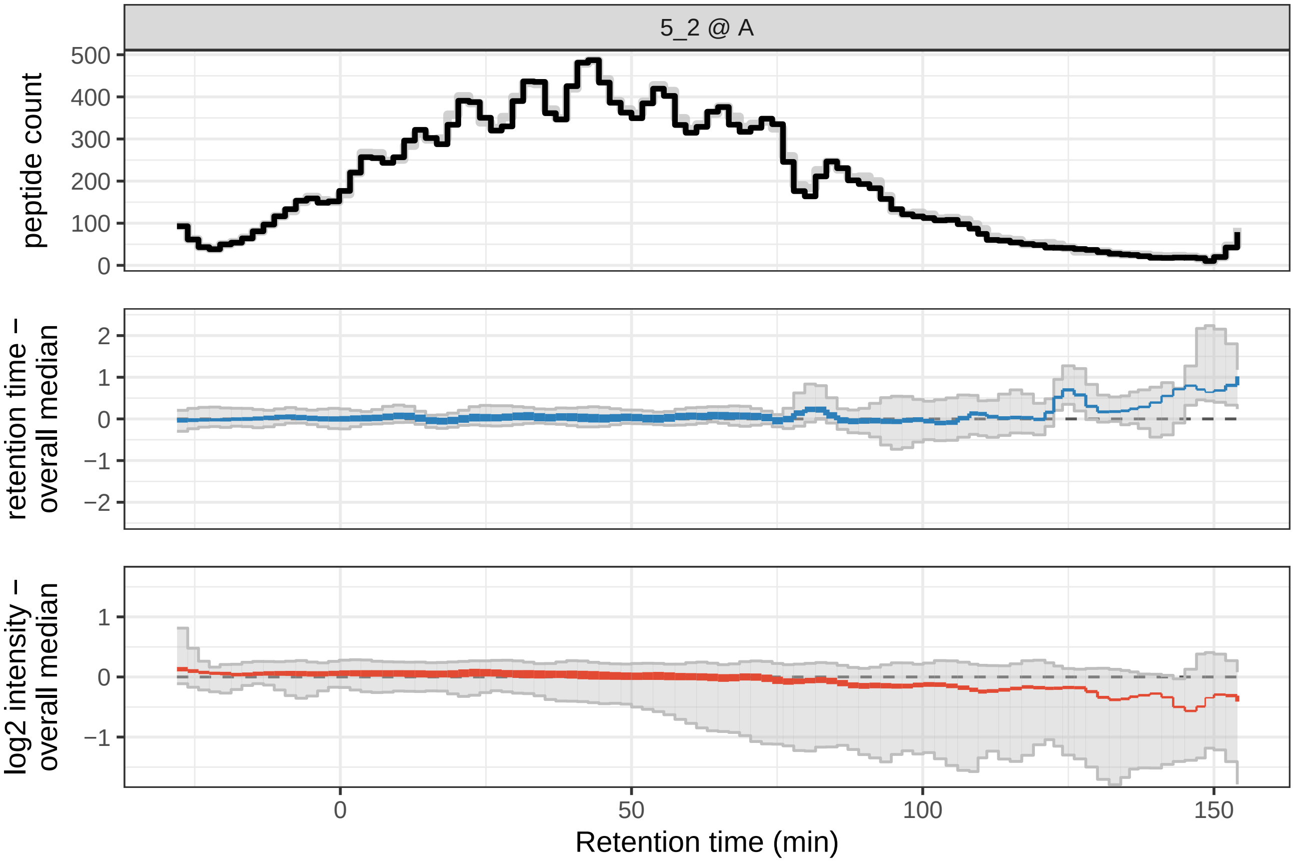
5_2 @ A
peptide count
0
100
200
300
400
500
retention time –
overall median
−2
−1
0
1
2
log2 intensity –
overall median
−1
0
1
0
50
100
150
Retention time (min)

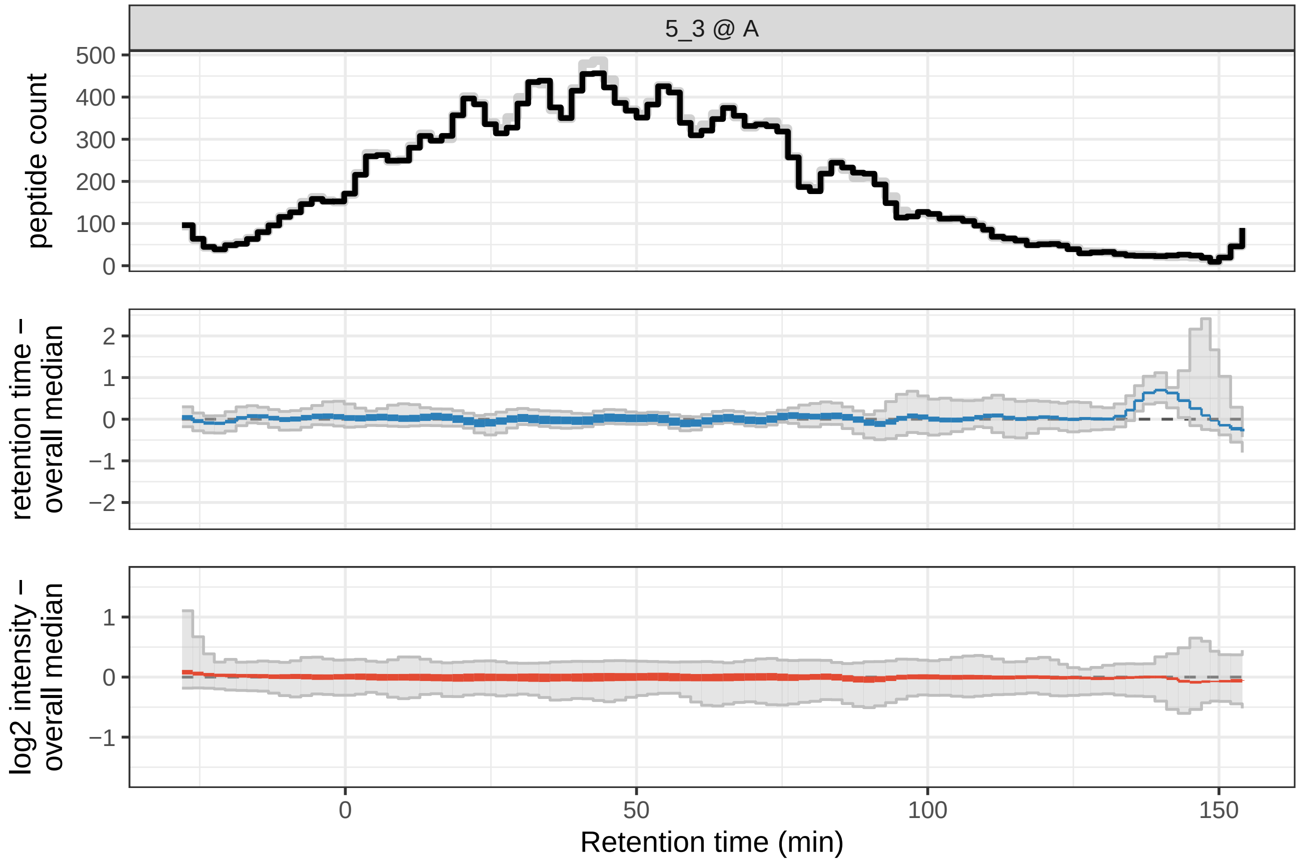
5_3 @ A
peptide count
500
400
300
200
100
0
retention time –
overall median
2
1
0
−1
−2
log2 intensity –
overall median
1
0
−1
0
50
100
150
Retention time (min)

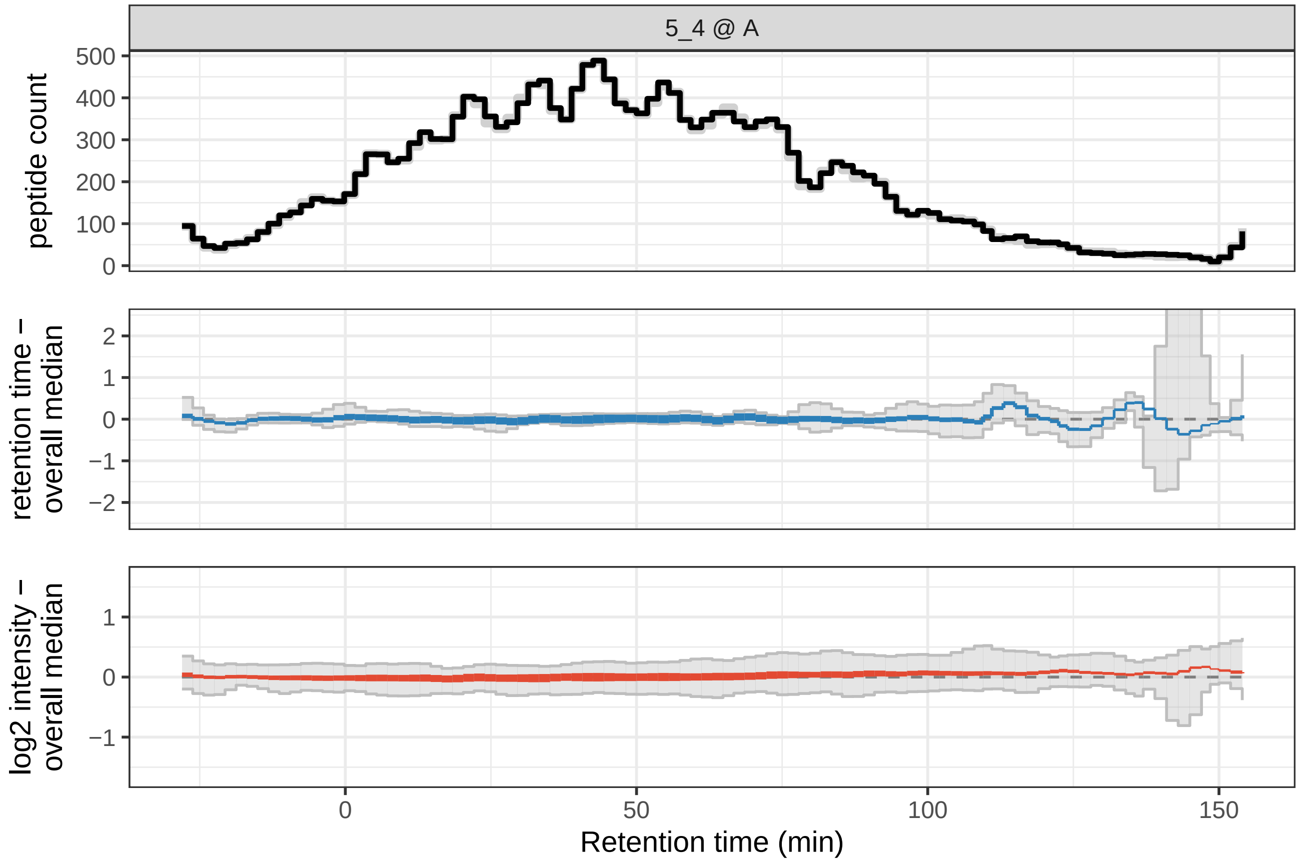
5_4 @ A
peptide count
500
400
300
200
100
0
retention time –
overall median
2
1
0
−1
−2
log2 intensity –
overall median
1
0
−1
0
50
100
150
Retention time (min)

#### 1.5 variation among replicates

The reproducibility of replicate measurements is expressed in three different analyses. First, we visualize for each sample how different its peptide intensities are compared to the mean value among replicates. Next, we use the Coefficient of Variation (CoV) as a metric for reproducibility and explore how much the CoV within a sample group can be improved by removing a single sample (eg; if CoV strongly improved after removing sample s, it was an outlier). Finally, we visualize the CoV within each sample group as a boxplot and a violin plot, figures commonly seen in proteomics literature and useful for comparing across experiments (of similar protocol).

##### 1.5.1 foldchange distributions

The foldchange of all peptides in a sample is compared to their respective mean value over all samples in the group. This visualizes how strongly each sample deviates from other samples in the same group which helps identify outlier samples. The same data was used as detailed in the "retention time" section above. Samples marked as 'exclude' in the provided sample metadata table are visualized as dashed lines.

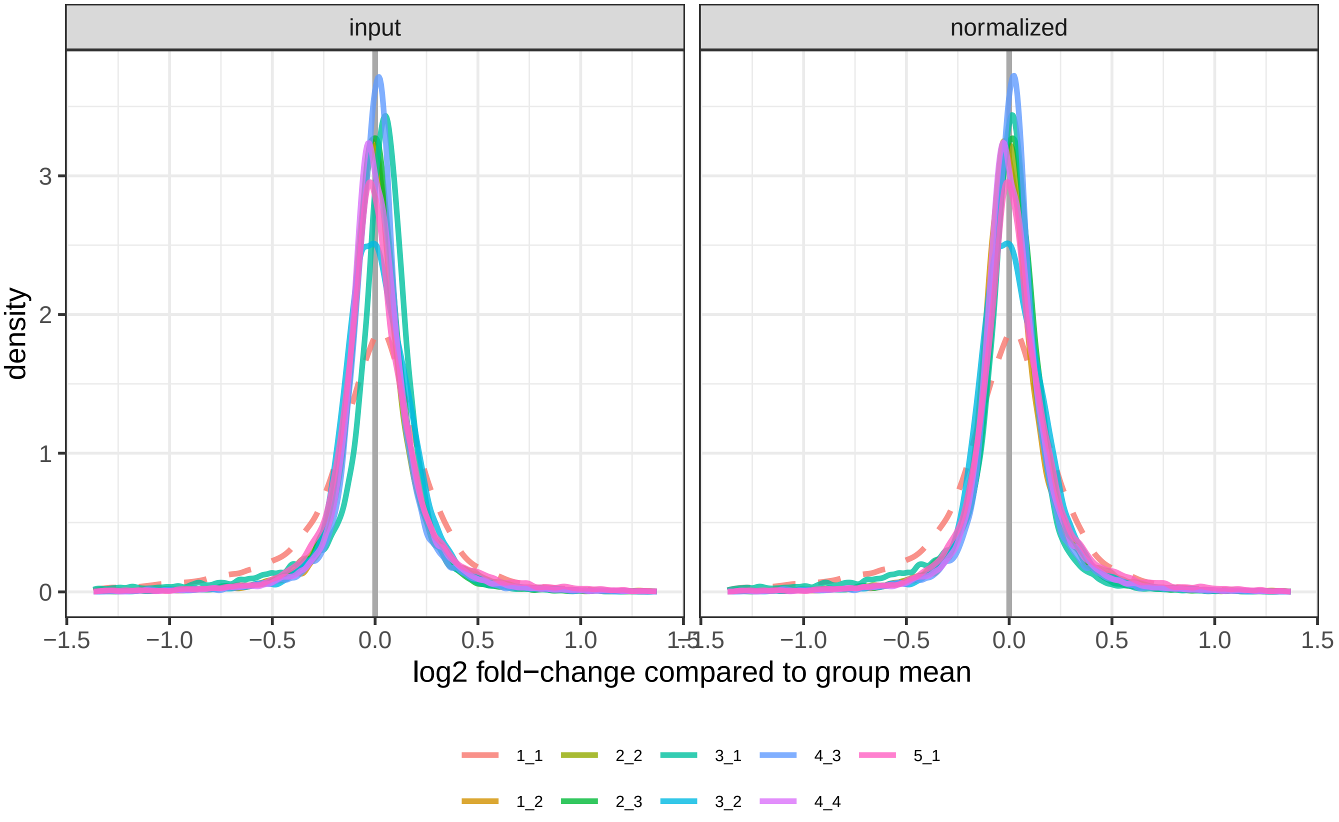

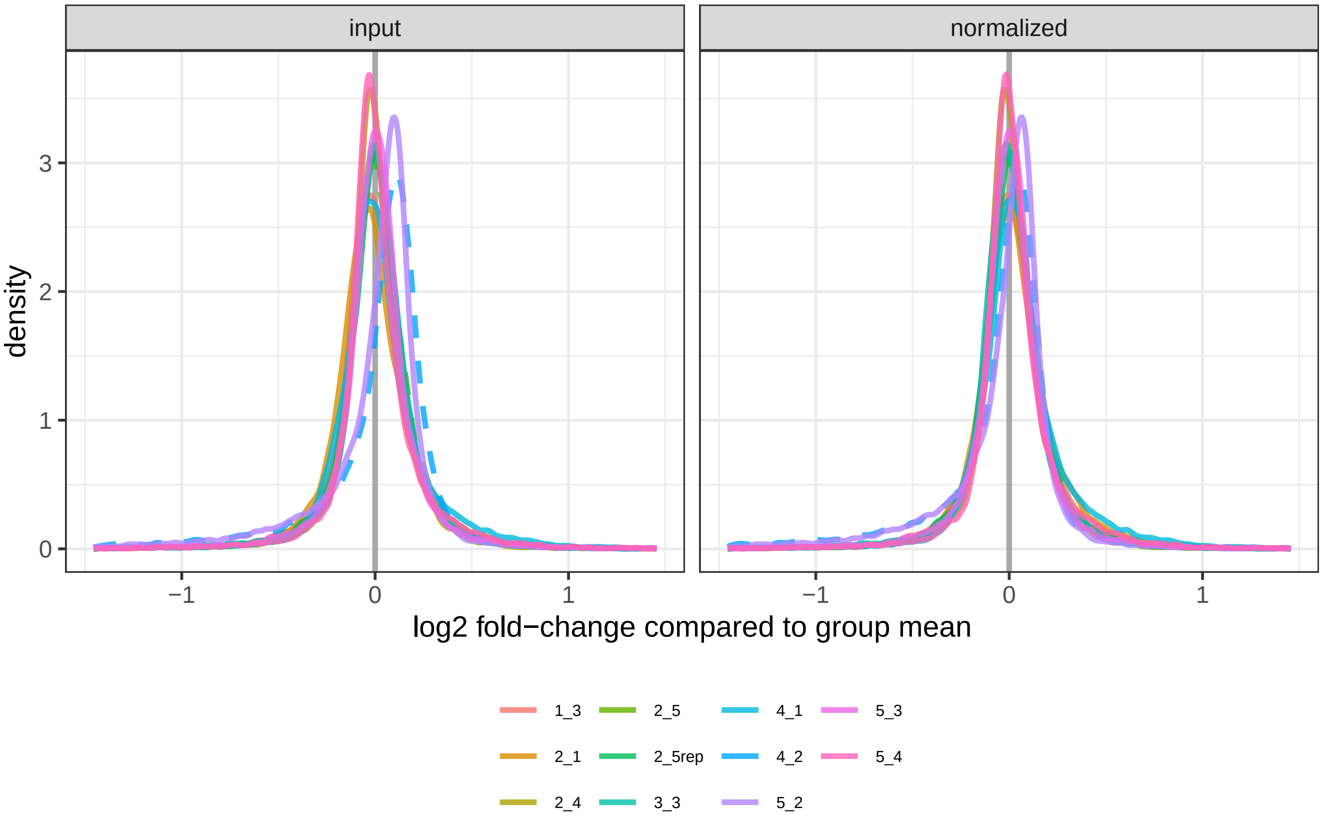
input
normalized
density
3
2
1
0
−1
0
1
log2 fold−change compared to group mean
1_3
2_5
4_1
5_3
2_1
2_5rep
4_2
5_4
2_4
3_3
5_2

##### 1.5.2 CoV, leave-one-out

progress: leave-one-out CoV plot computations took 20 seconds This set of figures describes the effect of removing a particular sample prior to within-group Coefficient of Variation (CoV) computation. The lower the CoV distribution is for a sample, the better reproducibility we get by excluding it. Only sample groups with at least 4 replicates can be used for this analysis, so 3 samples remain after leaving one out. Samples marked as 'exclude' by the user are included in these analyses (shown as dotted lines), and only peptides with at least 3 data points across replicates samples (after leave-one-out) are used for each CoV computation.

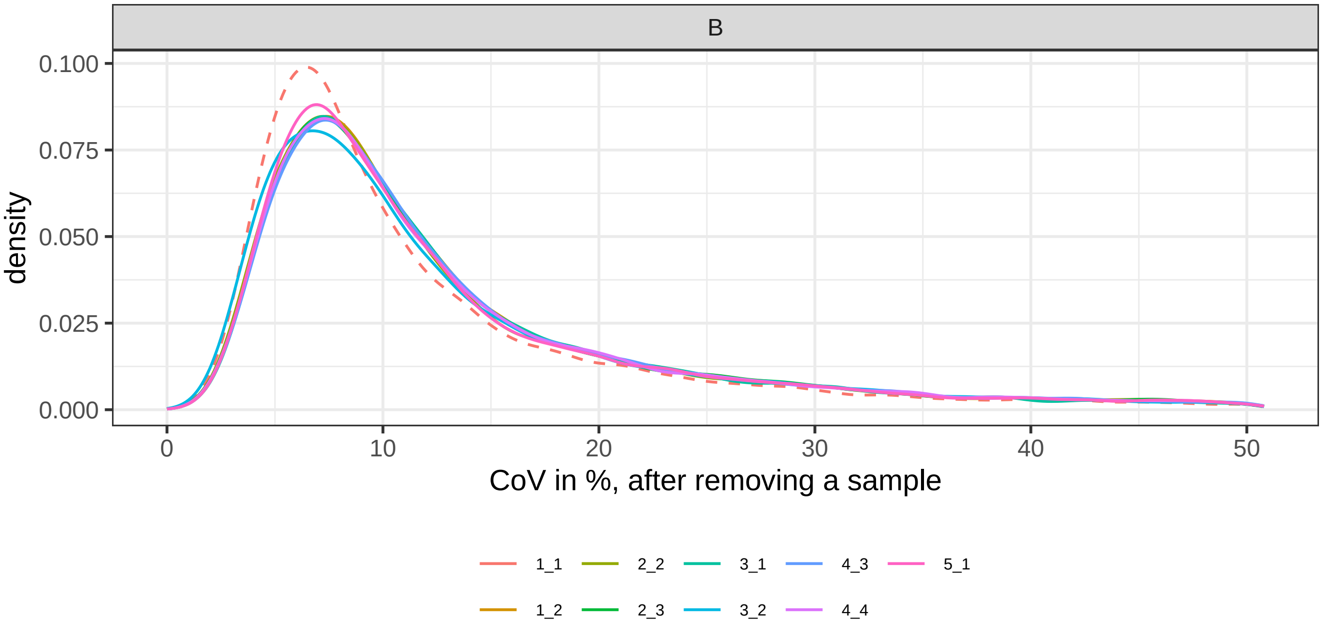

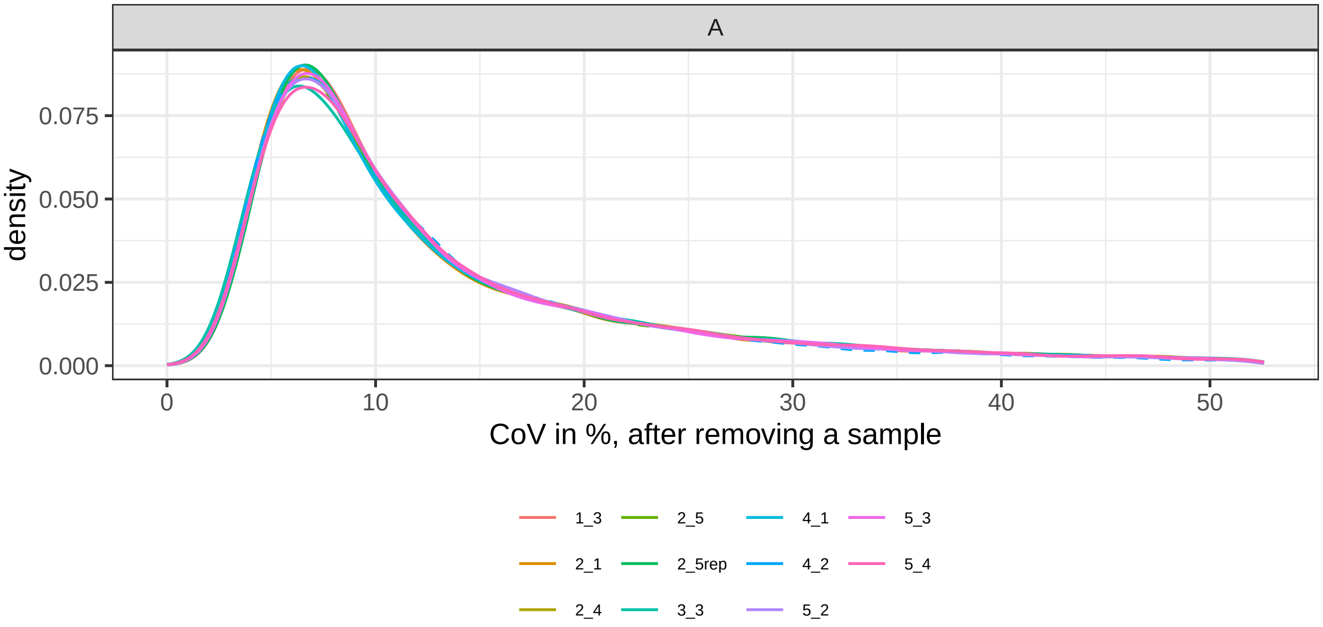

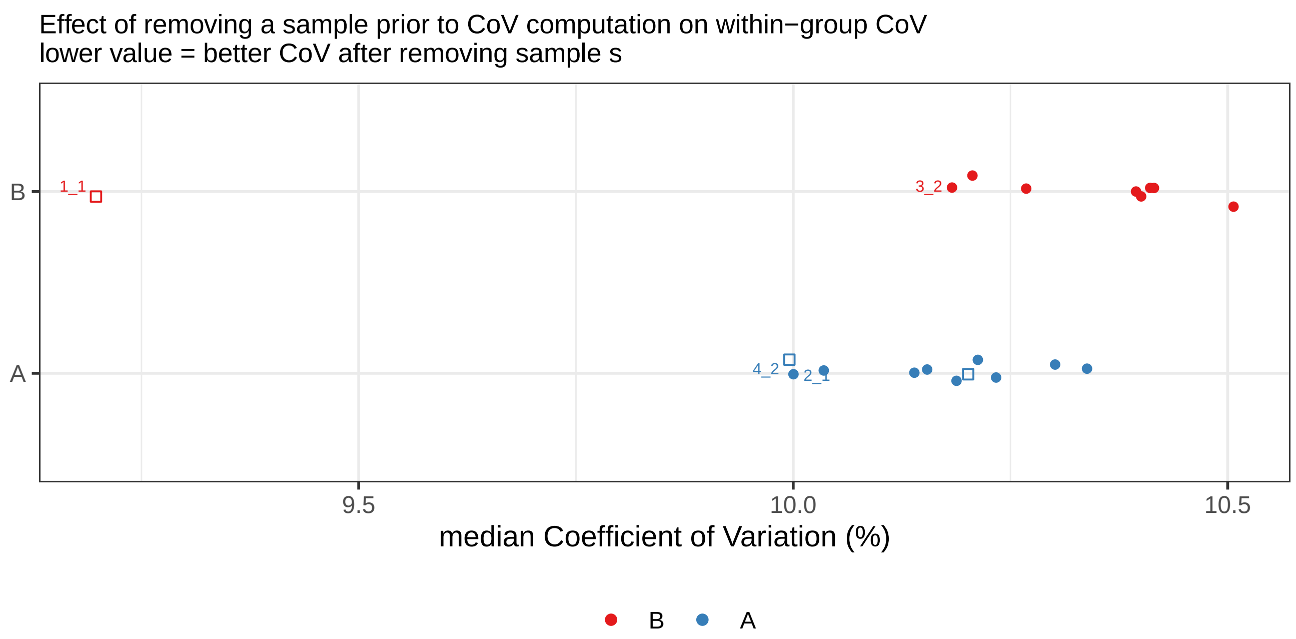

Effect of removing a sample prior to CoV computation on within−group CoV
lower value = better CoV after removing sample s
B
A
1_1
3_2
4_2
2_1
9.5
10.0
10.5
median Coefficient of Variation (%)
B
A

| shortname | group | exclude | median CoV |
|---|---|---|---|
| 1_1 | B | TRUE | 9.2 |
| 3_2 | B | FALSE | 10.2 |
| 5_1 | B | FALSE | 10.2 |
| 2_2 | B | FALSE | 10.3 |
| 3_1 | B | FALSE | 10.4 |
| 1_2 | B | FALSE | 10.4 |
| 4_4 | B | FALSE | 10.4 |
| 2_3 | B | FALSE | 10.4 |
| 4_3 | B | FALSE | 10.5 |
| 4_2 | A | TRUE | 10.0 |
| 2_1 | A | FALSE | 10.0 |
| 4_1 | A | FALSE | 10.0 |
| 1_3 | A | FALSE | 10.1 |
| 5_2 | A | FALSE | 10.2 |
| 2_5rep | A | FALSE | 10.2 |
| 2_5 | A | TRUE | 10.2 |
| 3_3 | A | FALSE | 10.2 |
| 5_3 | A | FALSE | 10.2 |
| 2_4 | A | FALSE | 10.3 |
| 5_4 | A | FALSE | 10.3 |

*Leave-one-out impact on within-group CoV (%)*

##### 1.5.3 Coefficient of Variation

Visualize the CoV, a quality metric for the reproducibility of replicate measurements, using box- and violin-plots.

The same data was used as detailed in the "retention time" section above, with only peptides quantified in at least 3 replicates used for CoV computation in a group. Only samples that are NOT marked 'exclude' and are in a sample group among at least 3 replicates are used for these figures.

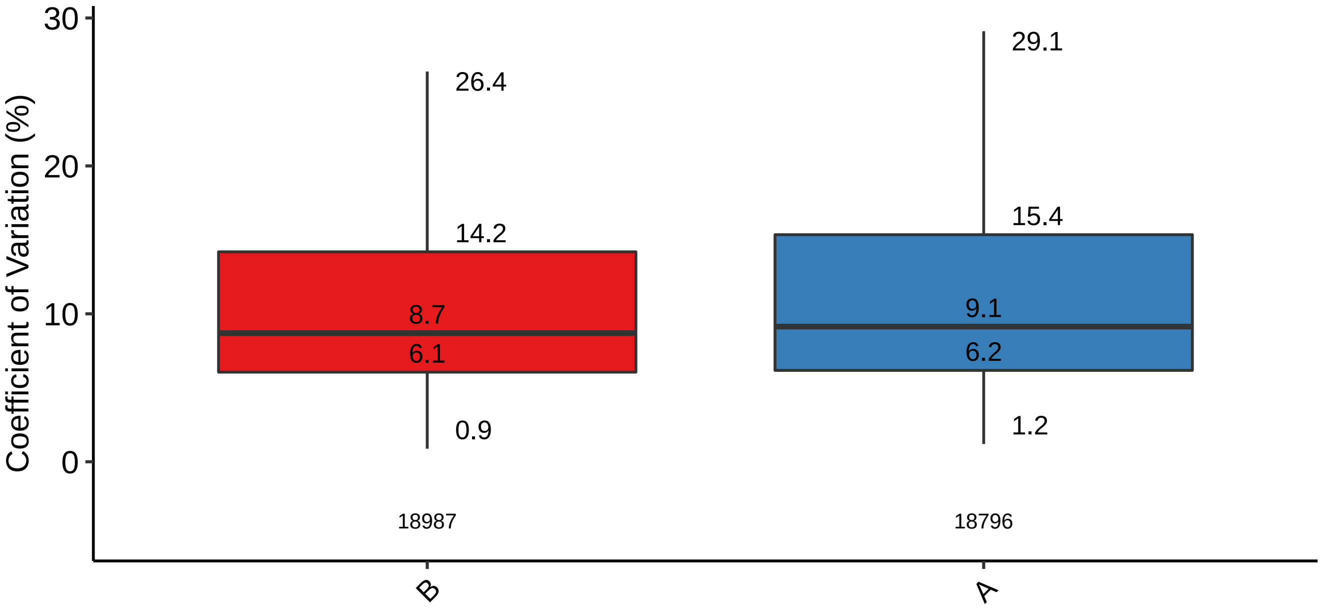

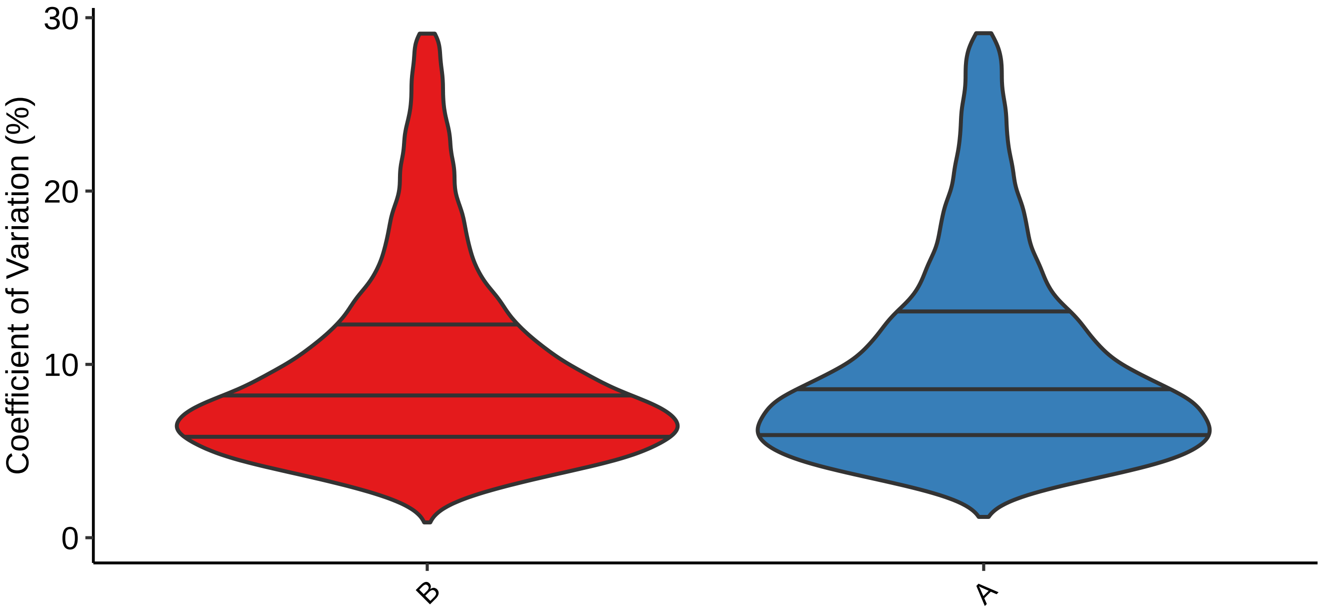

#### 1.6 PCA

A visualization of the first three PCA dimensions illustrates sample clustering. The goal of these figures is to pick up global effects from a quality control perspective, such as samples from the same experiment batch clustering together, not to be sensitive to a minor subset of differentially abundant proteins (for which specialized statistical models are applied downstream).

If additional sample metadata was provided, such as experiment batch, sample-prep dates, gel, etcetera, multiple PCA figures will be generated with respective color-codings. Users are encouraged to provide relevant experiment information as sample metadata and use these figures to inspect for unexpected batch effects.

The pcaMethods R package is used to perform the Probabilistic PCA (PPCA). The set of peptides used for this analysis consists of those peptides that pass your filter criteria in every sample group. If any samples are marked as 'exclude' in the sample metadata table, an additional PCA plot is generated with these samples included (depicting the 'exclude' samples as square symbols).

**Rationale behind data filter**
As mentioned above, the aim of these figures is to identify global effects. To achieve this, we compute sample clusters using the subset of peptides identified in each group which prevents rarely detected peptides/proteins from having a disproportionate effect on sample clustering. This pertains not only to 'randomly detected contaminant proteins' but also to proteins with abundance levels near the detection limit, which may be detected in only a subset of samples (eg; some measurements will be more successful/sensitive than others).

**Figure legends**
Left- and right-side figure panels on each row represent the same figure without and with sample labels. Samples that are flagged as 'exclude', if any, are visualized as square shapes.

**PCA of all samples, including those flagged as ‘exclude’, using 17955 peptides**

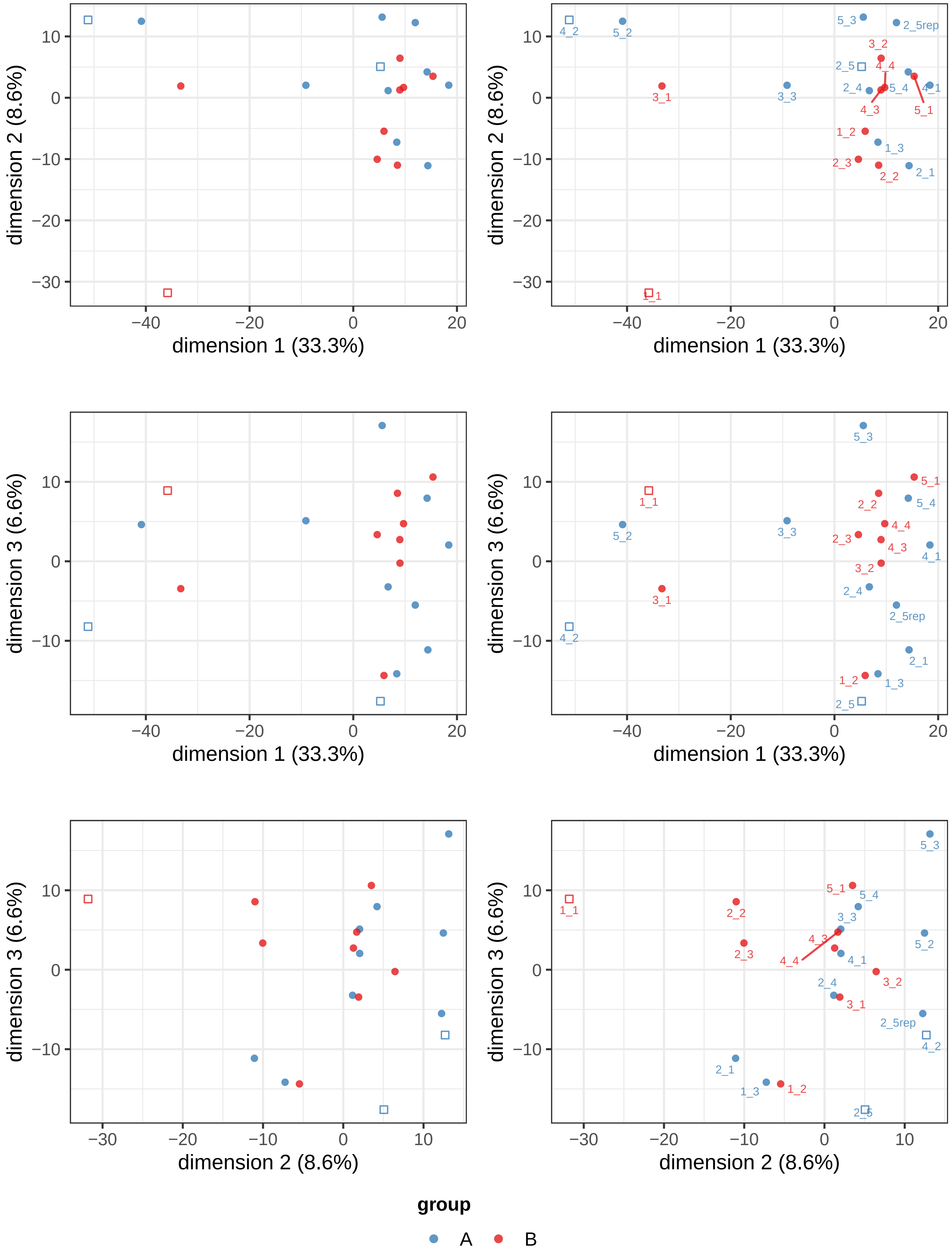

**PCA only on samples not flagged as 'exclude', using 18309 peptides**

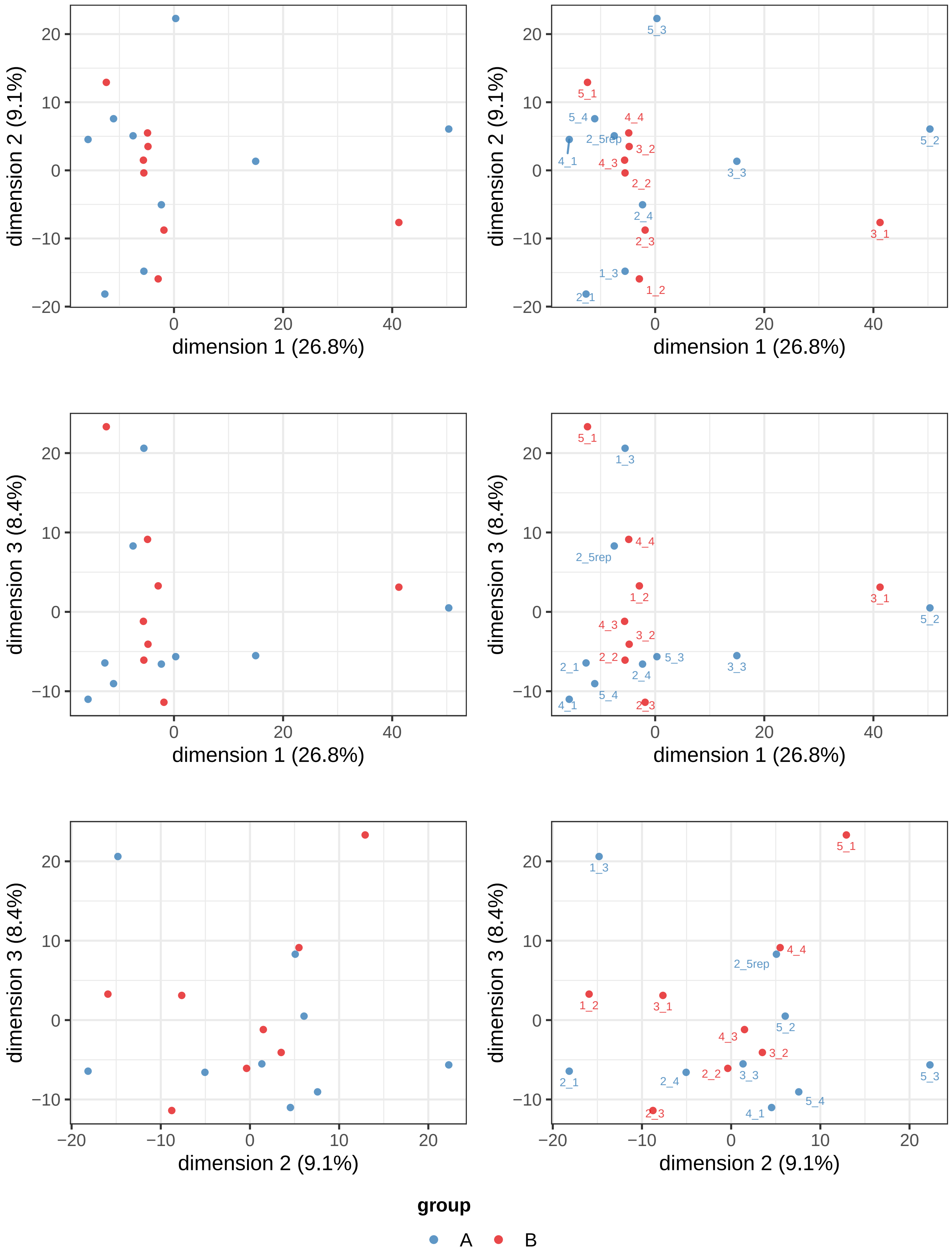

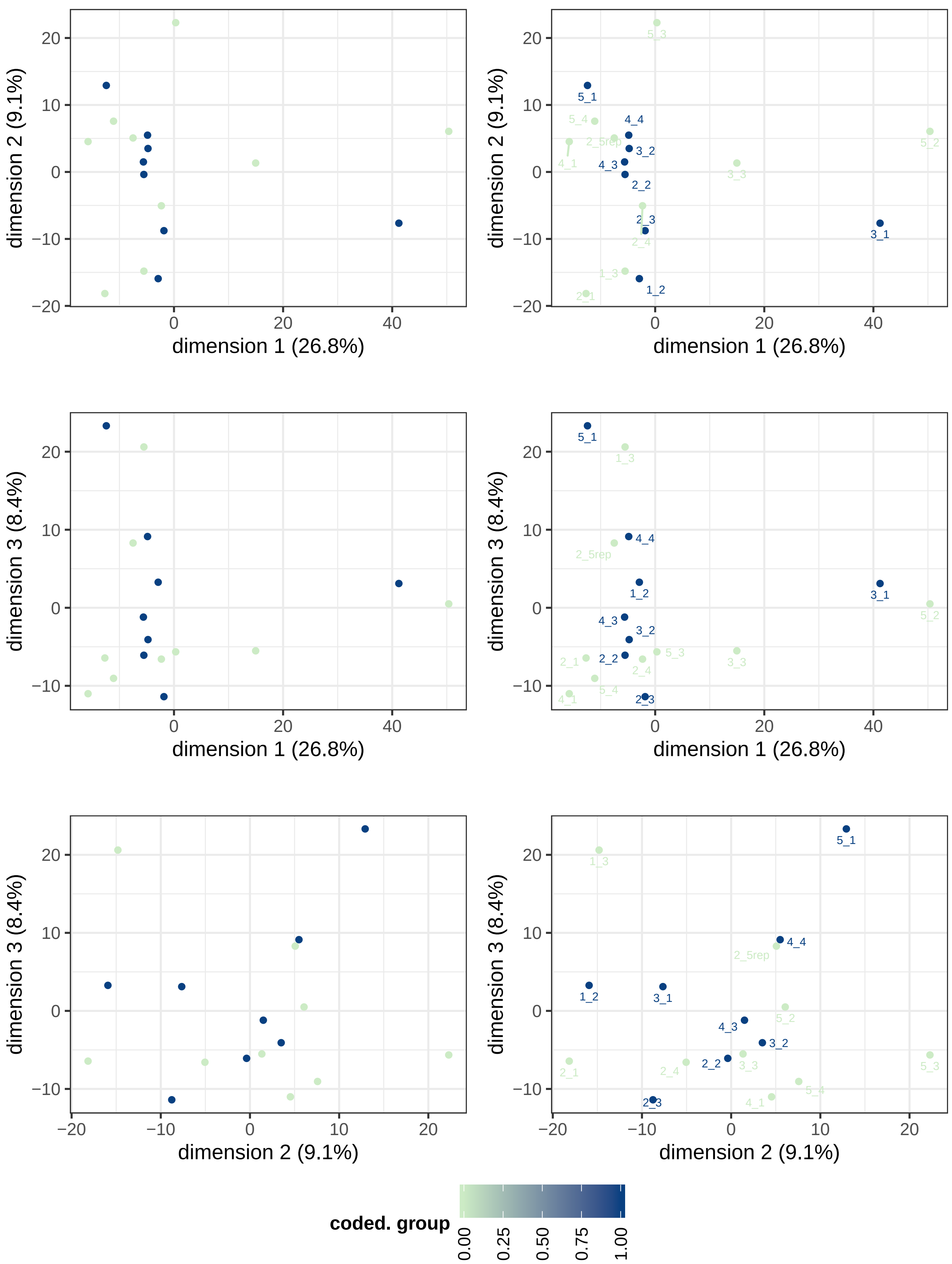
dimension 1 (26.8%)
dimension 2 (9.1%)
dimension 3 (8.4%)
5_3
5_1
5_4
4_4
2_5rep
5_2
4_1
3_2
4_3
3_3
2_2
2_3
2_4
3_1
1_3
1_2
2_1
5_4
coded. group
0.00
0.25
0.50
0.75
1.00

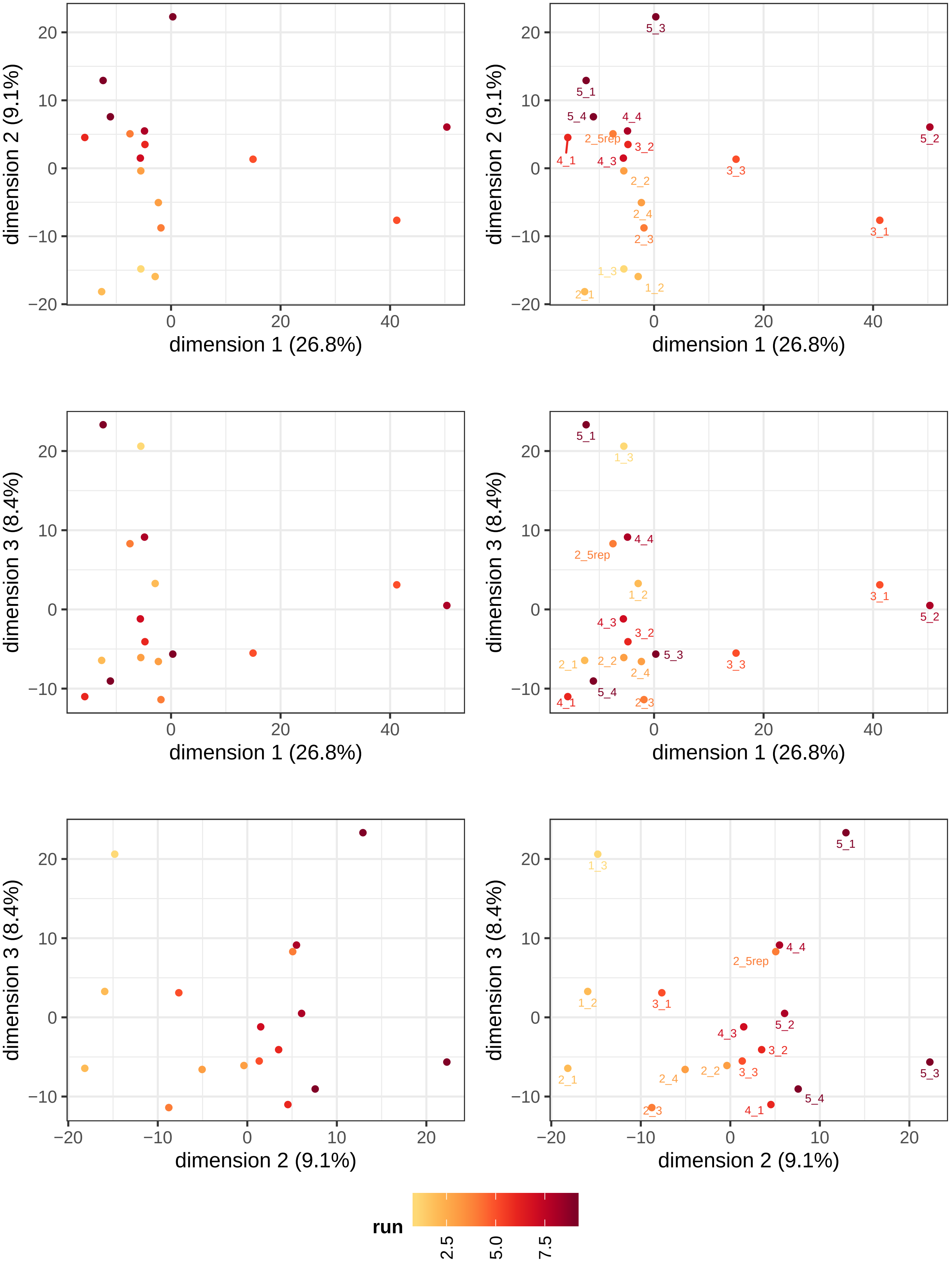
dimension 1 (26.8%)
dimension 2 (9.1%)
dimension 3 (8.4%)
5_3
5_1
5_4
4_4
2_5rep
3_2
4_1
4_3
5_2
3_3
2_2
2_4
2_3
3_1
1_3
1_2
2_1
run
2.5
5.0
7.5

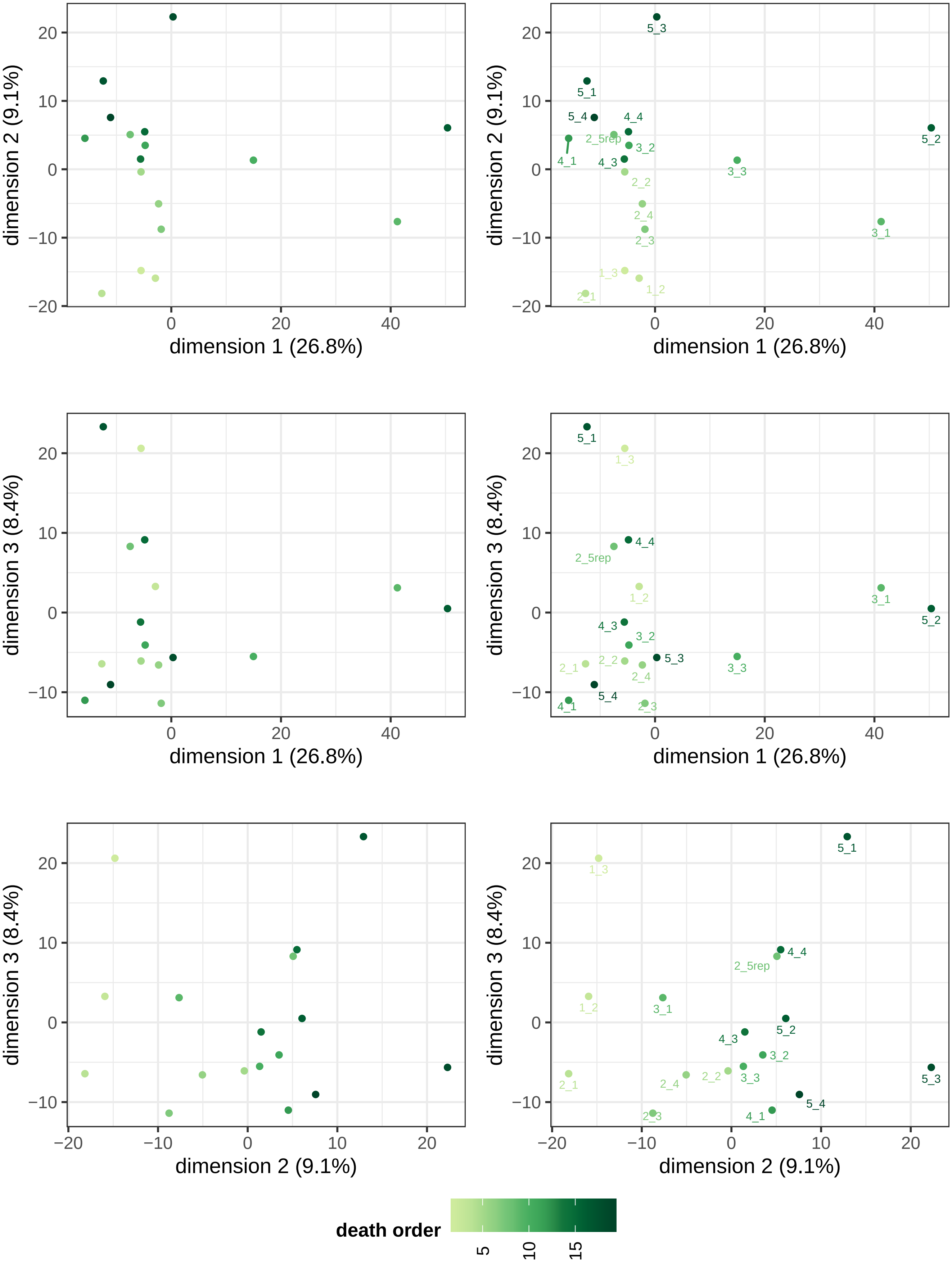

dimension 1 (26.8%)
dimension 2 (9.1%)
dimension 3 (8.4%)
5_1
5_2
5_3
5_4
4_1
4_3
4_4
3_1
3_2
3_3
2_1
2_2
2_3
2_4
2_5rep
1_2
1_3
death order
5
10
15

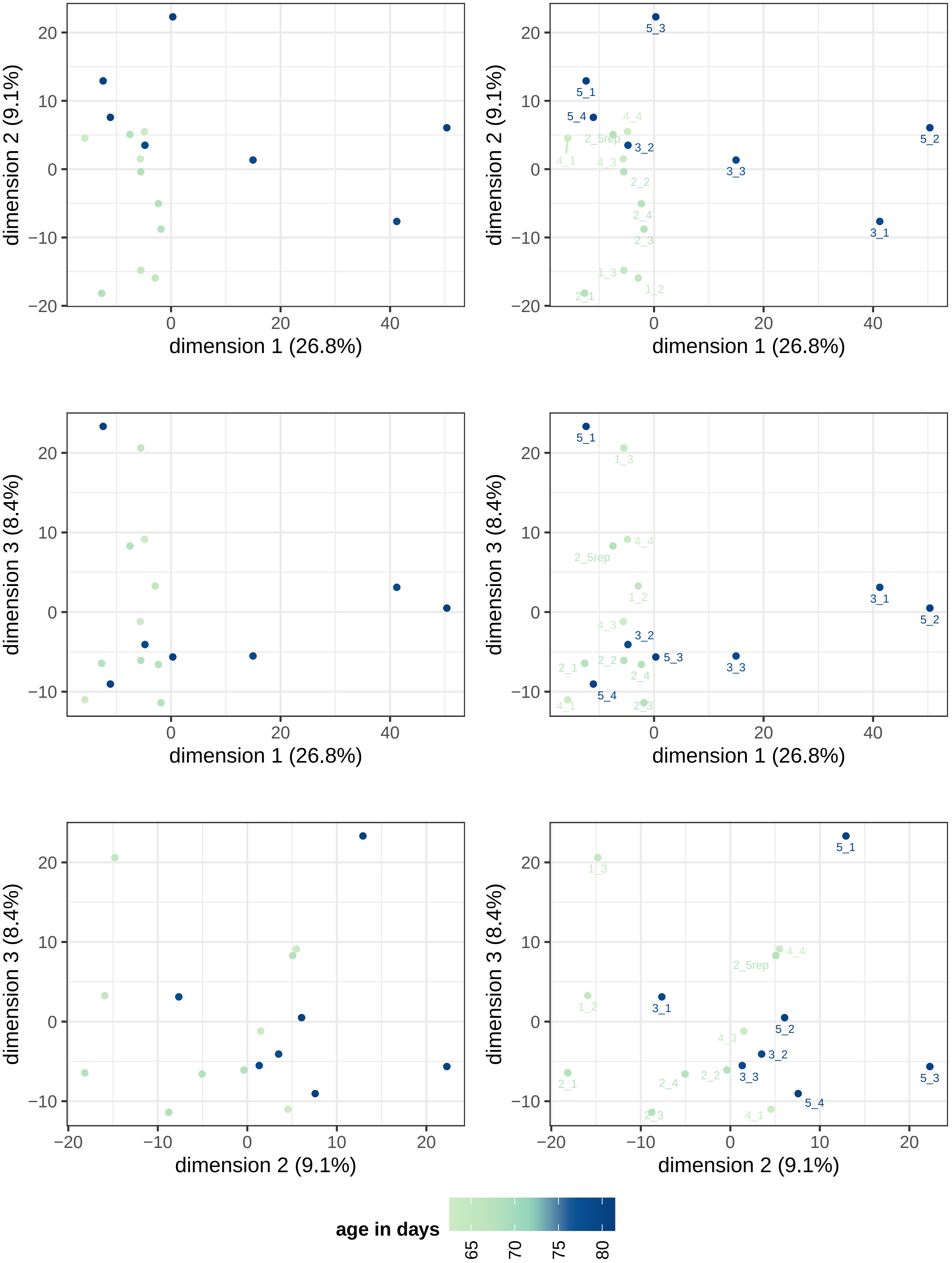
dimension 1 (26.8%)
dimension 2 (9.1%)
dimension 3 (8.4%)
5_1
5_2
5_3
5_4
3_1
3_2
3_3
4_1
4_3
4_4
2_1
2_2
2_3
2_4
2_5rep
1_2
1_3
age in days
65
70
75
80

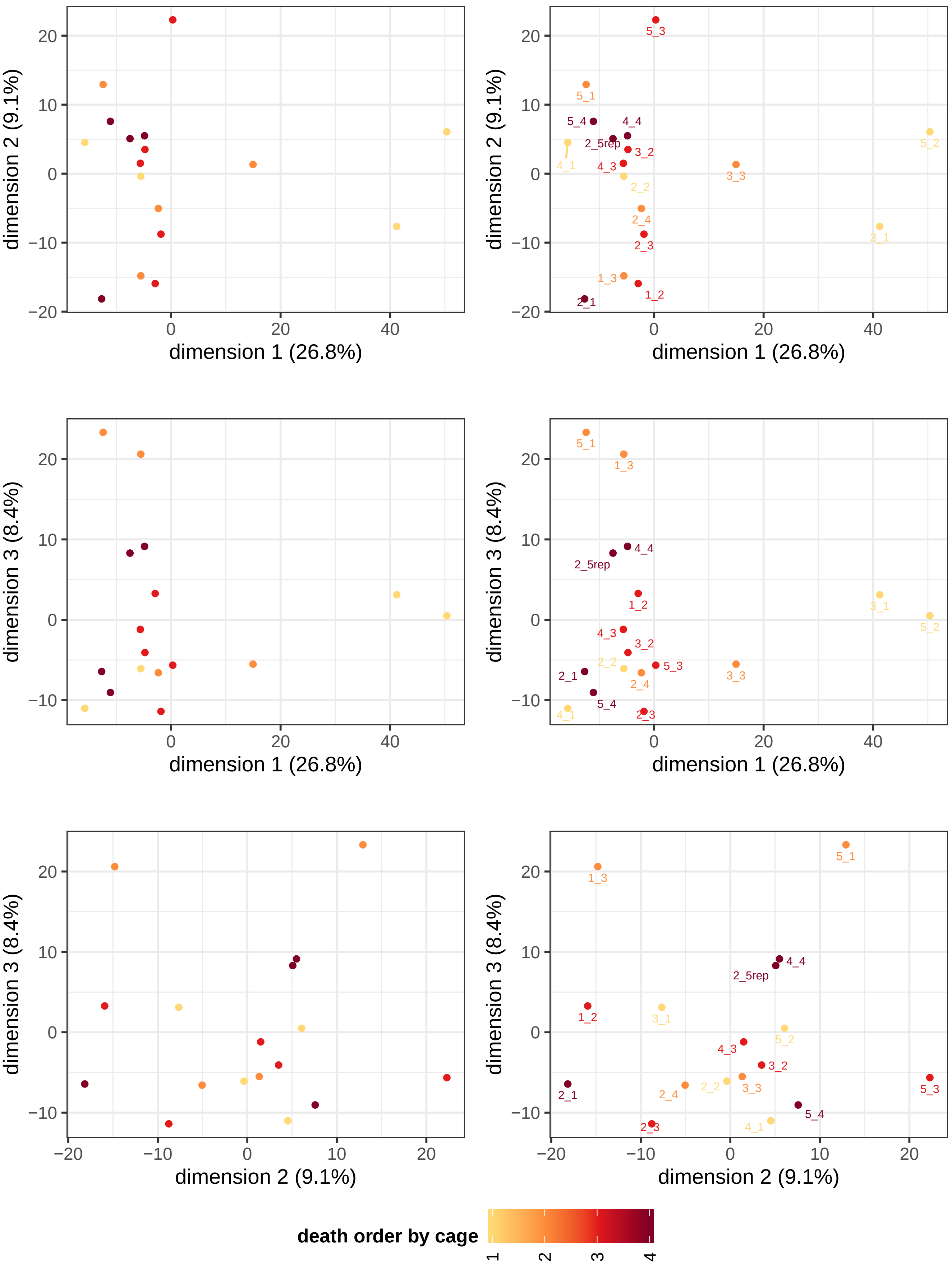
dimension 1 (26.8%)
dimension 2 (9.1%)
dimension 3 (8.4%)
5_3
5_1
5_4
4_4
2_5rep
3_2
4_1
4_3
2_2
3_3
5_2
2_4
2_3
3_1
1_3
1_2
2_1
death order by cage
1
2
3
4

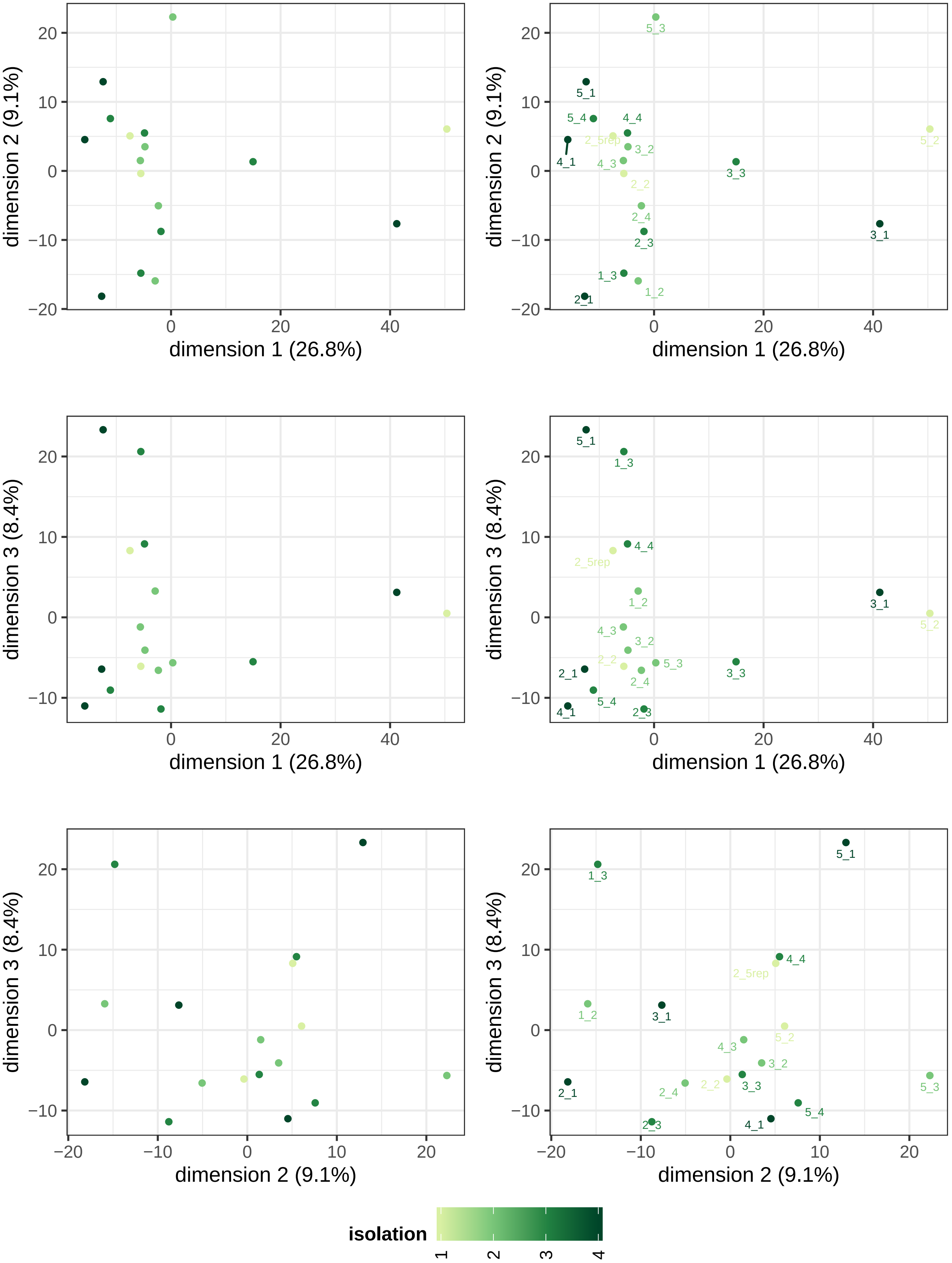
dimension 1 (26.8%)
dimension 2 (9.1%)
dimension 3 (8.4%)
5_3
5_1
5_4
4_4
2_5rep
4_1
3_2
4_3
2_2
3_3
5_2
2_4
2_3
3_1
1_3
1_2
2_1
isolation
1
2
3
4

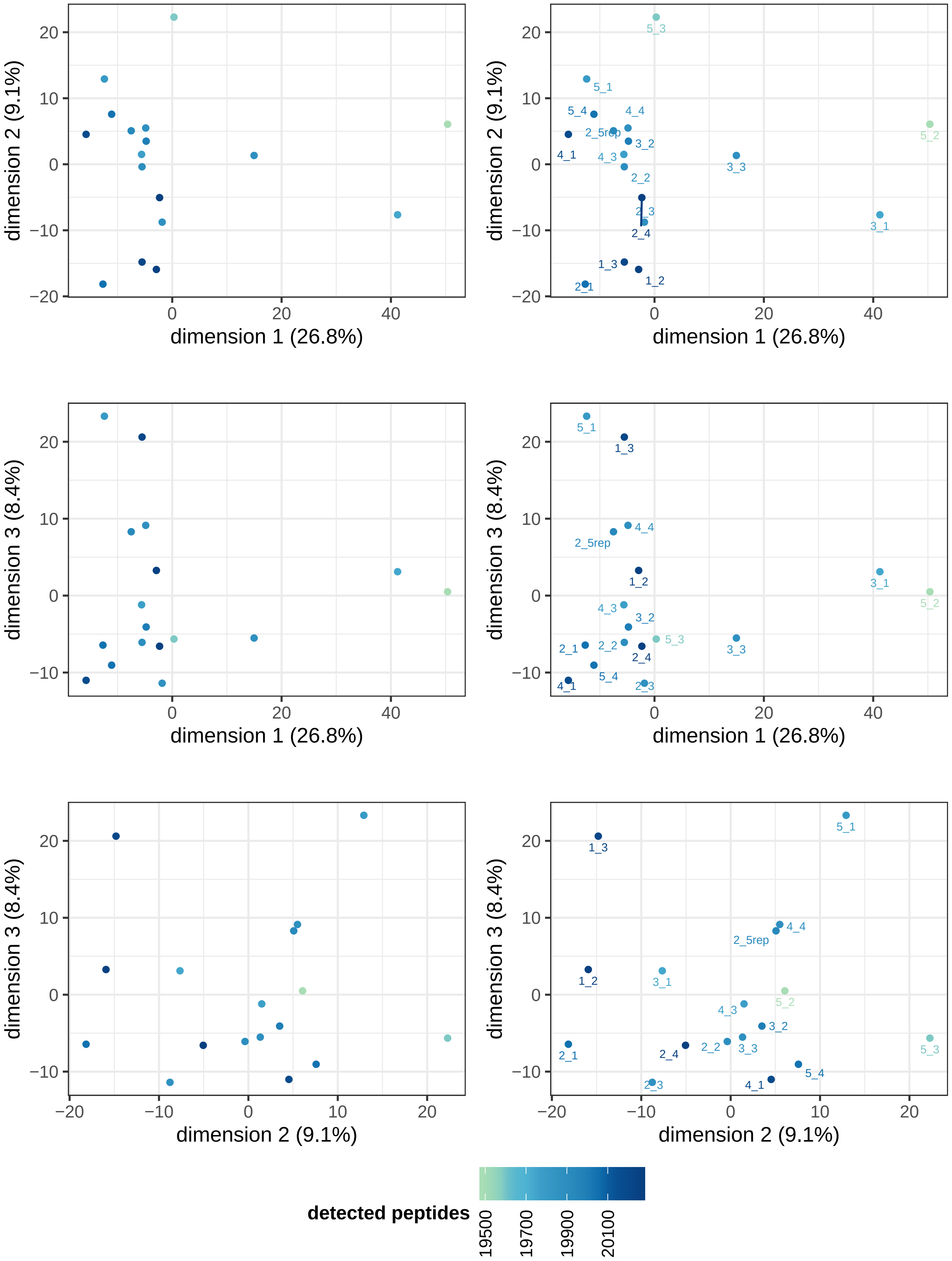
dimension 1 (26.8%)
dimension 2 (9.1%)
dimension 3 (8.4%)
5_3
5_1
5_4
4_4
2_5rep
3_2
4_1
4_3
5_2
3_3
2_2
2_3
2_4
3_1
1_3
1_2
2_1
detected peptides
19500
19700
19900
20100

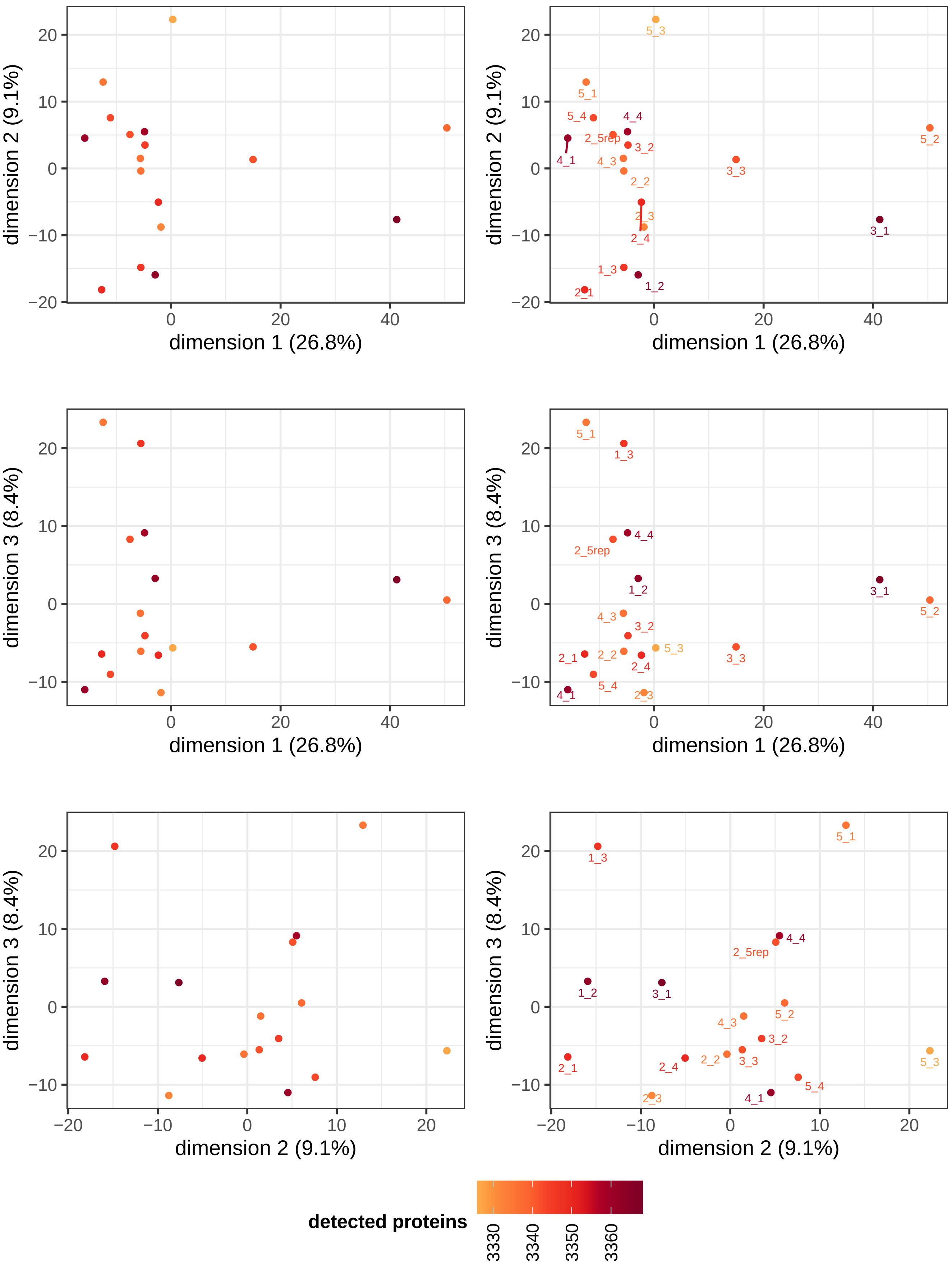

dimension 1 (26.8%)
dimension 2 (9.1%)
dimension 3 (8.4%)
5_3
5_1
5_4
4_4
2_5rep
3_2
4_1
4_3
2_2
5_2
3_3
2_3
2_4
3_1
1_3
2_1
1_2
detected proteins
3330
3340
3350
3360

### 2 Differential abundance analysis

**goal: maximize reliable features for quantification**

In a pairwise analysis of two groups of samples, only peptides with N data-points in both groups are used for quantitative analysis (where N = defined by user settings). For example; if peptide *p* is consistently quantified in sample groups A and B but not in C/D/E, it can be used when comparing group A *versus* group B but should not be used in any other group comparisons. This approach is particularly suited to maximize the number of peptides used for statistical analysis in experimental designs with many sample groups.

A common alternative strategy is a global filtering approach where peptides are selected based on their properties in the overall dataset (eg; present in x% of samples or x% of replicates in all groups) and subsequentially the resulting data matrix is used for all downstream statistical analyses. In the example above where peptide *p* is present in a subset of sample groups, *p* would either be left out (not present in majority of samples in entire dataset) or erronously used when applying t-statistics to groups B and C (since *p* is not present in group C, it may differentially detected but there are no features available for quantitative analysis)

## 2.1 A vs B

- **user setting:** using 'global data filter' peptide filtering approach
- 23479 peptides in 3477 proteins remain in the current contrast after peptide filters and are used for the statistical analysis in this section
- qvalue threshold: 0.01
- log2 foldchange threshold: 0.156096

##### 2.1.1 volcano

Left- and right-side figure panels on each row represent the same figure without and with labels for the 25 proteins with lowest p-value.

Bottom figure panels have limited x- and y-axis. For datasets with a small number of strong outliers in p-value or fold-change, which may have a profound effect on the plot scales, this allows you to inspect the remainder of the volcano plot without disproportionate influence by 'extreme' values.

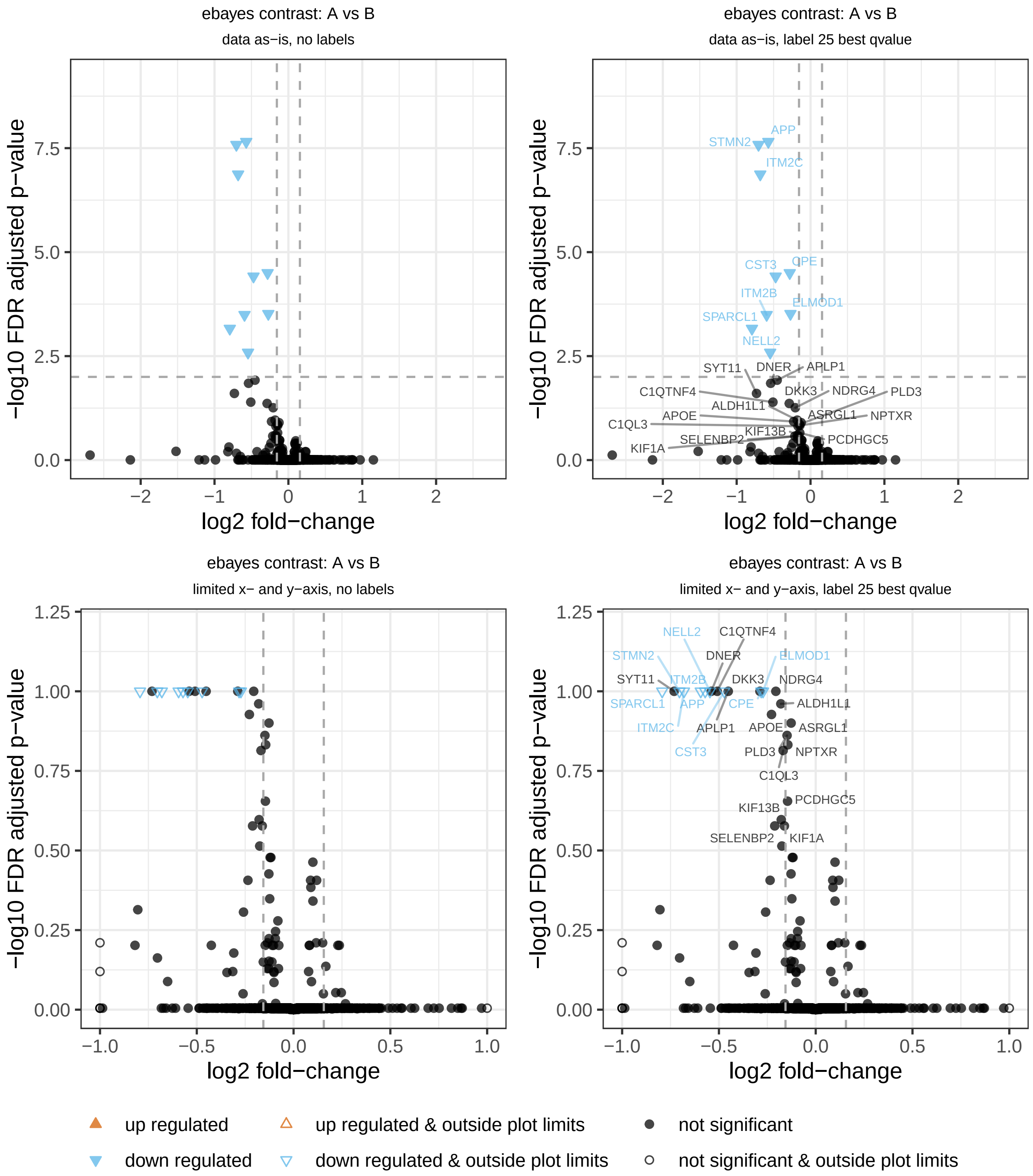
ebayes contrast: A vs B
data as−is, no labels
data as−is, label 25 best qvalue
limited x− and y−axis, no labels
limited x− and y−axis, label 25 best qvalue
−log10 FDR adjusted p−value
log2 fold−change
APP
STMN2
ITM2C
CST3
CPE
ITM2B
ELMOD1
SPARCL1
NELL2
SYT11
DNER
APLP1
C1QTNF4
DKK3
NDRG4
PLD3
ALDH1L1
APOE
ASRGL1
NPTXR
C1QL3
KIF13B
SELENBP2
PCDHGC5
KIF1A
up regulated
up regulated & outside plot limits
not significant
down regulated
down regulated & outside plot limits
not significant & outside plot limits

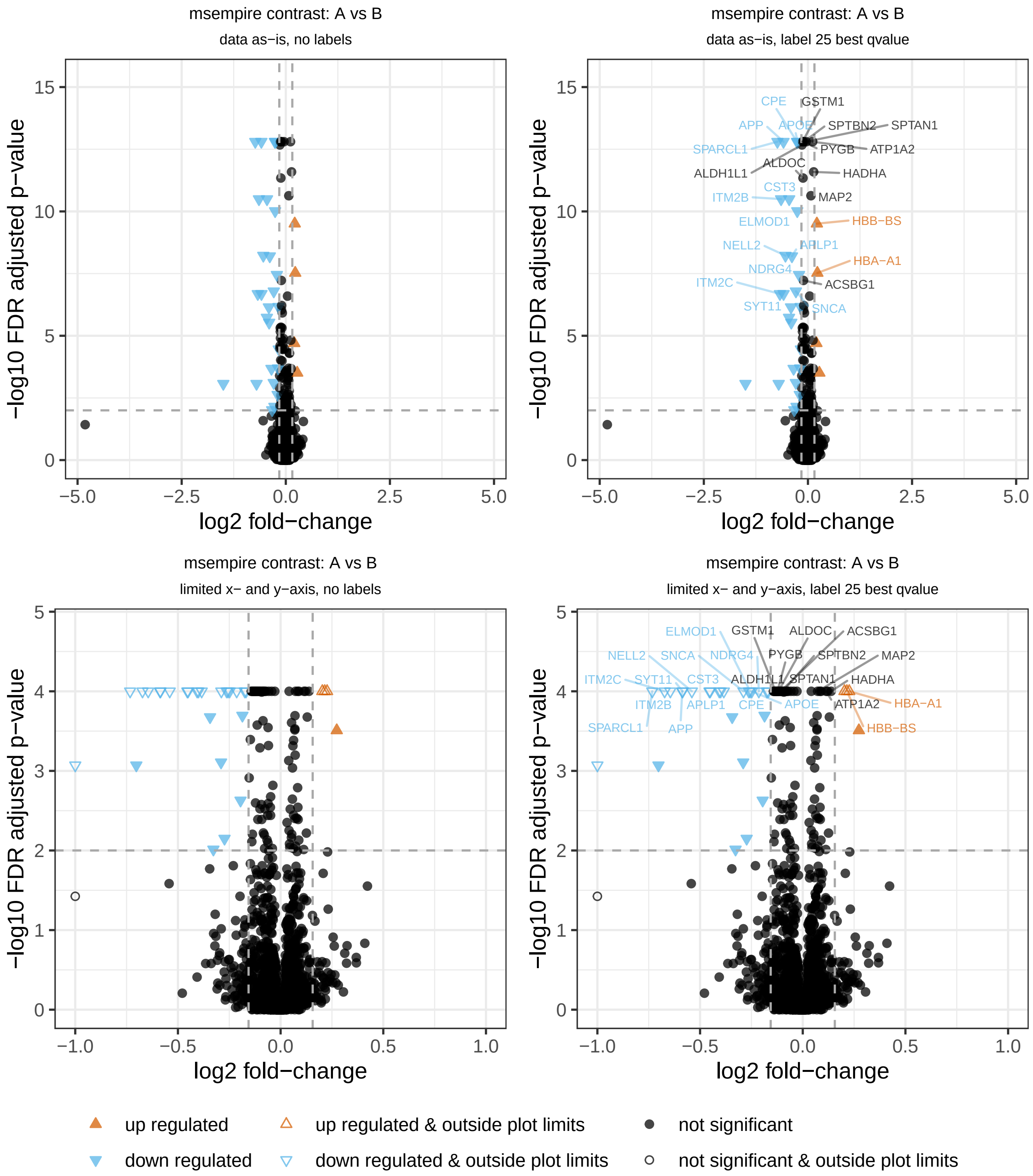

msempire contrast: A vs B
data as−is, no labels
msempire contrast: A vs B
data as−is, label 25 best qvalue
msempire contrast: A vs B
limited x− and y−axis, no labels
msempire contrast: A vs B
limited x− and y−axis, label 25 best qvalue
−log10 FDR adjusted p−value
log2 fold−change
up regulated
down regulated
up regulated & outside plot limits
down regulated & outside plot limits
not significant
not significant & outside plot limits

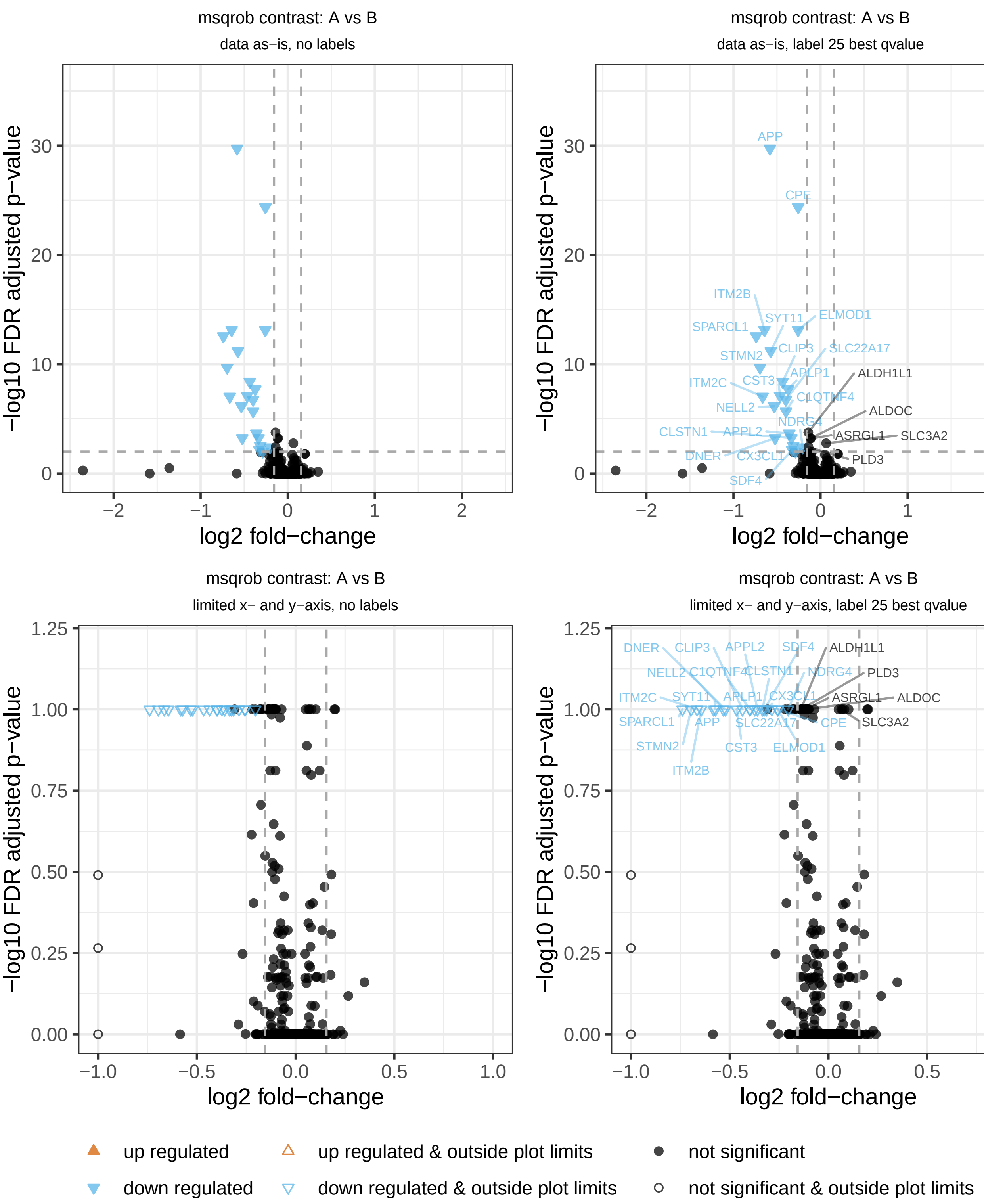

msqrob contrast: A vs B
data as−is, no labels
msqrob contrast: A vs B
data as−is, label 25 best qvalue
msqrob contrast: A vs B
limited x− and y−axis, no labels
msqrob contrast: A vs B
limited x− and y−axis, label 25 best qvalue
−log10 FDR adjusted p−value
log2 fold−change
up regulated
down regulated
up regulated & outside plot limits
down regulated & outside plot limits
not significant
not significant & outside plot limits

##### 2.1.2 foldchange distribution

Distributions of estimated foldchanges produced by the statistical models. If the mode is far from 0, consider alternative normalization strategies. Do note the scale on the x-axis, for some experiments the foldchanges are very low which in turn may exaggerate this figure.

*note; the MSqRob model tends to assign zero (log)foldchange for proteins with minor difference between conditions where the model is very sure the null hypothesis cannot be rejected (shrinkage by the ridge regression model). As a result, many foldchanges will be zero and the density plot for MSqRob may look like a spike instead of the expected Gaussian shape observed in other models*

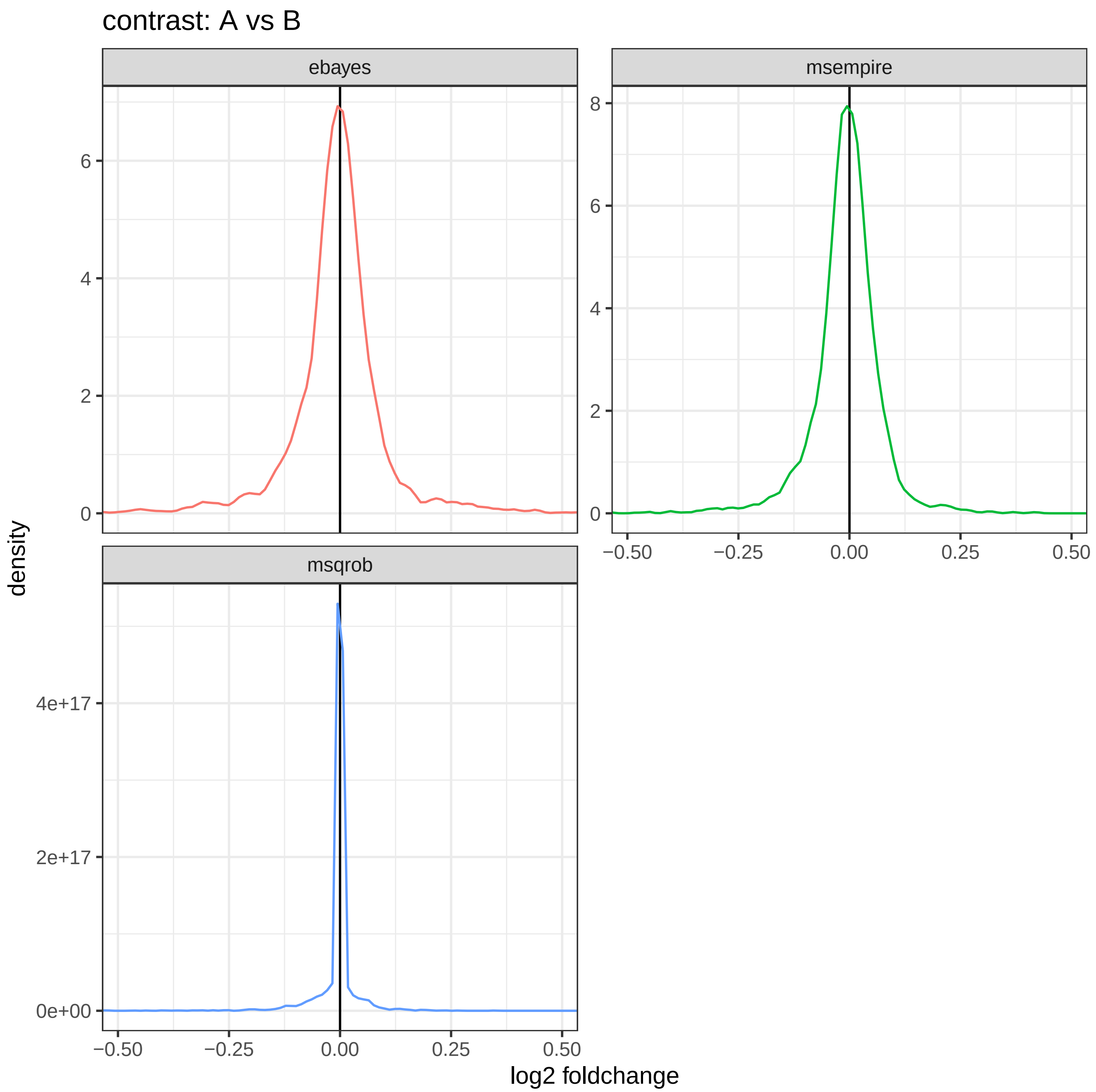

##### 2.1.3 p-value distribution

Histogram of p-values computed by differential expression analysis algorithms, as-is, for quality-control inspection. The horizontal line indicates the expected counts assuming a uniform distribution (total number of p-values divided by number of histogram bins)

See further: https://www.ncbi.nlm.nih.gov/pmc/articles/PMC6164648/

See further: http://varianceexplained.org/statistics/interpreting-pvalue-histogram/

*note; the MSqRob and MS-EmpiRe models often yield p-value distributions that show a large peak at p-value 1, these are typically proteins with estimated log foldchanges at/near zero where these models are very sure the null hypothesis cannot be rejected*

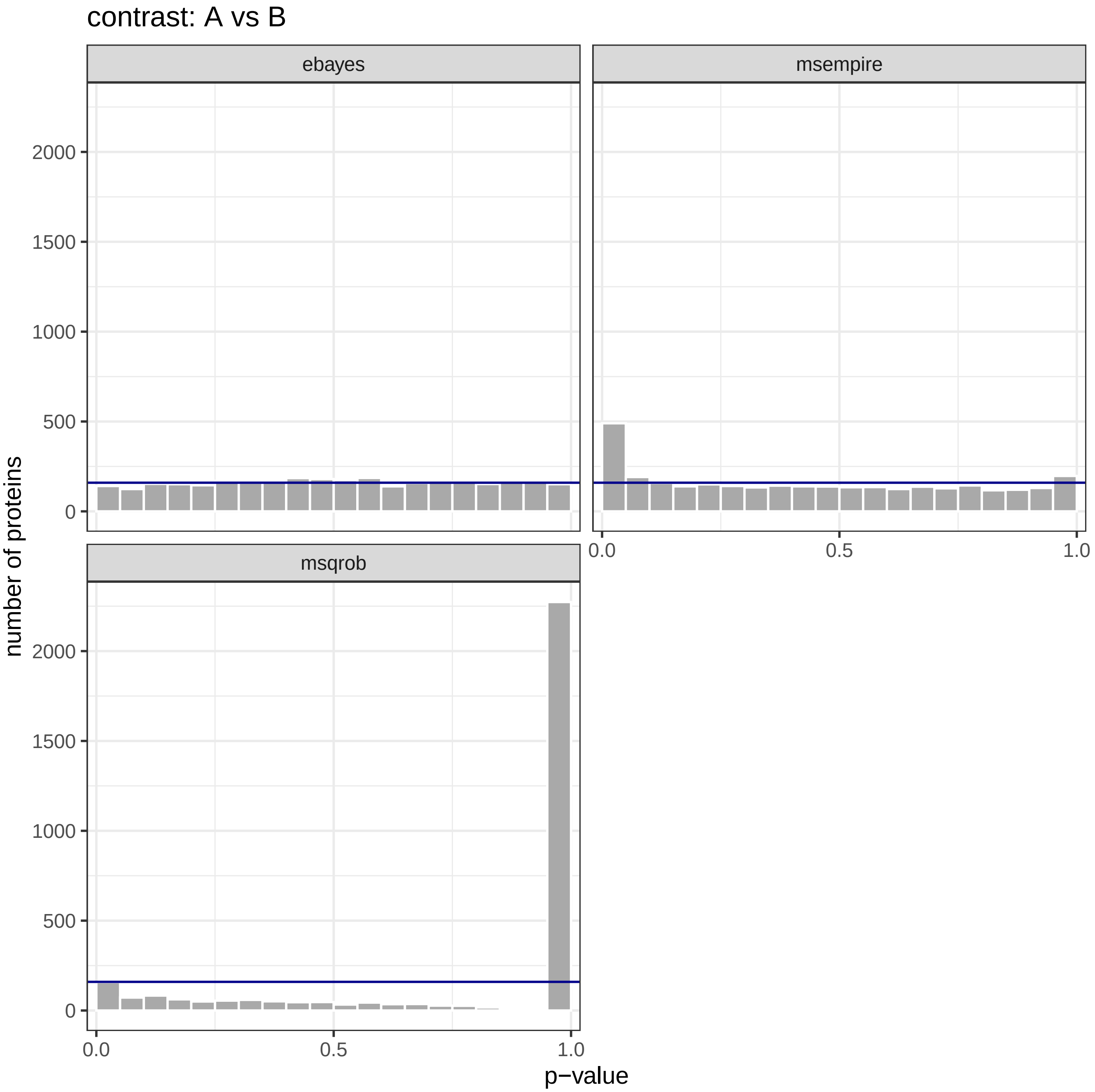

### 3 Summary of statistical results

Number of proteins found statistically significant in each comparison of sample groups.

| contrast | algorithm | #tested | #significant |
|---|---|---|---|
| A vs B | ebayes | 3188 | 9 |
| A vs B | msempire | 3188 | 30 |
| A vs B | msqrob | 3188 | 21 |

Table 1: Significant hits

### 4 log

reading Spectronaut report...

to plot target/decoy score distributions, include columns 'cscore' and 'isdecoy' (with at least some decoy entries) in the peptide report

23484/24992 precursors remain after selecting the 'best' precursor for each modified sequence

5 / 22486 unique target (plain)sequences were not in the provided spectral library and thus removed

8842/8842 protein accessions and 3477/3477 protein groups were mapped to provided fasta file(s)

protein filters didn't match anything

using 7 threads for multiprocessing

differential abundance analysis for contrast: A vs B

using data from peptide filter: global data filter

normalizing protein-level data for DEA by 'modebetween' (doesn't affect peptide-level models)

log2 foldchange threshold estimated by bootstrap analysis: 0.156

### 5 R command history

This shows the history commands from your R script that starts this pipeline, thereby automatically documenting the parameters/settings used. All lines of executed code since (last) importing data using this R package are shown.

##### Using this feature

Do not use RStudio's `source` option to execute our pipeline since it will only write `source(...yourscript.R)` to the session history, and consequentially that is all you see in this 'code log'. Instead, select all lines in your script (`control + A`) and then "run" the selected code (either click the run button in RStudio, or use `control + enter`). All lines shown in this section are the same as shown in the RStudio 'History' pane (a tab on the top-right of its UI).

```
norm_algorithm = "vwmb",
dea_algorithm = "ebayes",
# dea_algorithm = c("ebayes", "msempire", "msqrob"), # ebayes, msqrob, msempire, msqrobsum
dea_qvalue_threshold = 0.01,
dea_log2foldchange_threshold = 0,
output_qc_report = TRUE,
output_peptide_plots = "none",
output_abundance_tables = FALSE,
output_dir = "C:/DATA/Annemiek/data_analysis_results__subset",
output_within_timestamped_subdirectory = TRUE)
### Annemiek; subset
rm(list = ls(all.names = TRUE)) # clear everything from memory
cat("\014") # clear terminal (send the control+L character)
devtools::load_all() # load our R package
dataset = import_dataset_spectronaut("C:/DATA/Annemiek/spectronaut/20200517_182020_pilot1_ngn2_induced_neurons__subset_Report.xls", conf
dataset = update_protein_mapping_from_maxquant(dataset, path_maxquant = "C:/DATA/Annemiek/maxquant_txt", remove_shared = TRUE)
dataset = import_fasta(dataset, files = c("C:/DATA/Annemiek/UniProt_2019-11/UP000005640_9606.fasta", "C:/DATA/Annemiek/UniProt_2019-11/UI
dataset = remove_proteins_by_name(dataset, remove_irt_peptides = TRUE) # optional
## create metadata file by subsetting original table
# x = openxlsx::read.xlsx("C:/DATA/Annemiek/pilot1_ngn2_induced_neurons_sample-metadata.xlsx")
# openxlsx::write.xlsx(x[x$sample_id %in% dataset$peptides$sample_id,], "C:/DATA/Annemiek/pilot1_ngn2_induced_neurons_sample-metadata__sub
dataset = import_sample_metadata(dataset, filename = "C:/DATA/Annemiek/pilot1_ngn2_induced_neurons_sample-metadata__subset.xlsx")
# -       42_GM_neuron_glia vs. 42_bioni_neuron_glia
# -        42_GM_neuron_glia vs. 42_C001_neuron_glia
# -        42_bioni_neuron_glia vs. 42_C001_neuron_glia
# -       15_GM_neuron_glia vs. 42_GM_neuron_glia
dataset = setup_contrasts(dataset, contrast_list = list(c("42_GM_neuron", "42_bioni_neuron"),
c("42_GM_neuron", "42_C001_neuron"),
c("42_bioni_neuron", "42_C001_neuron"),
c("42_GM_neuron_glia", "42_bioni_neuron_glia"),
c("42_GM_neuron_glia", "42_C001_neuron_glia"),
c("42_bioni_neuron_glia", "42_C001_neuron_glia"),
c("15_GM_neuron_glia", "42_GM_neuron_glia")))
#
dataset = analysis_quickstart(dataset,
filter_min_detect = 2,
# filter_fraction_detect = .75,
filter_by_contrast = TRUE,
filter_topn_peptides = 0,
filter_min_peptide_per_prot = 1,
# norm_algorithm = c("vsn", "modebetween"),
norm_algorithm = "vwmb",
dea_algorithm = "ebayes",
# dea_algorithm = c("ebayes", "msempire", "msqrob"), # ebayes, msqrob, msempire, msqrobsum
dea_qvalue_threshold = 0.05,
dea_log2foldchange_threshold = 0,
output_qc_report = TRUE,
output_peptide_plots = "none",
output_abundance_tables = FALSE,
output_dir = "C:/DATA/Annemiek/data_analysis_results__subset",
output_within_timestamped_subdirectory = TRUE)
### Annemiek; subset
rm(list = ls(all.names = TRUE)) # clear everything from memory
cat("\014") # clear terminal (send the control+L character)
devtools::load_all() # load our R package
dataset = import_dataset_spectronaut("C:/DATA/Annemiek/spectronaut/20200517_182020_pilot1_ngn2_induced_neurons__subset_Report.xls", conf
dataset = update_protein_mapping_from_maxquant(dataset, path_maxquant = "C:/DATA/Annemiek/maxquant_txt", remove_shared = TRUE)
```

```
dataset = import_fasta(dataset, files = c("C:/DATA/Annemiek/UniProt_2019-11/UP000005640_9606.fasta", "C:/DATA/Annemiek/UniProt_2019-11/U
dataset = remove_proteins_by_name(dataset, remove_irt_peptides = TRUE) # optional
## create metadata file by subsetting original table
# x = openxlsx::read.xlsx("C:/DATA/Annemiek/pilot1_ngn2_induced_neurons_sample-metadata.xlsx")
# openxlsx::write.xlsx(x[x$sample_id %in% dataset$peptides$sample_id,], "C:/DATA/Annemiek/pilot1_ngn2_induced_neurons_sample-metadata__sub
dataset = import_sample_metadata(dataset, filename = "C:/DATA/Annemiek/pilot1_ngn2_induced_neurons_sample-metadata__subset.xlsx")
# -       42_GM_neuron_glia vs. 42_bioni_neuron_glia
# -        42_GM_neuron_glia vs. 42_C001_neuron_glia
# -        42_bioni_neuron_glia vs. 42_C001_neuron_glia
# -       15_GM_neuron_glia vs. 42_GM_neuron_glia
dataset = setup_contrasts(dataset, contrast_list = list(c("42_GM_neuron", "42_bioni_neuron"),
c("42_GM_neuron", "42_C001_neuron"),
c("42_bioni_neuron", "42_C001_neuron"),
c("42_GM_neuron_glia", "42_bioni_neuron_glia"),
c("42_GM_neuron_glia", "42_C001_neuron_glia"),
c("42_bioni_neuron_glia", "42_C001_neuron_glia"),
c("15_GM_neuron_glia", "42_GM_neuron_glia")))
#
dataset = analysis_quickstart(dataset,
filter_min_detect = 2,
# filter_fraction_detect = .75,
filter_by_contrast = TRUE,
filter_topn_peptides = 0,
filter_min_peptide_per_prot = 1,
# norm_algorithm = c("vsn", "modebetween"),
norm_algorithm = "vwmb",
dea_algorithm = "ebayes",
# dea_algorithm = c("ebayes", "msempire", "msqrob"), # ebayes, msqrob, msempire, msqrobsum
dea_qvalue_threshold = 0.01,
dea_log2foldchange_threshold = 0,
output_qc_report = TRUE,
output_peptide_plots = "none",
output_abundance_tables = TRUE,
output_dir = "C:/DATA/Annemiek/data_analysis_results__subset",
output_within_timestamped_subdirectory = TRUE)
### Annemiek; subset
rm(list = ls(all.names = TRUE)) # clear everything from memory
cat("\014") # clear terminal (send the control+L character)
devtools::load_all() # load our R package
dataset = import_dataset_spectronaut("C:/DATA/Annemiek/spectronaut/20200517_182020_pilot1_ngn2_induced_neurons__subset_Report.xls", confi
dataset = update_protein_mapping_from_maxquant(dataset, path_maxquant = "C:/DATA/Annemiek/maxquant_txt", remove_shared = TRUE)
dataset = import_fasta(dataset, files = c("C:/DATA/Annemiek/UniProt_2019-11/UP000005640_9606.fasta", "C:/DATA/Annemiek/UniProt_2019-11/U
dataset = remove_proteins_by_name(dataset, remove_irt_peptides = TRUE) # optional
## create metadata file by subsetting original table
# x = openxlsx::read.xlsx("C:/DATA/Annemiek/pilot1_ngn2_induced_neurons_sample-metadata.xlsx")
# openxlsx::write.xlsx(x[x$sample_id %in% dataset$peptides$sample_id,], "C:/DATA/Annemiek/pilot1_ngn2_induced_neurons_sample-metadata__sub
dataset = import_sample_metadata(dataset, filename = "C:/DATA/Annemiek/pilot1_ngn2_induced_neurons_sample-metadata__subset.xlsx")
# -       42_GM_neuron_glia vs. 42_bioni_neuron_glia
# -        42_GM_neuron_glia vs. 42_C001_neuron_glia
# -        42_bioni_neuron_glia vs. 42_C001_neuron_glia
# -       15_GM_neuron_glia vs. 42_GM_neuron_glia
dataset = setup_contrasts(dataset, contrast_list = list(c("42_GM_neuron", "42_bioni_neuron"),
c("42_GM_neuron", "42_C001_neuron"),
c("42_bioni_neuron", "42_C001_neuron"),
c("42_GM_neuron_glia", "42_bioni_neuron_glia"),
c("42_GM_neuron_glia", "42_C001_neuron_glia"),
c("42_bioni_neuron_glia", "42_C001_neuron_glia"),
c("15_GM_neuron_glia", "42_GM_neuron_glia")))
#
dataset = analysis_quickstart(dataset,
filter_min_detect = 2,
# filter_fraction_detect = .75,
filter_by_contrast = TRUE,
filter_topn_peptides = 0,
filter_min_peptide_per_prot = 1,
# norm_algorithm = c("vsn", "modebetween"),
norm_algorithm = "vwmb",
dea_algorithm = "ebayes",
# dea_algorithm = c("ebayes", "msempire", "msqrob"), # ebayes, msqrob, msempire, msqrobsum
dea_qvalue_threshold = 0.01,
dea_log2foldchange_threshold = 0,
output_qc_report = TRUE,
output_peptide_plots = "none",
output_abundance_tables = TRUE,
output_dir = "C:/DATA/Annemiek/data_analysis_results__subset",
```

```
output_within_timestamped_subdirectory = TRUE)
rm(list = ls(all.names = TRUE)) # clear everything from memory
cat("\014") # clear terminal (send the control+L character)
devtools::load_all() # load our R package
dataset = import_dataset_spectronaut("C:/DATA/Annemiek/spectronaut/20200517_182020_pilot1_ngn2_induced_neurons__subset_Report.xls", confi
dataset = update_protein_mapping_from_maxquant(dataset, path_maxquant = "C:/DATA/Annemiek/maxquant_txt", remove_shared = TRUE)
dataset = import_fasta(dataset, files = c("C:/DATA/Annemiek/UniProt_2019-11/UP000005640_9606.fasta", "C:/DATA/Annemiek/UniProt_2019-11/UI
dataset = remove_proteins_by_name(dataset, remove_irt_peptides = TRUE) # optional
append_log("******  EXPERIMENTAL DATA FILTER", type = "warning")
nrow(dataset$peptides)
dataset$peptides = dataset$peptides %>% filter(is.finite(confidence) & confidence < 0.05)
nrow(dataset$peptides)
dataset = import_sample_metadata(dataset, filename = "C:/DATA/Annemiek/pilot1_ngn2_induced_neurons_sample-metadata__subset.xlsx")
# -       42_GM_neuron_glia vs. 42_bioni_neuron_glia
# -        42_GM_neuron_glia vs. 42_C001_neuron_glia
# -        42_bioni_neuron_glia vs. 42_C001_neuron_glia
# -       15_GM_neuron_glia vs. 42_GM_neuron_glia
dataset = setup_contrasts(dataset, contrast_list = list(c("42_GM_neuron", "42_bioni_neuron"),
c("42_GM_neuron", "42_C001_neuron"),
c("42_bioni_neuron", "42_C001_neuron"),
c("42_GM_neuron_glia", "42_bioni_neuron_glia"),
c("42_GM_neuron_glia", "42_C001_neuron_glia"),
c("42_bioni_neuron_glia", "42_C001_neuron_glia"),
c("15_GM_neuron_glia", "42_GM_neuron_glia")))
#
dataset = analysis_quickstart(dataset,
filter_min_detect = 2,
# filter_fraction_detect = .75,
filter_by_contrast = TRUE,
filter_topn_peptides = 0,
filter_min_peptide_per_prot = 1,
# norm_algorithm = c("vsn", "modebetween"),
norm_algorithm = "vwmb",
dea_algorithm = "ebayes",
# dea_algorithm = c("ebayes", "msempire", "msqrob"), # ebayes, msqrob, msempire, msqrobsum
dea_qvalue_threshold = 0.01,
dea_log2foldchange_threshold = 0,
output_qc_report = TRUE,
output_peptide_plots = "none",
output_abundance_tables = TRUE,
output_dir = "C:/DATA/Annemiek/data_analysis_results__subset",
output_within_timestamped_subdirectory = TRUE)
rm(list = ls(all.names = TRUE)) # clear everything from memory
cat("\014") # clear terminal (send the control+L character)
devtools::load_all() # load our R package
dataset = import_dataset_spectronaut("C:/DATA/Annemiek/spectronaut/20200517_182020_pilot1_ngn2_induced_neurons_Report.xls", confidence_th
dataset = update_protein_mapping_from_maxquant(dataset, path_maxquant = "C:/DATA/Annemiek/maxquant_txt", remove_shared = TRUE)
dataset = import_fasta(dataset, files = c("C:/DATA/Annemiek/UniProt_2019-11/UP000005640_9606.fasta", "C:/DATA/Annemiek/UniProt_2019-11/UI
dataset = remove_proteins_by_name(dataset, remove_irt_peptides = TRUE) # optional
## create metadata file by subsetting original table
# x = openxlsx::read.xlsx("C:/DATA/Annemiek/pilot1_ngn2_induced_neurons_sample-metadata.xlsx")
# openxlsx::write.xlsx(x[x$sample_id %in% dataset$peptides$sample_id,], "C:/DATA/Annemiek/pilot1_ngn2_induced_neurons_sample-metadata__sub
rm(list = ls(all.names = TRUE)) # clear everything from memory
cat("\014") # clear terminal (send the control+L character)
devtools::load_all() # load our R package
dataset = import_dataset_spectronaut("C:/DATA/Annemiek/spectronaut/20200517_182020_ngn2_induced_neurons_Report.xls", confidence_threshold
dataset = update_protein_mapping_from_maxquant(dataset, path_maxquant = "C:/DATA/Annemiek/maxquant_txt", remove_shared = TRUE)
dataset = import_fasta(dataset, files = c("C:/DATA/Annemiek/UniProt_2019-11/UP000005640_9606.fasta", "C:/DATA/Annemiek/UniProt_2019-11/UI
dataset = remove_proteins_by_name(dataset, remove_irt_peptides = TRUE) # optional
## create metadata file by subsetting original table
# x = openxlsx::read.xlsx("C:/DATA/Annemiek/pilot1_ngn2_induced_neurons_sample-metadata.xlsx")
# openxlsx::write.xlsx(x[x$sample_id %in% dataset$peptides$sample_id,], "C:/DATA/Annemiek/pilot1_ngn2_induced_neurons_sample-metadata__sub
dataset = import_sample_metadata(dataset, filename = "C:/DATA/Annemiek/ngn2_induced_neurons_sample-metadata.xlsx")
dataset = setup_contrasts(dataset, contrast_list = list(c("42_GM_neuron", "42_bioni_neuron"),
c("42_GM_neuron", "42_C001_neuron"),
c("42_bioni_neuron", "42_C001_neuron"),
c("42_GM_neuron_glia", "42_bioni_neuron_glia"),
c("42_GM_neuron_glia", "42_C001_neuron_glia"),
c("42_bioni_neuron_glia", "42_C001_neuron_glia"),
c("15_GM_neuron_glia", "42_GM_neuron_glia")))
#
dataset = analysis_quickstart(dataset,
filter_min_detect = 2,
# filter_fraction_detect = .75,
filter_by_contrast = TRUE,
```

```
filter_topn_peptides = 0,
filter_min_peptide_per_prot = 1,
# norm_algorithm = c("vsn", "modebetween"),
norm_algorithm = "vwmb",
dea_algorithm = "ebayes",
# dea_algorithm = c("ebayes", "msempire", "msqrob"), # ebayes, msqrob, msempire, msqrobsum
dea_qvalue_threshold = 0.01,
dea_log2foldchange_threshold = 0,
output_qc_report = TRUE,
output_peptide_plots = "none",
output_abundance_tables = TRUE,
output_dir = "C:/DATA/Annemiek/data_analysis_results",
output_within_timestamped_subdirectory = TRUE)
dataset = import_sample_metadata(dataset, filename = "C:/DATA/Annemiek/ngn2_induced_neurons_sample-metadata.xlsx")
dataset = analysis_quickstart(dataset,
filter_min_detect = 2,
# filter_fraction_detect = .75,
filter_by_contrast = TRUE,
filter_topn_peptides = 0,
filter_min_peptide_per_prot = 1,
# norm_algorithm = c("vsn", "modebetween"),
norm_algorithm = "vwmb",
dea_algorithm = "ebayes",
# dea_algorithm = c("ebayes", "msempire", "msqrob"), # ebayes, msqrob, msempire, msqrobsum
dea_qvalue_threshold = 0.01,
dea_log2foldchange_threshold = 0,
output_qc_report = F,
output_peptide_plots = "none",
output_abundance_tables = TRUE,
output_dir = "C:/DATA/Annemiek/data_analysis_results",
output_within_timestamped_subdirectory = TRUE)
devtools::load_all() # load our R package
dataset = analysis_quickstart(dataset,
filter_min_detect = 2,
# filter_fraction_detect = .75,
filter_by_contrast = TRUE,
filter_topn_peptides = 0,
filter_min_peptide_per_prot = 1,
# norm_algorithm = c("vsn", "modebetween"),
norm_algorithm = "vwmb",
dea_algorithm = "ebayes",
# dea_algorithm = c("ebayes", "msempire", "msqrob"), # ebayes, msqrob, msempire, msqrobsum
dea_qvalue_threshold = 0.01,
dea_log2foldchange_threshold = 0,
output_qc_report = F,
output_peptide_plots = "none",
output_abundance_tables = TRUE,
output_dir = "C:/DATA/Annemiek/data_analysis_results",
output_within_timestamped_subdirectory = TRUE)
### Annemiek; subset
rm(list = ls(all.names = TRUE)) # clear everything from memory
cat("\014") # clear terminal (send the control+L character)
devtools::load_all() # load our R package
dataset = import_dataset_spectronaut("C:/DATA/Annemiek/spectronaut/20200517_182020_ngn2_induced_neurons_Report.xls", confidence_threshol
dataset = update_protein_mapping_from_maxquant(dataset, path_maxquant = "C:/DATA/Annemiek/maxquant_txt", remove_shared = TRUE)
dataset = import_fasta(dataset, files = c("C:/DATA/Annemiek/UniProt_2019-11/UP000005640_9606.fasta", "C:/DATA/Annemiek/UniProt_2019-11/U
dataset = remove_proteins_by_name(dataset, remove_irt_peptides = TRUE) # optional
## create metadata file by subsetting original table
# x = openxlsx::read.xlsx("C:/DATA/Annemiek/pilot1_ngn2_induced_neurons_sample-metadata.xlsx")
# openxlsx::write.xlsx(x[x$sample_id %in% dataset$peptides$sample_id,], "C:/DATA/Annemiek/pilot1_ngn2_induced_neurons_sample-metadata__sub
dataset = import_sample_metadata(dataset, filename = "C:/DATA/Annemiek/ngn2_induced_neurons_sample-metadata.xlsx")
dataset = setup_contrasts(dataset, contrast_list = list(c("42_GM_neuron", "42_bioni_neuron"),
c("42_GM_neuron", "42_C001_neuron"),
c("42_bioni_neuron", "42_C001_neuron"),
c("42_GM_neuron_glia", "42_bioni_neuron_glia"),
c("42_GM_neuron_glia", "42_C001_neuron_glia"),
c("42_bioni_neuron_glia", "42_C001_neuron_glia"),
c("15_GM_neuron_glia", "42_GM_neuron_glia")))
#
dataset = analysis_quickstart(dataset,
filter_min_detect = 2,
# filter_fraction_detect = .75,
filter_by_contrast = TRUE,
filter_topn_peptides = 0,
filter_min_peptide_per_prot = 1,
```

```
# norm_algorithm = c("vsn", "modebetween"),
norm_algorithm = "vwmb",
dea_algorithm = "ebayes",
# dea_algorithm = c("ebayes", "msempire", "msqrob"), # ebayes, msqrob, msempire, msqrobsum
dea_qvalue_threshold = 0.01,
dea_log2foldchange_threshold = 0,
output_qc_report = F,
output_peptide_plots = "none",
output_abundance_tables = TRUE,
output_dir = "C:/DATA/Annemiek/data_analysis_results",
output_within_timestamped_subdirectory = TRUE)
eset_peptides = tibble_as_eset(dataset$peptides %>% filter(is.finite(intensity) & detect == TRUE),
dataset$proteins,
dataset$samples)
MSnbase::exprs(eset_peptides) = normalize_matrix(MSnbase::exprs(eset_peptides), algorithm = norm_algorithm, mask_sample_groups = MSnbase
eset_proteins = eset_from_peptides_to_proteins(eset_peptides)
norm_algorithm="vwmb"
MSnbase::exprs(eset_peptides) = normalize_matrix(MSnbase::exprs(eset_peptides), algorithm = norm_algorithm, mask_sample_groups = MSnbase
eset_proteins = eset_from_peptides_to_proteins(eset_peptides)
### Annemiek; subset
rm(list = ls(all.names = TRUE)) # clear everything from memory
cat("\014") # clear terminal (send the control+L character)
devtools::load_all() # load our R package
dataset = import_dataset_spectronaut("C:/DATA/Annemiek/spectronaut/20200517_182020_ngn2_induced_neurons_Report.xls", confidence_threshol
dataset = update_protein_mapping_from_maxquant(dataset, path_maxquant = "C:/DATA/Annemiek/maxquant_txt", remove_shared = TRUE)
dataset = import_fasta(dataset, files = c("C:/DATA/Annemiek/UniProt_2019-11/UP000005640_9606.fasta", "C:/DATA/Annemiek/UniProt_2019-11/U
dataset = remove_proteins_by_name(dataset, remove_irt_peptides = TRUE) # optional
## create metadata file by subsetting original table
# x = openxlsx::read.xlsx("C:/DATA/Annemiek/pilot1_ngn2_induced_neurons_sample-metadata.xlsx")
# openxlsx::write.xlsx(x[x$sample_id %in% dataset$peptides$sample_id,], "C:/DATA/Annemiek/pilot1_ngn2_induced_neurons_sample-metadata__sub
dataset = import_sample_metadata(dataset, filename = "C:/DATA/Annemiek/ngn2_induced_neurons_sample-metadata.xlsx")
dataset = setup_contrasts(dataset, contrast_list = list(c("42_GM_neuron", "42_bioni_neuron"),
c("42_GM_neuron", "42_C001_neuron"),
c("42_bioni_neuron", "42_C001_neuron"),
c("42_GM_neuron_glia", "42_bioni_neuron_glia"),
c("42_GM_neuron_glia", "42_C001_neuron_glia"),
c("42_bioni_neuron_glia", "42_C001_neuron_glia"),
c("15_GM_neuron_glia", "42_GM_neuron_glia")))
#
dataset = analysis_quickstart(dataset,
filter_min_detect = 2,
# filter_fraction_detect = .75,
filter_by_contrast = TRUE,
filter_topn_peptides = 0,
filter_min_peptide_per_prot = 1,
# norm_algorithm = c("vsn", "modebetween"),
norm_algorithm = "vwmb",
dea_algorithm = "ebayes",
# dea_algorithm = c("ebayes", "msempire", "msqrob"), # ebayes, msqrob, msempire, msqrobsum
dea_qvalue_threshold = 0.01,
dea_log2foldchange_threshold = 0,
output_qc_report = F,
output_peptide_plots = "none",
output_abundance_tables = TRUE,
output_dir = "C:/DATA/Annemiek/data_analysis_results",
output_within_timestamped_subdirectory = TRUE)
nchar("protein_intensity_only_where_identified")
nchar("protein_intensity_where_identified")
nchar("protein_intensity_only_identified")
nchar("protein_intensity_only_identify")
### Annemiek; subset
rm(list = ls(all.names = TRUE)) # clear everything from memory
cat("\014") # clear terminal (send the control+L character)
devtools::load_all() # load our R package
dataset = import_dataset_spectronaut("C:/DATA/Annemiek/spectronaut/20200517_182020_ngn2_induced_neurons_Report.xls", confidence_threshol
dataset = update_protein_mapping_from_maxquant(dataset, path_maxquant = "C:/DATA/Annemiek/maxquant_txt", remove_shared = TRUE)
dataset = import_fasta(dataset, files = c("C:/DATA/Annemiek/UniProt_2019-11/UP000005640_9606.fasta", "C:/DATA/Annemiek/UniProt_2019-11/U
dataset = remove_proteins_by_name(dataset, remove_irt_peptides = TRUE) # optional
## create metadata file by subsetting original table
# x = openxlsx::read.xlsx("C:/DATA/Annemiek/pilot1_ngn2_induced_neurons_sample-metadata.xlsx")
# openxlsx::write.xlsx(x[x$sample_id %in% dataset$peptides$sample_id,], "C:/DATA/Annemiek/pilot1_ngn2_induced_neurons_sample-metadata__sub
dataset = import_sample_metadata(dataset, filename = "C:/DATA/Annemiek/ngn2_induced_neurons_sample-metadata.xlsx")
dataset = setup_contrasts(dataset, contrast_list = list(c("42_GM_neuron", "42_bioni_neuron"),
c("42_GM_neuron", "42_C001_neuron"),
c("42_bioni_neuron", "42_C001_neuron"),
```

```r
c("42_GM_neuron_glia", "42_bioni_neuron_glia"),
c("42_GM_neuron_glia", "42_C001_neuron_glia"),
c("42_bioni_neuron_glia", "42_C001_neuron_glia"),
c("15_GM_neuron_glia", "42_GM_neuron_glia")))
#
dataset = analysis_quickstart(dataset,
filter_min_detect = 2,
# filter_fraction_detect = .75,
filter_by_contrast = TRUE,
filter_topn_peptides = 0,
filter_min_peptide_per_prot = 1,
# norm_algorithm = c("vsn", "modebetween"),
norm_algorithm = "vwmb",
dea_algorithm = "ebayes",
# dea_algorithm = c("ebayes", "msempire", "msqrob"), # ebayes, msqrob, msempire, msqrobsum
dea_qvalue_threshold = 0.01,
dea_log2foldchange_threshold = 0,
output_qc_report = F,
output_peptide_plots = "none",
output_abundance_tables = TRUE,
output_dir = "C:/DATA/Annemiek/data_analysis_results",
output_within_timestamped_subdirectory = TRUE)
### Annemiek; subset
rm(list = ls(all.names = TRUE)) # clear everything from memory
cat("\014") # clear terminal (send the control+L character)
devtools::load_all() # load our R package
dataset = import_dataset_spectronaut("C:/DATA/Annemiek/spectronaut/20200517_182020_ngn2_induced_neurons_Report.xls", confidence_threshol
dataset = update_protein_mapping_from_maxquant(dataset, path_maxquant = "C:/DATA/Annemiek/maxquant_txt", remove_shared = TRUE)
dataset = import_fasta(dataset, files = c("C:/DATA/Annemiek/UniProt_2019-11/UP000005640_9606.fasta", "C:/DATA/Annemiek/UniProt_2019-11/U
dataset = remove_proteins_by_name(dataset, remove_irt_peptides = TRUE) # optional
## create metadata file by subsetting original table
# x = openxlsx::read.xlsx("C:/DATA/Annemiek/pilot1_ngn2_induced_neurons_sample-metadata.xlsx")
# openxlsx::write.xlsx(x[x$sample_id %in% dataset$peptides$sample_id,], "C:/DATA/Annemiek/pilot1_ngn2_induced_neurons_sample-metadata__sub
dataset = import_sample_metadata(dataset, filename = "C:/DATA/Annemiek/ngn2_induced_neurons_sample-metadata.xlsx")
# dataset = setup_contrasts(dataset, contrast_list = list(c("42_GM_neuron", "42_bioni_neuron"),
#                                                         c("42_GM_neuron", "42_C001_neuron"),
#                                                         c("42_bioni_neuron", "42_C001_neuron"),
#
#                                                         c("42_GM_neuron_glia", "42_bioni_neuron_glia"),
#                                                         c("42_GM_neuron_glia", "42_C001_neuron_glia"),
#                                                         c("42_bioni_neuron_glia", "42_C001_neuron_glia"),
#
#                                                         c("15_GM_neuron_glia", "42_GM_neuron_glia")))
#
dataset = analysis_quickstart(dataset,
filter_min_detect = 2,
# filter_fraction_detect = .75,
filter_by_contrast = TRUE,
filter_topn_peptides = 0,
filter_min_peptide_per_prot = 1,
# norm_algorithm = c("vsn", "modebetween"),
norm_algorithm = "vwmb",
dea_algorithm = "ebayes",
# dea_algorithm = c("ebayes", "msempire", "msqrob"), # ebayes, msqrob, msempire, msqrobsum
dea_qvalue_threshold = 0.01,
dea_log2foldchange_threshold = 0,
output_qc_report = F,
output_peptide_plots = "none",
output_abundance_tables = TRUE,
output_dir = "C:/DATA/Annemiek/data_analysis_results",
output_within_timestamped_subdirectory = TRUE)
### Annemiek; subset
rm(list = ls(all.names = TRUE)) # clear everything from memory
cat("\014") # clear terminal (send the control+L character)
devtools::load_all() # load our R package
dataset = import_dataset_spectronaut("C:/DATA/Annemiek/spectronaut/20200517_182020_ngn2_induced_neurons_Report.xls", confidence_threshol
dataset = update_protein_mapping_from_maxquant(dataset, path_maxquant = "C:/DATA/Annemiek/maxquant_txt", remove_shared = TRUE)
dataset = import_fasta(dataset, files = c("C:/DATA/Annemiek/UniProt_2019-11/UP000005640_9606.fasta", "C:/DATA/Annemiek/UniProt_2019-11/U
dataset = remove_proteins_by_name(dataset, remove_irt_peptides = TRUE) # optional
## create metadata file by subsetting original table
# x = openxlsx::read.xlsx("C:/DATA/Annemiek/pilot1_ngn2_induced_neurons_sample-metadata.xlsx")
# openxlsx::write.xlsx(x[x$sample_id %in% dataset$peptides$sample_id,], "C:/DATA/Annemiek/pilot1_ngn2_induced_neurons_sample-metadata__sub
dataset = import_sample_metadata(dataset, filename = "C:/DATA/Annemiek/ngn2_induced_neurons_sample-metadata.xlsx")
dataset = setup_contrasts(dataset, contrast_list = list(c("42_GM_neuron", "42_bioni_neuron"),
c("42_GM_neuron", "42_C001_neuron"),
```

```
c("42_bioni_neuron", "42_C001_neuron"),
c("42_GM_neuron_glia", "42_bioni_neuron_glia"),
c("42_GM_neuron_glia", "42_C001_neuron_glia"),
c("42_bioni_neuron_glia", "42_C001_neuron_glia"),
c("15_GM_neuron_glia", "42_GM_neuron_glia")))
#
dataset = analysis_quickstart(dataset,
filter_min_detect = 2,
# filter_fraction_detect = .75,
filter_by_contrast = TRUE,
filter_topn_peptides = 0,
filter_min_peptide_per_prot = 1,
# norm_algorithm = c("vsn", "modebetween"),
norm_algorithm = "vwmb",
dea_algorithm = "ebayes",
# dea_algorithm = c("ebayes", "msempire", "msqrob"), # ebayes, msqrob, msempire, msqrobsum
dea_qvalue_threshold = 0.01,
dea_log2foldchange_threshold = 0,
output_qc_report = TRUE,
output_peptide_plots = "none",
output_abundance_tables = TRUE,
output_dir = "C:/DATA/Annemiek/data_analysis_results",
output_within_timestamped_subdirectory = TRUE)
### Mandy
rm(list = ls(all.names = TRUE)) # clear everything from memory
cat("\014") # clear terminal (send the control+L character)
devtools::load_all() # load our R package
dataset = import_dataset_spectronaut(filename="C:/DATA/Mandy__Joseph_Dougherty/20200401_101418_Mark_Joseph_Report.xls", confidence_thresh
dataset = update_protein_mapping_from_maxquant(dataset, path_maxquant = "C:/DATA/remco_speclib_20190301_HC_P2_Full mouse proteome 201804'
dataset = import_fasta(dataset, files = c("C:/DATA/Remco P2 lib 2018_04 fasta/UP000000589_10090.fasta","C:/DATA/Remco P2 lib 2018_04 fast
# write_template_for_sample_metadata(dataset, "C:/temp/samples", T)
dataset = remove_proteins_by_name(dataset, remove_irt_peptides = TRUE) # optional
dataset = import_sample_metadata(dataset, filename = "C:/DATA/Mandy__Joseph_Dougherty/samples.xlsx")
dataset = setup_contrasts(dataset, contrast_list = list(c("A", "B")))
dataset = analysis_quickstart(dataset,
filter_min_detect = 5,
filter_fraction_detect = 0.75,
filter_by_contrast = FALSE,
filter_topn_peptides = 0,
filter_min_peptide_per_prot = 1,
norm_algorithm = "vwmb",
# dea_algorithm = "ebayes",
dea_algorithm = c("ebayes", "msempire", "msqrob"), # ebayes, msqrob, msempire, msqrobsum
dea_qvalue_threshold = 0.01,
dea_log2foldchange_threshold = NA,
output_qc_report = TRUE,
plot_pca_label_by_shortname = TRUE,
output_peptide_plots = "none",
output_abundance_tables = FALSE,
output_dir = "C:/DATA/Mandy__Joseph_Dougherty/data_analysis_results",
output_within_timestamped_subdirectory = TRUE)
print_dataset_summary(dataset)
### Mandy
rm(list = ls(all.names = TRUE)) # clear everything from memory
cat("\014") # clear terminal (send the control+L character)
devtools::load_all() # load our R package
dataset = import_dataset_spectronaut(filename="C:/DATA/Mandy__Joseph_Dougherty/20200401_101418_Mark_Joseph_Report.xls", confidence_thresh
dataset = update_protein_mapping_from_maxquant(dataset, path_maxquant = "C:/DATA/remco_speclib_20190301_HC_P2_Full mouse proteome 201804'
dataset = import_fasta(dataset, files = c("C:/DATA/Remco P2 lib 2018_04 fasta/UP000000589_10090.fasta","C:/DATA/Remco P2 lib 2018_04 fast
# write_template_for_sample_metadata(dataset, "C:/temp/samples", T)
dataset = remove_proteins_by_name(dataset, remove_irt_peptides = TRUE) # optional
dataset = import_sample_metadata(dataset, filename = "C:/DATA/Mandy__Joseph_Dougherty/samples.xlsx")
dataset = setup_contrasts(dataset, contrast_list = list(c("A", "B")))
dataset = analysis_quickstart(dataset,
filter_min_detect = 5,
filter_fraction_detect = 0.75,
filter_by_contrast = FALSE,
filter_topn_peptides = 0,
filter_min_peptide_per_prot = 1,
norm_algorithm = "vwmb",
# dea_algorithm = "ebayes",
dea_algorithm = c("ebayes", "msempire", "msqrob"), # ebayes, msqrob, msempire, msqrobsum
dea_qvalue_threshold = 0.01,
dea_log2foldchange_threshold = NA,
output_qc_report = TRUE,
```

```
plot_pca_label_by_shortname = TRUE,
output_peptide_plots = "none",
output_abundance_tables = FALSE,
output_dir = "C:/DATA/Mandy__Joseph_Dougherty/data_analysis_results",
output_within_timestamped_subdirectory = TRUE)
```

*Could not pretty-print your R code. Perhaps there are syntax errors in your R history? If so, either clear your R history or restart RStudio to amend.*

### 6 R session info

The computer system and versioning of all R packages used to run this analysis are shown below to facilitate, in combination with the previous section, reproducibility.

Table 2: System

| setting | value |
|---|---|
| version | R version 3.6.1 (2019-07-05) |
| os | Windows 10 x64 |
| system | x86_64, mingw32 |
| ui | RStudio |
| language | (EN) |
| collate | English_United States.1252 |
| ctype | English_United States.1252 |
| tz | Europe/Berlin |
| date | 2020-05-18 |

Table 3: Attached packages

| package | loadedversion | source |
|---|---|---|
| dplyr | 0.8.4 | CRAN (R 3.6.2) |
| ggplot2 | 3.2.1 | CRAN (R 3.6.1) |
| msdap | 0.1.7.2 | NA |
| rlang | 0.4.4 | CRAN (R 3.6.2) |
| testthat | 2.3.1 | CRAN (R 3.6.2) |
| tibble | 2.1.3 | CRAN (R 3.6.1) |
| tidyr | 1.0.2 | CRAN (R 3.6.2) |

Table 4: Packages that are not attached

| package | loadedversion | source |
|---|---|---|
| affy | 1.64.0 | Bioconductor |
| affyio | 1.56.0 | Bioconductor |
| askpass | 1.1 | CRAN (R 3.6.1) |
| assertthat | 0.2.1 | CRAN (R 3.6.1) |
| backports | 1.1.5 | CRAN (R 3.6.1) |
| Biobase | 2.46.0 | Bioconductor |
| BiocGenerics | 0.32.0 | Bioconductor |
| BiocManager | 1.30.10 | CRAN (R 3.6.2) |
| BiocParallel | 1.20.1 | Bioconductor |
| bit | 1.1-14 | CRAN (R 3.6.0) |
| bit64 | 0.9-7 | CRAN (R 3.6.0) |
| boot | 1.3-24 | CRAN (R 3.6.2) |
| callr | 3.4.2 | CRAN (R 3.6.2) |
| cli | 2.0.1 | CRAN (R 3.6.2) |
| codetools | 0.2-16 | CRAN (R 3.6.1) |
| colorspace | 1.4-1 | CRAN (R 3.6.1) |
| cowplot | 1.0.0 | CRAN (R 3.6.1) |
| crayon | 1.3.4 | CRAN (R 3.6.1) |

| package | loadedversion | source |
|---|---|---|
| data.table | 1.12.8 | CRAN (R 3.6.2) |
| desc | 1.2.0 | CRAN (R 3.6.1) |
| devtools | 2.2.2 | CRAN (R 3.6.2) |
| digest | 0.6.25 | CRAN (R 3.6.1) |
| doParallel | 1.0.15 | CRAN (R 3.6.1) |
| ellipsis | 0.3.0 | CRAN (R 3.6.1) |
| evaluate | 0.14 | CRAN (R 3.6.1) |
| fansi | 0.4.1 | CRAN (R 3.6.2) |
| farver | 2.0.3 | CRAN (R 3.6.2) |
| forcats | 0.4.0 | CRAN (R 3.6.1) |
| foreach | 1.4.8 | CRAN (R 3.6.2) |
| formatR | 1.7 | CRAN (R 3.6.1) |
| fs | 1.3.1 | CRAN (R 3.6.1) |
| ggpubr | 0.2.5 | CRAN (R 3.6.2) |
| ggrepel | 0.8.1 | CRAN (R 3.6.1) |
| ggsignif | 0.6.0 | CRAN (R 3.6.1) |
| glue | 1.3.1 | CRAN (R 3.6.1) |
| gridExtra | 2.3 | CRAN (R 3.6.1) |
| gtable | 0.3.0 | CRAN (R 3.6.1) |
| gtools | 3.8.1 | CRAN (R 3.6.0) |
| highr | 0.8 | CRAN (R 3.6.1) |
| hms | 0.5.3 | CRAN (R 3.6.2) |
| htmltools | 0.4.0 | CRAN (R 3.6.1) |
| impute | 1.60.0 | Bioconductor |
| IRanges | 2.20.2 | Bioconductor |
| iterators | 1.0.12 | CRAN (R 3.6.1) |
| knitr | 1.28 | CRAN (R 3.6.2) |
| labeling | 0.3 | CRAN (R 3.6.0) |
| lattice | 0.20-40 | CRAN (R 3.6.1) |
| lazyeval | 0.2.2 | CRAN (R 3.6.1) |
| lifecycle | 0.1.0 | CRAN (R 3.6.1) |
| limma | 3.42.2 | Bioconductor |
| lme4 | 1.1-21 | CRAN (R 3.6.1) |
| magrittr | 1.5 | CRAN (R 3.6.1) |
| MALDIquant | 1.19.3 | CRAN (R 3.6.1) |
| MASS | 7.3-51.5 | CRAN (R 3.6.2) |
| Matrix | 1.2-18 | CRAN (R 3.6.2) |
| matrixStats | 0.55.0 | CRAN (R 3.6.1) |
| memoise | 1.1.0 | CRAN (R 3.6.1) |
| minqa | 1.2.4 | CRAN (R 3.6.1) |
| msEmpiRe | 0.1.0 | Github (zimmerlab/MS-EmpiRe@ae985a9) |
| MSnbase | 2.12.0 | Bioconductor |
| munsell | 0.5.0 | CRAN (R 3.6.1) |
| mzID | 1.24.0 | Bioconductor |
| mzR | 2.20.0 | Bioconductor |
| ncdf4 | 1.17 | CRAN (R 3.6.1) |
| nlme | 3.1-144 | CRAN (R 3.6.2) |
| nloptr | 1.2.1 | CRAN (R 3.6.1) |
| openxlsx | 4.1.4 | CRAN (R 3.6.2) |
| packrat | 0.5.0 | CRAN (R 3.6.3) |
| patchwork | 1.0.0 | CRAN (R 3.6.3) |

| package | loadedversion | source |
|---|---|---|
| pcaMethods | 1.78.0 | Bioconductor |
| pdftools | 2.3 | CRAN (R 3.6.1) |
| pillar | 1.4.3 | CRAN (R 3.6.2) |
| pkgbuild | 1.0.6 | CRAN (R 3.6.1) |
| pkgconfig | 2.0.3 | CRAN (R 3.6.1) |
| pkgload | 1.0.2 | CRAN (R 3.6.1) |
| plyr | 1.8.5 | CRAN (R 3.6.2) |
| preprocessCore | 1.48.0 | Bioconductor |
| prettyunits | 1.1.1 | CRAN (R 3.6.2) |
| pROC | 1.16.1 | CRAN (R 3.6.2) |
| processx | 3.4.2 | CRAN (R 3.6.2) |
| ProtGenerics | 1.18.0 | Bioconductor |
| ps | 1.3.2 | CRAN (R 3.6.2) |
| purrr | 0.3.3 | CRAN (R 3.6.1) |
| qpdf | 1.1 | CRAN (R 3.6.1) |
| R6 | 2.4.1 | CRAN (R 3.6.1) |
| RColorBrewer | 1.1-2 | CRAN (R 3.6.0) |
| Rcpp | 1.0.3 | CRAN (R 3.6.1) |
| readr | 1.3.1 | CRAN (R 3.6.1) |
| remotes | 2.1.1 | CRAN (R 3.6.2) |
| reshape2 | 1.4.3 | CRAN (R 3.6.1) |
| rmarkdown | 2.1 | CRAN (R 3.6.2) |
| rprojroot | 1.3-2 | CRAN (R 3.6.1) |
| rstudioapi | 0.11 | CRAN (R 3.6.2) |
| S4Vectors | 0.24.3 | Bioconductor |
| scales | 1.1.0 | CRAN (R 3.6.1) |
| sessioninfo | 1.1.1 | CRAN (R 3.6.1) |
| stringi | 1.4.6 | CRAN (R 3.6.2) |
| stringr | 1.4.0 | CRAN (R 3.6.1) |
| styler | 1.3.2 | CRAN (R 3.6.2) |
| tidyselect | 1.0.0 | CRAN (R 3.6.2) |
| tinytex | 0.20 | CRAN (R 3.6.2) |
| usethis | 1.5.1 | CRAN (R 3.6.3) |
| utf8 | 1.1.4 | CRAN (R 3.6.1) |
| vctrs | 0.2.3 | CRAN (R 3.6.2) |
| viridis | 0.5.1 | CRAN (R 3.6.1) |
| viridisLite | 0.3.0 | CRAN (R 3.6.1) |
| vsn | 3.54.0 | Bioconductor |
| withr | 2.1.2 | CRAN (R 3.6.1) |
| xfun | 0.12 | CRAN (R 3.6.2) |
| XML | 3.98-1.20 | CRAN (R 3.6.0) |
| xtable | 1.8-4 | CRAN (R 3.6.1) |
| yaml | 2.2.1 | CRAN (R 3.6.2) |
| zip | 2.0.4 | CRAN (R 3.6.1) |
| zlibbioc | 1.32.0 | Bioconductor |
